# SMAD3-driven cell state transitions underpin immunoresistance in bladder cancer

**DOI:** 10.64898/2026.10.21.760742

**Authors:** Sarah Y.T. Robertson, Jing Yang, Neil O. Carragher, Alison Munro, Nora Rauch, Thomas Sevrin, Kieran Wynne, Walter Kolch, Oleksii S. Rukhlenko, Boris N. Kholodenko

## Abstract

Despite the clinical success of anti-PD-1/PD-L1 immunotherapy in a subset of patients with high-risk bladder cancer (BLCA), most tumors remain refractory to treatment. We hypothesized that immunotherapy resistance is encoded within the intrinsic signaling state of BLCA cells and arises through specific cell-state transitions that precede therapeutic exposure. To test this hypothesis, we applied a cell State Transition Assessment and Regulation (cSTAR) framework to identify key signaling modules and aberrant causal network connections that (i) drive oncogenic progression and acquisition of aggressive phenotypes and (ii) govern BLCA cell susceptibility to T cell-mediated cytotoxicity. cSTAR analysis identified TGFβ/SMAD3 signaling as the principal driver of both oncogenic transformation and immunoresistance in BLCA cells. Inhibiting SMAD3 sensitized BLCA cells to PD-1-expressing Jurkat cells and primary human CD4^+^ T cell-mediated cytotoxicity through suppression of autophagy and activation of multiple cell death pathways. By integrating cell state transition analysis with functional immune co-culture assays, our work demonstrates that tumor cell state transitions control sensitivity to T cell killing, complementing established immune-intrinsic mechanisms of immunotherapy response.

## INTRODUCTION

Immunotherapy has transformed the treatment landscape for bladder cancer (BLCA), yet durable clinical responses to anti-PD-1/PD-L1 therapy are achieved in only a subset of patients. Objective response rates for high-risk muscle-invasive BLCA (MIBC) typically range from 20 to 40%, irrespective of treatment setting^1–11^. This limited efficacy is particularly striking given the high tumor mutational burden of BLCA, a feature generally associated with favorable responses to immunotherapy^12,13^. Similarly, intravesical Bacillus Calmette–Guérin (BCG) therapy can induce complete responses in 50–70% of patients with non-muscle-invasive BLCA (NMIBC)^14^, yet many patients eventually relapse or develop treatment resistance. Although anti-PD-1/PD-L1 therapies provide clinical benefit in BCG-unresponsive NMIBC^10,15^ and in muscle-invasive BLCA (MIBC) failure^13,16^, most tumors remain refractory to treatment, highlighting our incomplete understanding of the mechanisms that determine immunotherapy response.

Efforts to explain variability in immunotherapy response have primarily focused on the immune compartment. High expression of PD-L1 in tumor and immune cells has been widely investigated as a biomarker of response to immunotherapy, but no single biomarker predicts BLCA patient response to immunotherapy^1,17^. Extensive work has characterized inhibitory checkpoint pathways that suppress T cell activation, promote T cell exhaustion, and limit anti-tumor immunity^1,17^. Molecular classification studies have further established BLCA subtypes, including multiple luminal subtypes, a stroma-rich subtype, a basal-squamous subtype, and a neuroendocrine-like subtype^18,19^. Collectively, current studies suggest that expression of PD-L1 and other surface molecules, genomic mutational patterns, stromal composition, and immune-cell infiltration are major determinants of immunotherapy outcome^19–24^.

Successful immunotherapy requires not only functional immune effector cells, but tumor cells must remain susceptible to immune-mediated killing. Recent studies have identified cytotoxic CD4^+^ T cells as major mediators of anti-tumor immunity in BLCA, and signatures of these cells predict response to BCG^22^ and anti-PD-L1 therapy^24^. Like CD8^+^ T cells, CD4^+^ T cells eliminate tumor cells through multiple mechanisms, including engagement of death receptors and secretion of cytotoxic mediators such as perforin, granzymes, IFNγ, and TNFα^25^. While considerable attention has been devoted to mechanisms that impair T-cell function, substantially less is known about the mechanisms that determine the intrinsic sensitivity of BLCA cells to these cytotoxic signals. We, therefore, investigated whether BLCA cells can evade immune destruction not only through checkpoint engagement^26,27^ but also by acquiring resistance to T cell cytotoxic factors^28,29^. We posit that BLCA cells dynamically regulate their susceptibility to immune attack through coordinated cell-state transitions and associated cell intrinsic signaling programs that control survival pathways, stress responses, and cellular responses to cytotoxic stimuli. Under this hypothesis, immunotherapy resistance is not solely a consequence of impaired T cell activation but also reflects tumor-cell state transitions that reduce sensitivity to immune-mediated killing.

To test this hypothesis, we applied the cell State Transition Assessment and Regulation (cSTAR) framework^30^ to identify signaling pathways that govern BLCA cell responsiveness to activated T cells and map cancer-cell state transitions associated with resistance or sensitivity to T cell cytotoxicity. We then experimentally validated the cSTAR-predicted, key pathway that controls immunosensitivity by targeting SMAD3 in co-cultures of BLCA cells with PD-1-expressing Jurkat cells and primary human CD4^+^ T cells. Our findings establish a mechanistic link between tumor-cell state regulation and CD4^+^ T cell-mediated cytotoxicity and demonstrate that immunotherapy outcomes are determined by the interplay between T cell function and dynamic tumor-cell state transitions that regulate intrinsic susceptibility to immune-mediated killing.

## RESULTS

### BASAL-LIKE BLCA CELLS ARE MORE IMMUNORESISTANT THAN NON-TYPE OR LUMINAL-LIKE CELLS

To systematically evaluate the susceptibility of BLCA cells to T cell-mediated cytotoxicity, we established an *in vitro* co-culture model pairing a diverse panel of human-derived BLCA cell lines with either the Jurkat E6.1 CD4^+^ leukemic T cell model or primary human CD4^+^ T cells. Jurkat E6.1 cells serve as a robust model for characterizing target cell engagement and lytic factor deployment, having recently been shown to be cytotoxic against lung cancer cells in an *in vitro* model of lung cancer by engaging the cancer cells and releasing cytotoxic factors such as perforin and granzymes, especially after activation^31^. Therefore, we used Jurkat E6.1 cells that express GFP under the control of an NF-κB-responsive promoter, providing a fluorescent readout of activation^32^. To model the influence of inhibitory immune checkpoints, a subset of these cells was further modified to stably express PD-1 receptor (“Jurkat-PD-1”)^32^. Jurkat-PD-1 cells consistently express CD4 and PD-1 over multiple passages (Extended Data Fig. 1a). To provide a physiological way of stimulation by cell-cell contact, murine Bw5147 cells stably expressing a membrane bound anti-CD3 antibody fragment were used (T cell stimulator, TCS). TCS were additionally engineered to express PD-L1, the ligand for PD-1 (TCS-PD-L1)^32^. Jurkat-PD-1 cells exhibited significantly increased NF-κB activation when co-cultured with TCS, and reduced NF-κB activation when co-cultured with TCS-PD-L1 cells (Extended Data Fig. 1b). Importantly, the PD-1 effects were reversed by the anti-PD-1 monoclonal antibody pembrolizumab, confirming functional PD-1 receptor fidelity in Jurkat-PD-1 cells (Extended Data Fig. 1b)^32^. No significant changes in apoptosis of Jurkat-PD-1 cells were observed in any activation condition (Extended Data Fig. 1c).

To ensure translational relevance, our cell panel spanned five BLCA lines representing distinct clinical subtypes and invasive potentials alongside the non-malignant immortalized bladder epithelial line HCV-29^33^ as a control. Specifically, the panel included the non-muscle-invasive, luminal-like RT4 line^34^; the non-type lines T24 (non-invasive) and J82 (invasive)^35,36^; and the highly invasive, basal-like lines HT1376^37^ and BC-3C^38^, classified according to consensus subtypes^39,40^.

To test whether Jurkat-PD-1 cells can kill BLCA cells, Jurkat-PD-1 cells were activated with PMA and ionomycin and co-cultured with either non-malignant or BLCA cells at a 10:1 effector-to-target ratio. BLCA cell viability was assessed via MTS assay after three days (Fig. 1a). In the non-cancerous control line HCV-29, co-culture with non-activated Jurkat-PD-1 cells reduced viability by 40%, whereas activated Jurkat cells achieved 60% inhibition (Fig. 1b). The non-invasive luminal-like RT4 line was highly sensitive to activated Jurkat-PD-1 cells, with a drastic 75% loss of viability compared with 40% in the presence of non-activated controls. The non-invasive non-type T24 line was also sensitive to activated Jurkat-PD-1 cells, with viability loss increasing from 50% with non-activated Jurkat-PD-1 cells to 75% following their activation (Fig. 1b). In contrast, the invasive non-type line J82 and the basal-like line BC-3C exhibited robust immunoresistance. J82 viability was only reduced by 40% following both non-activated and activated Jurkat-PD-1 co-culture, while BC-3C cell viability showed a maximum of 25% reduction. HT1376 exhibited even stronger immunoresistance, showing no reduction in viability following co-culture with non-activated or activated Jurkat-PD-1 cells. These data define a spectrum of immunosensitivity across the BLCA panel (Fig. 1b), with HCV-29, RT4, and T24 categorized as “immunosensitive,” and J82, BC-3C, and HT1376 as “immunoresistant.” Crucially, the cytotoxicity of both non-activated and activated Jurkat-PD-1 cells are significantly reduced by approximately 30% on HCV-29 cells and 50% on RT4 and T24 cells when a Transwell co-culture system is used. This shows that the full cytotoxic effect of Jurkat-PD-1 cells requires direct, physical cell-to-cell contact rather than paracrine signaling (Extended Data Figs. 2a and 2b).

**Figure 1.**
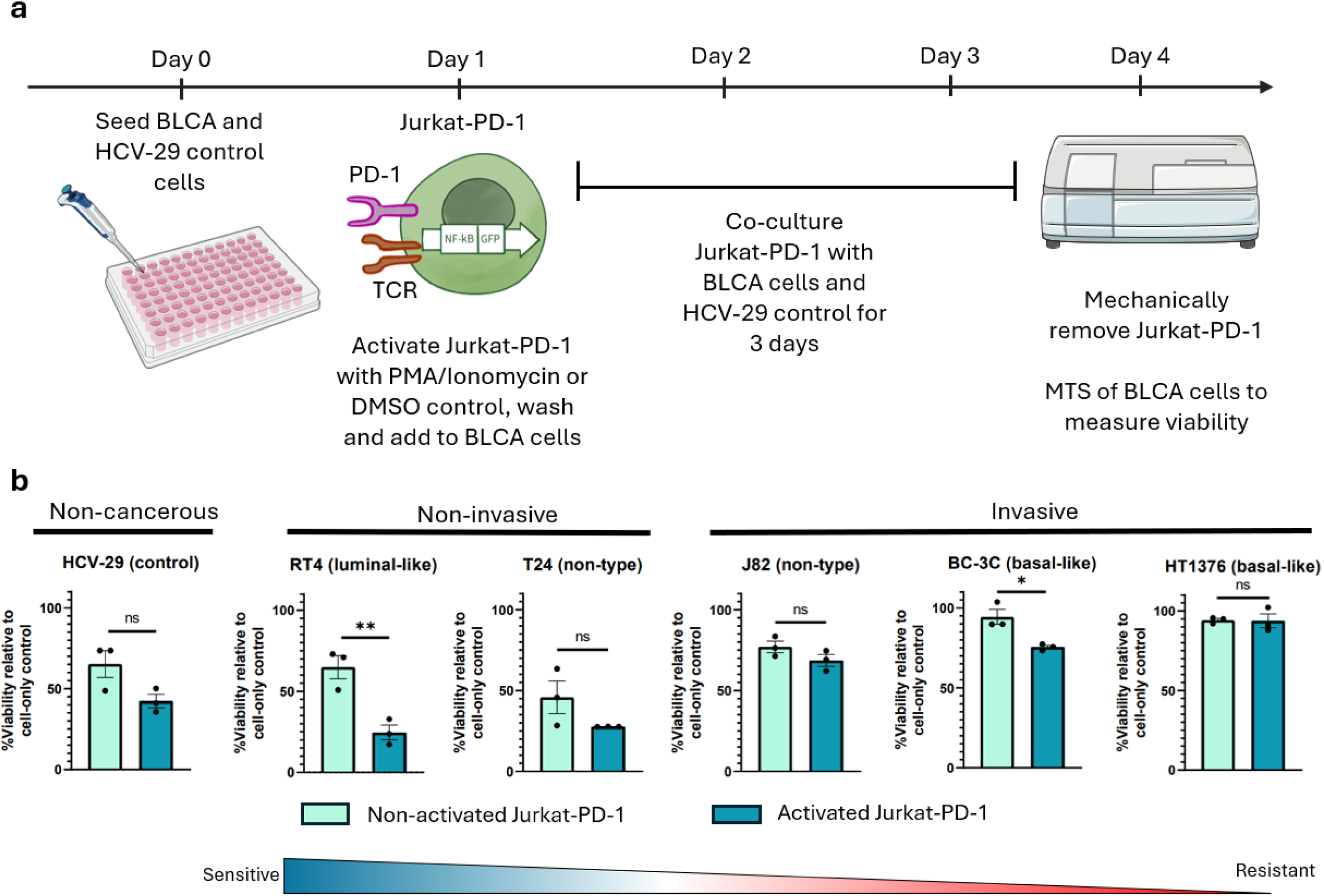
Immunoresistant and immunosensitive BLCA cells. **a.** Experimental schematic. Jurkat cells were engineered to express GFP under the control of an NF-κB reporter, with a subset further engineered to express PD-1. Jurkat-PD-1 cells were activated with PMA and Ionomycin (100 nM each) for 3 h, washed with serum-containing media to remove stimuli, and co-cultured with BLCA cells at a 10:1 ratio (Jurkat:BLCA) for 3 days as a model of CD4⁺ T cell cytotoxicity. **b.** MTS-based viability analysis of BLCA cells 72 h post-Jurkat-PD-1 co-culture. BLCA cells were ranked left-to-right sensitive to resistant based on their sensitivity to Jurkat-PD-1 cells. **\***p<0.05, **p<0.01 using two-sample t-test, n=3 independent experiments. Illustrations from NIAID NIH BioArt Source (bioart.niaid.nih.gov/bioart/408) and Servier Medical Art (https://smart.servier.com), used under CC BY 4.0 license (https://creativecommons.org/licenses/by/4.0/).

Because CD4⁺ T cells can recognize antigens via major histocompatibility complex class II (MHC-II), we assessed whether this pathway was required for Jurkat-mediated BLCA cytotoxicity. We evaluated surface expression of HLA-DR (the human MHC-II molecule) via flow cytometry, excluding dead cells with 7-AAD dye. RT4 exhibited the highest HLA-DR expression (30%), followed by J82 (18%), while HCV-29 and HT1376 showed minimal levels (<5%), and expression was undetectable in BC-3C and T24 (Extended Data Fig. 3a). To determine functional relevance, we added an MHC-II neutralizing antibody to co-cultures of Jurkat-PD-1 and immunosensitive BLCA lines. MHC-II blockade did not attenuate Jurkat-PD-1 cytotoxicity against HCV-29 or RT4 cells (Extended Data Fig. 3b), suggesting that Jurkat-PD-1-mediated killing occurs through an MHC-II-independent mechanism.

In summary, BLCA cell lines exhibit a spectrum of immunosensitivity to activated Jurkat-PD-1 cells, with immunoresistance increasing in invasive and basal-like subtypes independently of HLA-DR expression.

### INTERROGATING BLCA PATHWAYS TO IDENTIFY KEY NODES IN CELL-LINE SPECIFIC NETWORKS

To construct cell-specific core networks, we first identified the pathways that would serve as functional modules or nodes in these networks. We performed a targeted drug screen on pathways implicated in BLCA measuring the viabilities of our BLCA cells and non-malignant HCV-29 cells over 72 hours. Fig. 2a summarizes the results as a normalized Area Under the Curve (AUC) heatmap, while Extended Data Fig. 4 presents detailed viability dose-response curves. Inhibition of CDK1/2/5/9, mTOR, and GPX4 significantly reduced viability across all cell lines, including the non-malignant control, represented by low AUC. Invasive lines showed greater sensitivity to FGFR inhibition compared to non-invasive cell lines, whereas non-invasive lines were more sensitive to inhibition of IKK2, which mediates canonical NF-κB signaling. Cell lines with epithelial morphology (RT4 and HT1376) were more sensitive to MEK inhibition. Furthermore, luminal- and basal-like lines exhibited stronger sensitivity to SRC inhibition compared to non-type lines. RT4 was more sensitive to TGFβR inhibition, but more resistant to SMAD3 inhibition, than all other cell lines. All cell lines displayed profound resistance to XIAP and CDK4/6 inhibition.

**Figure 2.**
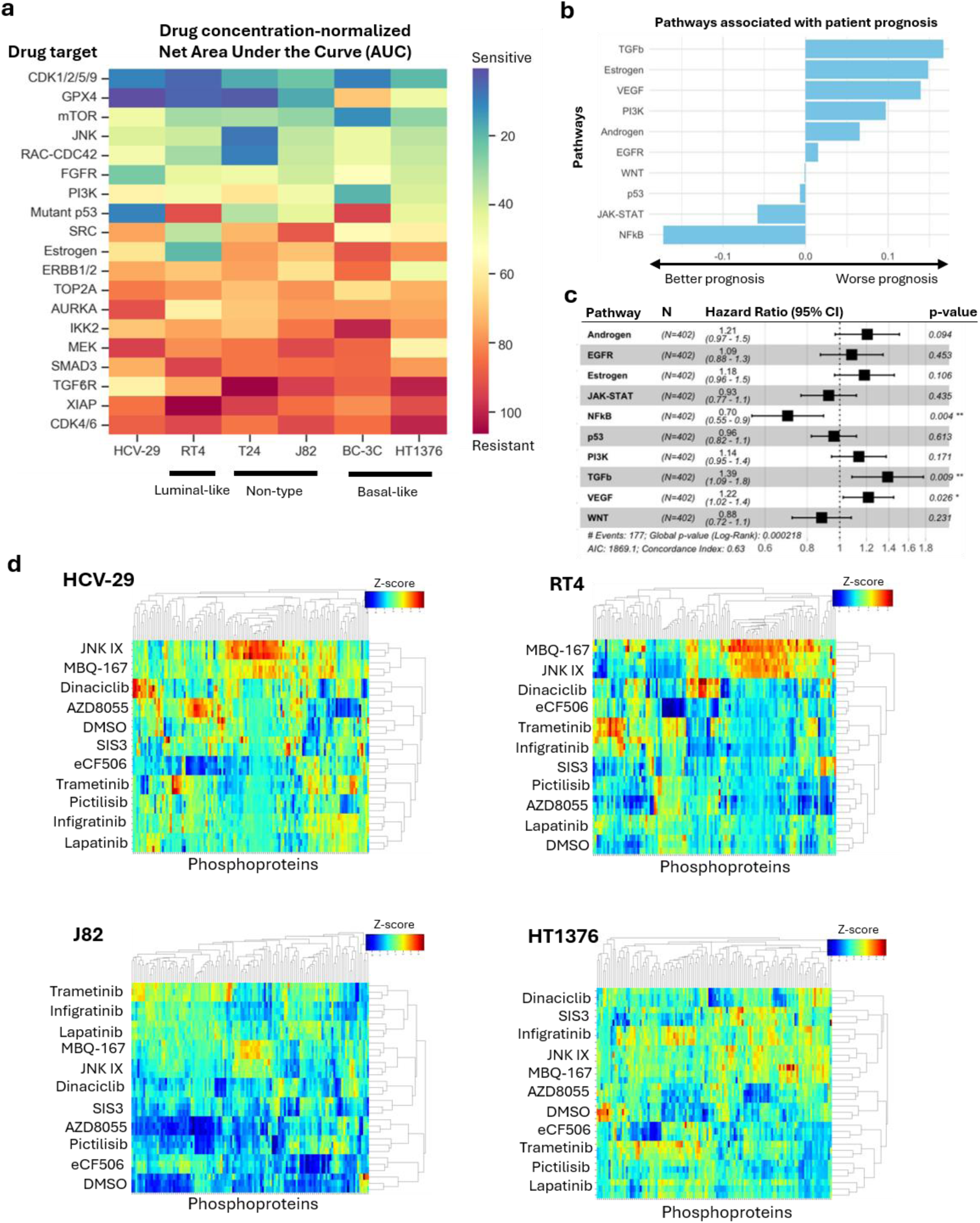
Core pathways involved in BLCA pathogenesis. **a.** Drug sensitivity profiles of BLCA cell lines treated with a range of concentrations of drugs targeting pathways implicated in BLCA pathogenesis. Area under the curve (AUC) was calculated and normalized by dividing by drug concentration, red denote drug resistance and blue denotes drug sensitivity. Drugs were ranked top to bottom based on overall cell sensitivity. Full MTS data are provided in Extended Data Fig. 4, and the full list of drugs and concentrations is provided in Extended Data Table S1. Cell lines were grouped as control non-malignant (HCV-29), luminal-like (RT4), non-type (T24 and J82), and basal-like (BC-3C and HT1376). **b.** A GLMNET model was applied to BLCA patient RNA-seq data from TCGA to identify pathways associated with clinical prognosis. Pathways with positive coefficients are associated with worse prognosis, whereas negative coefficients are associated with better prognosis. **c.** Hazard ratios were calculated by multivariate ordinary Cox regression (*p<0.05, **p<0.01). **d.** Non-malignant (HCV-29), luminal (RT4), non-type (J82), and basal (HT1376) cell lines were treated with single doses of drugs targeting selected pathways. Cellular responses were assessed by proteomics and RPPA. RPPA data (∼120 proteins, predominantly phosphoproteins) are shown as heatmaps and clustered by Z-score. n=3–4 independent experiments for all data.

In parallel, pathway activities were estimated by analyzing BLCA transcriptomic data from The Cancer Genome Atlas (TCGA)^18^ using PROGENY signatures^41^. These pathway activity scores were used as input to a regularized Cox regression model^42,43^ implemented within GLMNET R package^44^ (Methods) to identify pathways associated with patient prognosis (Fig. 2b). Subsequent multivariate Cox regression demonstrated that TGF-β and VEGF activation were significantly associated with poor prognosis, whereas NF-κB signaling was associated with favorable outcomes (Fig. 2c). Importantly, while TGFβ signaling was a strong predictor of poor prognosis, most cell lines were resistant to TGFβ receptor (TGFβR) and SMAD3 inhibitors (heat map in Fig. 2a and Extended Data Fig. 4). This suggests that the prognostic association with TGFβ and NF-κB may reflect immune-cell modulation rather than direct effects of BLCA cells.

We combined these findings with the existing literature^45^ to select 10 targetable pathways based on their impact on BLCA cell viability and clinical prognosis. These pathways were individually perturbed using specific inhibitors at doses corresponding approximately to the asymptotic floor of the viability curve in at least one cell line in our panel. We used JNK inhibitor IX (JNK), MBQ-167 (RAC-CDC42), Dinaciclib (CDK1/2/5/9), AZD8055 (mTOR), SIS3 (SMAD3), eCF506 (SRC), Trametinib (MEK), Pictilisib (PI3K),

Infigratinib (FGFR), and Lapatinib (ERBB1/2). We then generated a comprehensive library of phosphoproteomic responses in the control HCV-29 and malignant RT4, J82, and HT1376 cell lines using **R**everse-**P**hase **P**rotein **A**rray (RPPA; 120 antibodies) and mass spectrometry (MS) proteomics (Extended Data Fig. 5a). Each cell line exhibited a unique (phospho)proteomic response to these inhibitors (Fig. 2d), reflecting the profound molecular heterogeneity across BLCA states. Proteomic profiling of the cells before drug perturbation further confirmed this heterogeneity, particularly among the immunoresistant cell lines (Extended Data Fig. 5b–f).

Interestingly, JNK inhibitor IX paradoxically increased JNK pathway activity (Extended Data Fig. 6), likely through dimerization-driven allosteric effects, a mechanism analogous to the paradoxical activation previously described for RAF kinases^46,47^. This paradoxical activation does not preclude cSTAR network reconstruction, as Modular Response Analysis (MRA), a central component of the cSTAR algorithm, can be applied equally to both activating and inhibiting perturbations⁴⁷.

In summary, the drug response profiles of non-malignant and BLCA cells point to core signaling network pathways representing pathways critical to cell viability and patient survival. Furthermore, these drug responses determined the optimal dosages for subsequent perturbation experiments used in RPPA and proteomic analyses, which form the basis of the cSTAR network reconstruction^30^.

### MAPPING OF ONCOGENIC, INVASIVE, AND IMMUNORESISTANT CELL STATES BY CSTAR

#### BLCA Cell State Transition Analysis using MS Proteomic data

To characterize the molecular landscape of non-malignant and BLCA cells, we employed the cSTAR approach^30^. Using in-house generated MS proteomics data, we mapped unperturbed cell molecular states and separated them using support vector machines (SVM), a machine-learning classification method^48,49^. SVM constructs hyperplanes that maximize the separation between states in a multidimensional dataspace^30^. We then derived the State Transition Vectors (STVs) as the unit vectors normal to the separating hyperplanes. An STV defines the shortest path from one cell state to another in multidimensional omics space, which can span (phospho)proteomic, transcriptomic, or metabolomic data^30,50,51^. Based on proteomics data, we generated an aggregate oncogenic STV (STV_onc_) that represents the transition from non-malignant cells to the aggregate cancer cells’ state (Fig. 3a, Methods). Subsequently, we generated the immunoresistance-associated STV (STV_imres_), which captures the transition from the immunosensitive cell lines HCV-29 and RT4 to immunoresistant cell lines J82 and HT1376 in proteomic space (Fig. 3b).

**Figure 3.**
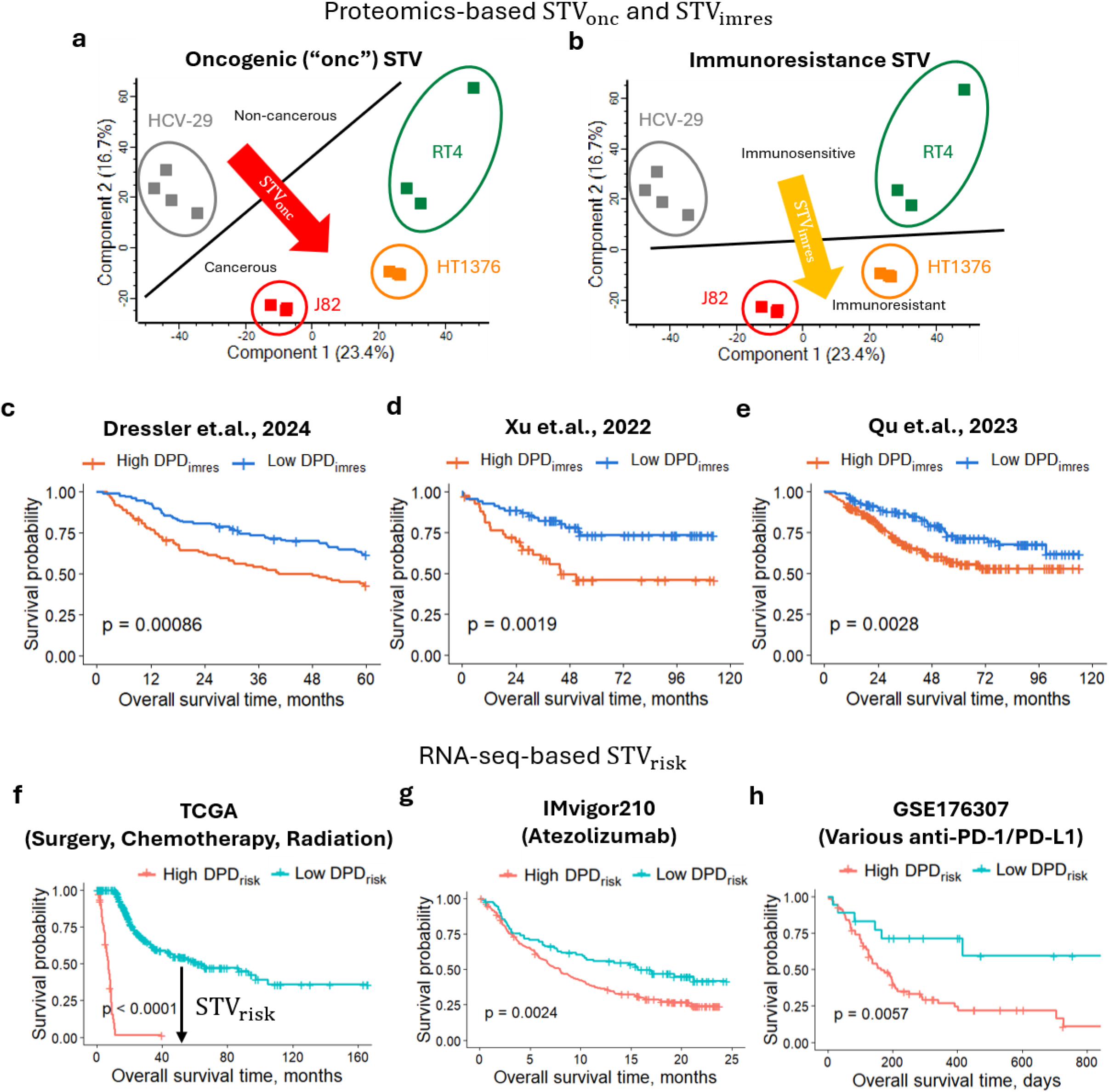
Construction and Validation of State Transition Vectors (STVs). **a-b:** Schematics illustrating the construction of STV_onc_ and STV_imres_ using proteomics data. **a.** The global oncogenic vector (STV_onc_; red arrow) is determined as the unit vector orthogonal to SVM–generated hyperplanes separating integrated proteomic profiles of RT4, HT1376, and J82 cancer cell lines from HCV-29 cells. **b.** The immunoresistance vector (STV_imres_) was constructed using the SVM-derived hyperplane separating proteomic profiles of immunosensitive cell lines (RT4, HCV-29) from immunoresistant cell lines (J82, HT1376). **c-e.** Validation of STV_imres_ using proteomics data from three independent studies of urothelial carcinoma: tumor tissue from patients with NMIBC and MIBC (c, d) and pre-operative plasma from patients with upper tract urothelial carcinoma (e). The Dynamic Phenotype Descriptor (DPD) generated from STV_imres_, DPD_imres_, significantly stratifies patients by prognosis in the Dressler et al. 2024 (**c)**, Xu et al. 2022 (**d**), and Qu et al. 2023 (**e**) studies. **f.** Construction of the STV_risk_ by integrating survival analysis and RNA-seq data from TCGA. Patients in the TCGA cohort were treated with surgery, chemotherapy, and/or radiation. Patients with favorable (cyan) and unfavourable (coral) prognoses were separated in the RNA-seq data space, which is encapsulated in the STV_risk_. The DPD_risk_ derived from STV_risk_ significantly stratifies patients by prognosis. **g-h.** DPD_risk_ was further validated using IMvigor210 trial bulk transcriptomic data from patients treated with the immune checkpoint inhibitor Atezolizumab (**g**) and from patients treated with various immune checkpoint inhibitors (GSE176307, **h**). DPD_risk_ significantly stratifies patients by prognosis as shown in the Kaplan-Meier curves.

The constructed STVs enable the quantification of the cell phenotypic features as Dynamic Phenotype Descriptor (DPD) scores^30,50,51^. The absolute value of the DPD score quantifies the distance from a proteomic or transcriptomic data point to the separating hyperplane constructed by the SVM. When drug or genetic perturbations alter this score, positive or negative DPD changes shift the cell state toward or away from a given phenotype, respectively.

We then validated the utility of STV_imres_ using survival analysis in three independent studies that included proteomics and survival data from patients with urothelial carcinoma (tumor tissue from NMIBC and MIBC patients, and plasma from patients with upper tract urothelial carcinoma) (Figs. 3c^52^, 3d^53^, 3e^54^). In all three studies, DPD_imres_ (calculated using projections of patient proteomic profiles onto STV_imres_ (Methods) significantly stratified patients into groups with favorable and unfavorable prognosis (Fig. 3c-e). Importantly, these results suggest that although STV_imres_ was developed using cell line proteomics data, it captures a protein signature predictive of overall survival of BLCA patients.

#### Transcriptomic Dataspace

To generate and validate a prognostic transcriptomic STV, which we designated STV_risk_, we integrated transcriptomic and longitudinal clinical data using regularized Cox regression (Methods). Applying this model to the TCGA bladder cancer dataset successfully stratified patients into favorable and unfavorable prognosis cohorts, as shown by Kaplan-Meier analysis using DPD_risk_ (Fig. 3f). We further validated STV_risk_ across two independent immunotherapy cohorts: the IMvigor210 trial^55^ and GSE176307^56^, where it robustly separated patients by outcome (Fig. 3g,h). Notably, the capacity of an STV derived from a non-immunotherapy baseline dataset (TCGA) to effectively predict survival in immunotherapy-treated patients suggests that immune-mediated cell death is a fundamental determinant of bladder cancer prognosis, independent of therapeutic modality. This discovery demonstrates the value added by cSTAR to the interpretation and prediction of complex biological responses.

To derive a transcriptomic signature of oncogenic BLCA transformation, we applied our framework to single-cell transcriptomics data from a semi-spontaneous mouse model of urothelial cancer (OH-BBN-exposed Hgf-Cdk4^R24C^ mice) comprising healthy bladder, non-muscle-invasive, and muscle-invasive bladder cancer^57^. We extracted data points for the urothelial cell population (Extended Data Fig. 7a, circled) and employed SVM to distinguish between healthy and malignant states in order to generate an oncogenic transformation STV (DPD^mouse^) based on mouse single cell RNA-seq (Extended Data Fig. 7b). This STV serves as an independent measure of oncogenic potential of cells, allowing us to avoid dataset-specific effects.

Table 1 defines all STVs and DPDs used to quantify the oncogenic potential, invasiveness, and immunoresistance of BLCA cells in proteomic and transcriptomic data spaces.

**Table 1:** DPD information.

| DPD | STV generated from: | Positive DPD | Negative DPD |
| --- | --- | --- | --- |
| $DPD_{apop}^{suppr}$ | RPPA data for cleaved Caspase 3 and cleaved-PARP | Suppresses apoptosis | Activates apoptosis |
| $DPD_{onc}$ | Comparison between HCV-29 (control) and all cancer cells | Promotes oncogenic transformation | Prevents oncogenic transformation |
| $DPD_{imres}$ | Comparison between HCV-29 + RT4 (immunosensitive) and J82 + HT1376 (immunoresistant) | Promotes immunoresistance | Promotes immunosensitivity |
| $DPD_{onc}^{mouse}$ | Comparison between urothelial cells from tumor-bearing (NMIBC and MIBC) and healthy bladders using scRNA-seq data of the semi-spontaneous OH-BBN-induced mouse bladder cancer model | Promotes oncogenic transformation | Prevents oncogenic transformation |

| DPD<sub>risk</sub> | Comparison between good prognosis and poor prognosis patients from TCGA RNA-seq data | Associated with poor prognosis | Associated with good prognosis |
| --- | --- | --- | --- |

Altogether, these results demonstrate that cSTAR-derived STVs provide a comprehensive framework for quantifying oncogenic potential, invasiveness, and immunoresistance across transcriptomic and proteomic dataspaces in both human and mouse models, thereby establishing a foundation for mapping the core signaling networks that govern these diverse phenotypes.

### cSTAR RECONSTRUCTION OF CELL-SPECIFIC SIGNALING NETWORKS IDENTIFIES SMAD3 AS A DRIVER OF ONCOGENICITY AND IMMUNORESISTANCE

While STVs map static cell states, uncovering the underlying regulatory logic requires dynamic, perturbation-based interrogation of the signaling circuitry. We, therefore, employed the cSTAR framework to reconstruct core regulatory networks, integrating RPPA drug perturbation data with phenotypic descriptors (DPD scores) derived from our MS proteomics data and the aforementioned transcriptomics datasets^30,50,51^. The DPDs, computed using the corresponding proteomic and transcriptomic STVs (see definitions of these phenotype descriptors in Table 1), function as additional network nodes. These nodes capture the phenotypic outcomes of specific perturbations, effectively linking the localized signaling events captured in the core network to the global molecular network driving cell fate transitions. By measuring how the (phospho)proteome shifts in response to small-molecule inhibition, the cSTAR algorithm reconstructs cell line-specific networks that quantify direct causal links between core signaling modules and phenotypic features (Methods). Reconstructing local causal connections enables the prediction of system-level (global) responses of diverse DPD-quantified phenotypic features following the activation or inhibition of core network pathways^30,58,59^.

Our RPPA analysis used both cleaved Caspase 3 and cleaved PARP as established apoptosis markers. The terminal apoptosis state corresponds to cell death, which cannot be directly characterized by omics measurements. However, cSTAR enables the use of quantitative drug-induced changes in apoptosis markers as surrogate readouts of the programmed cell death module (Methods). Using RPPA perturbation data, cSTAR reconstructs causal links from signaling modules to the programmed cell death module. For clarity, we characterize changes in this phenotypic module as the suppression of apoptosis 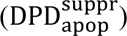, where positive values indicate suppression of apoptosis.

#### HCV-29 network reconstruction

The inferred systems-level phenotypic responses in the non-cancerous HCV-29 line suggest that CDK1/2 and JNK activities drive oncogenic transformation, as evidenced by their positive connections that increase the oncogenic DPD (DPD_onc_). Conversely, ERBB1/2, SRC, mTOR, PI3K, SMAD3, ERK, and RAC-CDC42 signaling appear to suppress oncogenicity in this context (Fig. 4a). Notably, the 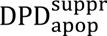 response in HCV-29 cells to JNK is anti-correlated with their oncogenic and immunoresistance DPD responses. This indicates that JNK promotes transformation and simultaneously triggers programmed cell death, potentially serving as intrinsic safeguard in HCV-29 against malignant conversion. Conversely, CDK1/2 is associated with a positive 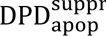, suggesting that CDK1/2 activity simultaneously promotes oncogenic transformation and prevents HCV-29 cell death.

**Figure 4.**
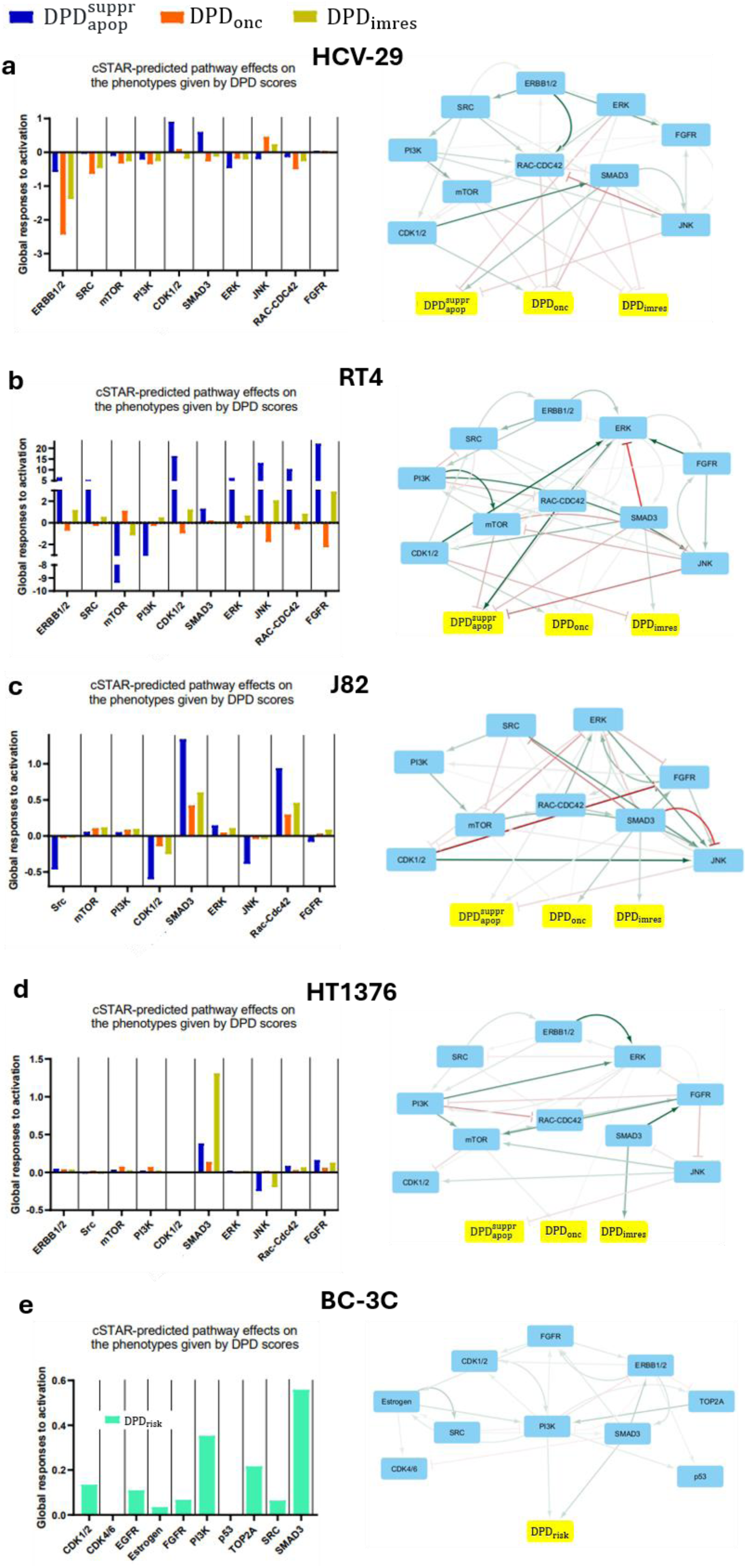
cSTAR-inferred cell-specific networks identify SMAD3 as a key driver of oncogenicity and immunoresistance. Cell-specific signaling networks were reconstructed using RPPA and mass spec proteomics data for HCV-29 (**a**), RT4 (**b**), J82 (**c**), and HT1376 cells (**d**), and RNA-seq data for BC-3C cells (**e**). In each panel, the left-hand bar graph shows the global phenotypic responses of each DPD to pathway perturbations in the corresponding cell line, while the network on the right shows the corresponding cSTAR-inferred local connections. In the network diagrams, individually perturbed core network modules are shown in blue, and the inferred DPD modules are in yellow. Green arrows indicate positive connections, while red blocked arrows show negative connections from one node to another. A direct connection from a core module to a DPD module indicates that the corresponding core pathway promotes (green arrow) or inhibits (red blocked arrow) the phenotypic process represented by the DPD: Suppression of apoptosis 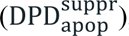, immunoresistance (DPD_imres_), oncogenicity (DPD_onc_), or patient risk (DPD_risk_).

#### RT4 network reconstruction

RT4 oncogenicity appears to be driven by mTOR signaling. This pathway also induces apoptosis, as reflected by the negative global changes in the 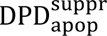 (Fig. 4b). Across all global DPD responses, DPD_imres_ anti-correlated with DPD_onc_, indicating that pathways driving oncogenicity simultaneously suppress immunoresistance.

#### J82 and HT1376 network reconstruction

Both invasive and immunoresistant cell lines, J82 and HT1376, are strongly driven toward oncogenicity and immunoresistance by SMAD3 signaling. In the J82 network, SMAD3 has direct, positive local connections to 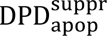, DPD_onc_, and DPD_imres_, all of which exhibit robust positive systems-level responses to pathway activation (Fig. 4c). J82 survival is further supported by RAC-CDC42, which suppresses apoptosis at both local and systems levels by increasing the global DPD^suppr^ response. RAC-CDC42 signaling also promotes DPD_imres_ and DPD_onc_ at the global level, likely via SMAD3 as shown in the local connections (Fig. 4c, right panel). Furthermore, a slight increase in DPD_onc_ upon PI3K and mTOR signaling reflects the reliance of this oncogenic phenotype on both these pathways. As in J82 cells, both oncogenicity and immunoresistance of the HT1376 line is globally supported by SMAD3. Global HT1376 network analysis suggests that JNK inhibits immunoresistance, whereas SMAD3, RAC-CDC42, and FGFR promote it. Notably, both invasive HT1376 and J82 cells exhibit a similar apoptotic profile. In both these lines, the activities of the TGFβ/SMAD3 and RAC-CDC42 pathways suppress apoptosis, whereas JNK activates it (Fig. 4c, d). Finally, both SMAD3 and RAC-CDC42 increase immunoresistance in these invasive cell lines.

#### BC-3C network reconstruction via integration of transcriptomic responses

The core regulatory network for the invasive, basal-like BC-3C line was constructed by integrating in-house bulk RNA-seq data following SMAD3 inhibition with transcriptomic data from the LINCS database and our computed STVs (STV^mouse^, STV_risk_) in the transcriptomic space as described previously^50^. This integrated dataset captured cellular responses to pharmacological perturbations of CDK1/2, CDK4/6, ERBB1/2, Estrogen, FGFR, PI3K, p53, TOP2A, SRC and SMAD3. Analysis of the resulting network revealed that DPDs, representing oncogenicity and poor clinical prognosis, exhibit strong positive systems-level responses to both PI3K and SMAD3 signaling. The mouse scRNA-seq-derived DPD^mouse^similarly predicted strong positive global and local responses to PI3K and SMAD3 signaling, matching the human clinical trial-derived bulk RNA-seq-derived DPD_risk_ (Extended Data Fig. 7c,d). These findings indicate that BC-3C, like the J82 and HT1376 lines, uses SMAD3 signaling to drive its oncogenic phenotype and immunoresistance (Fig. 4e).

In summary, cSTAR-driven reconstruction of causal regulatory network connections across non-malignant and BLCA lines allowed us to determine the systems-level effects of key pathway activations on cellular phenotypes. This analysis identified the SMAD3 pathway as a central driver of both oncogenicity and immune evasion in immunoresistant cell lines, where it concurrently acts as a potent suppressor of apoptosis.

### SMAD3 INHIBITION OVERCOMES IMMUNORESISTANCE IN BLCA CELL LINES CO-CULTURED WITH JURKAT-PD-1 AND PRIMARY HUMAN CD4⁺ T CELLS

cSTAR analysis predicts that activation of the pathways with negative DPD_imres_ scores, such as JNK, will increase immunosensitivity (Fig. 4). In contrast, inhibiting pathways with positive DPD_imres_ scores, is predicted to increase immunosensitivity. Therefore, we hypothesized that targeting the SMAD3, RAC-CDC42, or JNK pathways in immunoresistant J82 and HT1376 cell lines would sensitize them to activated Jurkat-PD-1-mediated cytotoxicity (Fig.4c,d). To validate these predictions, we primed J82 and HT1376 cells for 24 hours with the SMAD3 inhibitor SIS3, the RAC-CDC42 inhibitor MBQ-167, or the JNK inhibitor IX (which functions as a JNK activator in our system, so hereafter is referred to as “JNK IX”). Following this priming, the cancer cells were co-cultured with PMA/ionomycin-activated Jurkat-PD-1 cells, and the synergy between drug treatment and T cell cytotoxicity was measured after 72 hours.

In J82 cells, the combination of SIS3 with activated Jurkat-PD-1 displayed strong synergy, yielding a Loewe synergy score greater than 10 and concave Loewe isoboles (Fig. 5a). In contrast, activated Jurkat-PD-1 combinations with JNK IX or MBQ-167 exhibited lower (although still significantly positive) synergy scores, along with concave Loewe isoboles (Fig. 5a). Importantly, Jurkat-PD-1-drug synergy was markedly stronger with the SMAD3 inhibitor SIS3 than with the TGFβR inhibitor SB525334 (Fig. 5a, Extended Data Fig. 8a).

**Figure 5.**
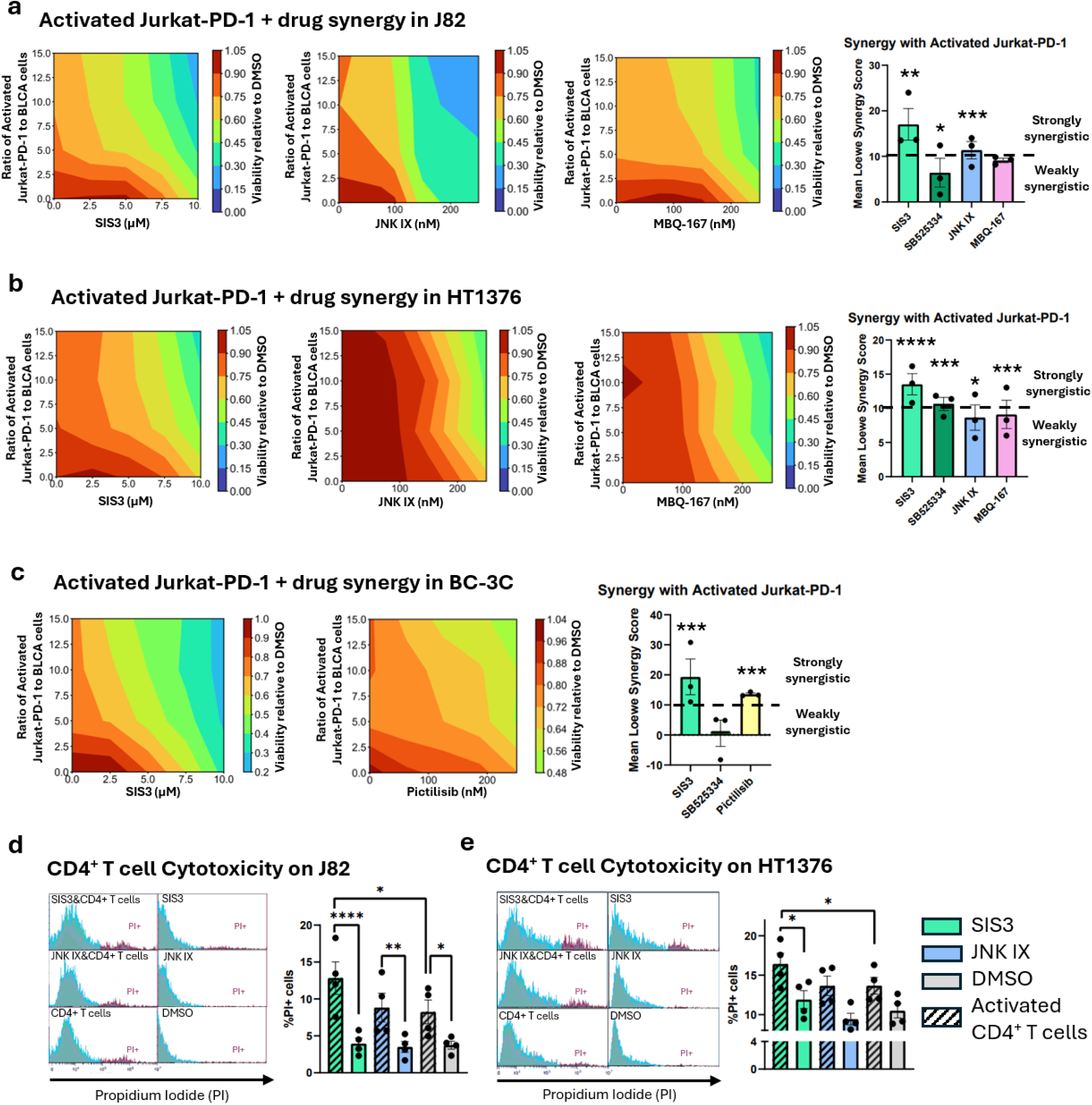
Experimental validation of cSTAR-inferred network predictions reveals synergy between SMAD3 inhibitor and activated Jurkat-PD-1 or primary human CD4⁺ T cells. **a-c.** J82 (**a**), HT1376 (**b**), or BC-3C cells (**c**) were pre-treated with drugs at 4 or 5 different concentrations for 24 h, followed by 72 h co-culture with PMA/Ionomycin-activated Jurkat-PD-1 cells in the continued presence of drugs. Activated Jurkat-PD-1 cells were then mechanically detached, and BLCA cell viability was assessed by MTS assay. Loewe isobolograms are shown for each combination of a drug with activated-Jurkat-PD-1 cells, convex and concave curves indicate synergy or antagonism, respectively. Additionally, synergy was quantified using Loewe synergy scores calculated by the SynergyFinder together with the corresponding p-values. Scores >0 indicate synergy and scores >10 indicate strong synergy. Because JNK inhibitor IX paradoxically activates the JNK pathway (main text), it is hereafter referred to as JNK IX. The other drugs tested were SIS3 (SMAD3 inhibitor), SB525334 (TGFβR inhibitor, Loewe isoboles shown in Extended Data Fig. 8), MBQ-167 (RAC-CDC42 inhibitor), and Pictilisib (PI3K inhibitor). Mean Loewe synergy values are mean of n=3 independent experiments for drug and Jurkat-PD-1 synergy assays, and Loewe isoboles represent an averaged summary of n=3 independent experiments. **d-e.** J82 (**d**) and HT1376 (**e**) cells were pre-treated for 24 h with 10 µM SIS3, 100 nM JNK IX (for J82) or 250 nM JNK IX (for HT1376), or DMSO control, followed by 72 h co-culture with CD4^+^ T cells activated with PMA/Ionomycin. The drug concentrations were selected based on the strongest synergy observed in activated Jurkat-PD-1 co-culture. Activated CD4⁺ T cell–mediated cytotoxicity in J82 and HT1376 cells was measured by propidium iodide (PI) staining and flow cytometry. n=4 independent donors for CD4⁺ T cell cytotoxicity assays. *p<0.05, **p<0.01, ***p<0.001, ****p<0.0001. Drug-Jurkat-PD-1 synergy significance was determined using SynergyFinder as described previously^118^, and CD4^+^ T cell significance measured using one-way ANOVA followed by Tukey’s correction.

A similar pattern was observed in HT1376 cells, where SIS3 and activated Jurkat-PD-1 cells showed strong synergy (Fig. 5b). By contrast, synergy with activated Jurkat-PD-1 cells was consistently weaker for the TGFβR inhibitor SB525334 than for the SMAD3 inhibitor SIS3 (Fig. 5b, Extended Data Fig. 8b). The JNK IX + Jurkat-PD-1 and MBQ-167 + Jurkat-PD-1 (Fig. 5b) combinations showed substantially weaker synergy.

Consistent with our observations in other immunoresistant lines, the combination of SIS3 and activated Jurkat-PD-1 cells was strongly synergistic in killing BC-3C cells, while the combination of SB525334 and activated Jurkat-PD-1 cells was not synergistic (Fig. 5c and Extended Data Fig. 8c). PI3K inhibition using Pictilisib was also synergistic with activated Jurkat-PD-1 cells in BC-3C co-culture (Fig. 5c), further validating cSTAR predictions. Crucially, these results demonstrate that the SMAD3 inhibitor SIS3 strongly potentiates Jurkat-PD-1 cytotoxicity across immunoresistant BLCA lines, whereas JNK IX and the RAC-CDC42 inhibitor MBQ-167 provide only weak sensitization, and the PI3K inhibitor Pictilisib sensitizes only BC-3C cells.

We then analyzed how these treatments affected Jurkat-PD-1 activity and survival. Although JNK IX and MBQ-167 reduced the number of activated Jurkat-PD-1 cells (Extended Data Fig. 9a), all drugs tested slightly increased overall Jurkat-PD-1 activation (Extended Data Fig. 9b), revealing a distinction between the number of activated cells and their overall activation state. All drugs tested also increased early apoptosis in the activated Jurkat-PD-1 cells (Extended Data Fig. 9c), but only JNK IX and MBQ-167 increased late apoptosis (Extended Data Fig. 9d), indicating that the apoptosis of Jurkat-PD-1 cells following TGFβR and SMAD3 inhibition did not dramatically diminish their cytotoxic capability.

Although almost all model predictions were successfully validated experimentally in immunoresistant cells, one prediction remained inconsistent with our observations. The model predicted that FGFR inhibition in HT1376 cells would produce slight immunosensitization, whereas co-culture experiments revealed inhibitor concentration-dependent effects. Mild synergy (positive Loewe synergy scores <10) was observed at low concentrations, but antagonism (negative Loewe synergy scores <−10) was observed at high concentrations of the FGFR inhibitor Infigratinib and activated Jurkat-PD-1 cells (Extended Data Fig. 10a). This discrepancy might be explained by FGFR inhibition also impairing Jurkat cell proliferation^60^ and, potentially, affecting their activity^61^.

Some additional discrepancies emerged between cSTAR predictions and experimental validation in the immunosensitive, non-malignant HCV-29 cell line. We found that Jurkat-PD-1 cell deactivation and apoptosis, and effects of residual PMA/Ionomycin specific to HCV-29 were most likely behind these discrepancies (Supplementary Text, Extended Data Figs. 10-11). Thus, these discrepancies appear to reflect cell line-specific experimental effects rather than a general limitation of our approach, supporting the robustness of the cSTAR predictions of cell state transitions in the immunoresistant cell lines J82, HT1376, and BC-3C.

#### Validating the effect of SMAD3 inhibition on BLCA cell immunoresistance using primary T cells

While Jurkat-PD-1 cells provided a useful model of CD4⁺ T cells, their leukemic origin could potentially lead to divergent pharmacological responses compared to primary cells. To confirm that SIS3 enhances primary CD4^+^ T cell cytotoxicity, we activated primary CD4⁺ T cells from four independent human PBMC donors and co-cultured them with the J82 and HT1376 immunoresistant cells at a 2:1 effector-to-target ratio. In J82 cells, SIS3 significantly enhanced primary CD4⁺ T cell-mediated cytotoxicity, whereas JNK IX had no effect. This sensitization was reflected by a substantial increase in propidium iodide (PI+) staining and decrease in the J82 cell population, as measured by flow cytometry (Fig. 5d). Similarly, in HT1376 cells, the combination of SIS3 and activated primary CD4^+^ T cells was substantially more cytotoxic than either treatment alone as measured by both PI+ staining and percentage of HT1376 cells, whereas no such potentiation was observed for JNK IX (Fig. 5e). Importantly, neither SIS3 nor JNK IX induced cell death in the primary CD4⁺ T cells (Extended Data Fig. 12). This suggests that JNK IX-induced cell death previously observed in Jurkat-PD-1 cells was likely related to their leukemic phenotype. Ultimately, these results confirm that the ability of SMAD3 inhibition to reverse immunoresistance is a robust phenomenon that is consistent across both a Jurkat-PD-1 model and primary human CD4^+^ T cells.

In summary, these experiments validate a central cSTAR prediction that SMAD3 inhibition effectively overcomes the immunoresistance of invasive BLCA cell lines. Furthermore, our findings provide a therapeutic rationale for combining SMAD3 inhibitors with immunoactivating agents, such as immune checkpoint inhibitors, to restore T cell-mediated antitumor activity.

### SMAD3 INHIBITION SENSITIZES BLCA CELLS TO JURKAT-PD-1 CYTOTOXICITY VIA MULTIPLE CELL DEATH MECHANISMS

To investigate the molecular mechanisms by which the SMAD inhibitor SIS3 sensitizes J82 and HT1376 cells to T cell-mediated cytotoxicity, we analyzed the expression of proteins involved in apoptosis, necroptosis, autophagy, ferroptosis, and cuproptosis. Proteomic analysis following 24-hour SIS3 treatment revealed distinct alterations in both immunoresistant lines. In J82 cells, we observed a strong upregulation of pro-apoptotic proteins BAD and TNFRSF10B, the autophagy inhibitor SQSTM1, and several ferroptosis-related markers, accompanied by a significant downregulation of autophagy-related proteins ATG101, ATG4B, and ATG9A (Extended Data Fig. 13a). In HT1376 cells, SIS3 induced the upregulation of BAX, alongside the increase in several necroptosis-related proteins, SQSTM1, and the cuproptosis marker DLAT (Extended Data Fig. 13b). These data suggest that SIS3 sensitization of immunoresistant BLCA cells may involve a combinatorial suppression of autophagy and a simultaneous priming of cells toward multiple cell death pathways.

#### Characterizing the signaling dynamics of SIS3-mediated sensitization

To further investigate these pathways, we performed Western blot analysis of J82 and HT1376 cells co-cultured with activated Jurkat-PD-1 cells following treatments with SIS3, JNK IX, or MBQ-167. To analyze the ferroptosis module, we monitored the inhibition of GPX4, the cysteine transporter SLC7A11, and the Nuclear Receptor Coactivator 4 (NCOA4) known as the selective cargo receptor for ferritinophagy, an iron-dependent form of autophagy^62^. For immune signaling pathways, we evaluated phosphorylated STAT1 (p-STAT1) as a marker of antitumorigenic canonical IFNγ activation and phosphorylated STAT3 (p-STAT3) as a pro-tumorigenic marker involved in anti-inflammatory responses^63^. Finally, we assessed alternative cell death modalities, including autophagy (phosphorylated SQSTM1/p62), necroptosis (phosphorylated MLKL), and overall cell death (NINJ1).

In J82 cells, MBQ-167 treatment alone activated p-STAT1 expression. SIS3 and SIS3+activated Jurkat-PD-1 co-culture led to a decrease in p-STAT3 expression relative to their respective controls. Notably, the levels of the ferroptosis inhibitor Xct and ferritinophagy marker NCOA4 were significantly reduced in J82 cells following the SIS3 + Jurkat-PD-1 treatment compared to SIS3 alone. p-SQSTM1/p62 was also decreased in SIS3+activated Jurkat-PD-1 relative to SIS3 alone, suggesting inhibition of autophagy (Extended Data Fig. 13c). In HT1376 cells, p-STAT1 and p-STAT3 levels increased across all activated Jurkat-PD-1 treatment groups, indicating a broad inflammatory response to Jurkat-PD-1 co-culture that is distinct from the J82 response. p-SQSTM1/p62 was increased following SIS3 treatment, and the decrease in p-SQSTM1/p62 following SIS3+activated Jurkat-PD-1 approached significance compared to SIS3 alone. Similarly to J82, the combination of SIS3+activated Jurkat-PD-1 significantly decreased NCOA4 and increased p-MLKL compared to SIS3 alone, indicating a pro-necroptotic and anti-autophagic response to SIS3+activated Jurkat-PD-1 in HT1376 cells. The levels of the ferroptosis inhibitors GPX4 and Xct were increased in HT1376 following SIS3 treatment, and GPX4 was significantly reduced following the combination of SIS3+activatedJurkat-PD-1relative to SIS3 alone (Extended Data Fig. 13d).

SMAD3 inhibition by SIS3 does not sensitize J82 and HT1376 cells to ferroptosis-induced cell death. Despite the downregulation of proteins involved in protection against ferroptosis, including the cystine transporter xCT, the ferritinophagy marker NCOA4, and the master regulator GPX4, both lipid peroxidation assays following co-culture with activated Jurkat-PD-1 cells in the presence of SIS3 and antagonism of SIS3 with the GPX4 inhibitor RSL3 indicated a trend toward decreased ferroptosis (Extended Data Fig. 14a,b). These findings suggest that the downregulation of xCT, NCOA4, and GPX4 does not translate into increased ferroptotic cell death. Instead, it may accompany broader alterations in autophagy, ferritinophagy, and antioxidant defenses that favor necroptosis. Thus, SMAD3 inhibition appears to sensitize BLCA cells to T-cell cytotoxicity primarily by weakening cellular defenses through impaired autophagy and ferritinophagy, thereby promoting necroptosis and committing BLCA cells toward a cell death fate.

Altogether, our experimental results demonstrate that SMAD3 inhibition effectively overcomes the immunoresistant phenotype of invasive BLCA cells. Rather than acting through a single, dominant signaling axis, SIS3-mediated sensitization involves the coordinated modulation of multiple programmed cell death pathways, indicating a multifaceted mechanism that is consistent with the integrated, systems-level predictions generated by the cSTAR framework.

## DISCUSSION

Productive T cell-mediated tumor eradication relies on simultaneous T effector activation and target cell vulnerability to cytolytic factors. To date, investigations into the limited efficacy of anti-PD-1/PD-L1 immunotherapies have largely centered on upstream immune dysfunction, focusing on pathways that suppress T cell activation, promote exhaustion, and impair cytotoxic function^64–69^. In BLCA, this paradigm is reflected in the extensive development of checkpoint inhibitors targeting PD-1^1,10,13,16,70–73^, CTLA^74,75^, and Siglecs^76^, alongside ongoing neoantigen research to enhance immune recruitment^77,78^. This activation-centric landscape is further highlighted by the recent approval of Nogapendekin alfa-inbakicept (NAI), an IL-15 agonist, for BCG-unresponsive high-risk papillary NMIBC^79^.

CD8^+^ T cells canonically mediate anti-tumor immunity. However, recent studies have increasingly focused on CD4^+^ T cells in BLCA. BCG induces recruitment of BCG-specific CD4^+^ T cells into the bladder, and CD4^+^ T cells are more abundant than CD8^+^ T cells^80^. CD4^+^ T cell density was significantly higher in patients with nonrecurrent upper tract urothelial carcinoma compared to patients with recurrent disease^81^. NF-κB signaling pathway emerged from our PROGENY pathway analysis of TCGA-BLCA transcriptomics data, which is likely indicative of T cell infiltration and cytotoxic activity in the tumor. Although Jurkat cells have been used as a model of T cell-mediated cytotoxicity in BLCA cells^28^, this is the first report demonstrating the utility of PD-1-expressing Jurkat cells to mirror human CD4^+^ T cell cytotoxic behavior.

The prevailing literature on anti-tumor immunity has largely focused on the potency and activation of cytotoxic lymphocytes, with comparatively less attention given to the intrinsic resistance of cancer cells to T-cell-mediated cytotoxicity, even when effector cells are fully activated. Here, we address this gap by using the cSTAR framework to characterize tumor-intrinsic molecular mechanisms that determine the response of bladder cancer (BLCA) cells to cytotoxicity of activated T cells. Previously applied to network reconstruction in breast cancer⁵¹, neuroblastoma³⁰, and atherosclerosis⁵⁰, cSTAR provides a general framework for investigating disease-associated cell state transitions.

A key feature of cSTAR is its use of dynamic phenotype descriptors (DPDs) as quantitative indicators of phenotypic state transitions, linking molecular profiles to biologically and clinically relevant outcomes. Even though the transcriptomics-based DPD_risk_ was generated using TCGA patient data where the patients did not receive immunotherapy, DPD_risk_ efficiently stratified patient survival in independent cohorts treated with immune checkpoint inhibitors. Furthermore, the proteomics-based DPD_imres_, derived from BLCA cell line data, successfully stratified survival in patients with non-muscle-invasive or muscle-invasive bladder cancer. These findings suggest that our DPDs capture tumor-intrinsic cell state changes that are informative of patient outcomes following immunotherapy and other treatments.

Applying these descriptors in cSTAR-reconstructed signaling networks, we identified SMAD3 as a key driver of both BLCA progression, including the acquisition of more aggressive phenotypes, and tumor-cell resistance to T-cell-mediated cytotoxicity. Together, our results extend the focus of anti-tumor immunity beyond the activation and potency of effector lymphocytes. Successful immunotherapy may also depend on shifting cancer cells toward an immunosensitive state, and not only on T cell immune activation.

TGFβ signaling has also previously been shown to induce resistance to T cell cytotoxic factors in the cancer cells^82^. We demonstrate that only SMAD3 inhibition, and not TGFβ receptor inhibition, consistently overcomes BLCA immunoresistance. Our cSTAR-driven reconstruction of causal network connections in J82 and in HT1376 cells, suggested that SMAD3 is the only immediate driver of immunoresistance. In J82 cells other modules and pathways, such as RAC-CDC42, JNK, mTOR and others, affect immunoresistance only via activating or inhibiting SMAD3. In HT1376 cells, JNK also displays an immediate connection to immunoresistance, which is negative, and all effects of other modules and pathways are mediated by changing activity of either SMAD3 or JNK. In BC-3C cells, immediate connections to the DPD_risk_ modules come from PI3K, SMAD3 and SRC modules. Thus, in all immunoresistant cells studied SMAD3 serves a key module, and in some cases the only module, which integrates signals from other modules and shapes the immunoresistance whole-cell proteomic pattern. This finding potentially explains why inhibiting SMAD3 exerts a much more profound effect on immunoresistance than inhibiting TGFβR, as SMAD3 can be activated by multiple intracellular pathways beyond TGFβR alone.

SMAD3 inhibition has a dual function of preventing T cell exhaustion by PD-1 and inhibiting the immunoresistance state transitions in BLCA cells. SMAD3 functions downstream of TGFβ signaling in BLCA cells^83–85^. SMAD3 is also phosphorylated and inhibited by CDK2, and PD-1 induces SMAD3 activation via impairment of CDK2 activity in T cells^86,87^. Notably, while TGFβ receptor inhibition, SMAD3 inhibition, JNK activation, and RAC-CDC42 inhibition all increase the overall activation of Jurkat-PD-1 cells in culture (Extended Data Fig. 9b), only SMAD3 inhibition emerges as an immunosensitizer across all immunoresistant cell lines. Although we cannot completely disentangle the effects of each drug treatment on cancer cells from those on Jurkat-PD-1 or CD4^+^ T cells in our co-culture model, it is important to consider the dual effects of drug treatments on both immune and cancer cell populations. These observations underscore that immunotherapy response is jointly determined by the functional state of infiltrating immune cells and the cell-intrinsic signaling state of the tumor cells.

Previous reports have linked a low TGFβ transcriptional signature to better responses to anti-PD-L1 immunotherapy, likely due to T cell exclusion by extracellular matrix deposited by stromal cells activated by TGFβ^88–90^. However, while TGFβ receptor inhibitors have been tested clinically as monotherapies and in combination with immunotherapies, most trials were halted due to severe toxicity^91^. Our findings suggest that TGFβ-induced immunoresistance is mediated predominantly through SMAD3 or its downstream effectors, providing a rationale for selective SMAD3 targeting as a promising alternative to upstream receptor blockade that may bypass the severe toxicities associated with TGFβ receptor inhibition.

To clarify how immune cytotoxicity synergizes with BLCA cell sensitivity, we explored cell death mechanisms in our co-culture model. We showed that direct cell-cell contact enhances Jurkat-PD-1 cytotoxicity, as cytotoxicity against cell lines in the panel was reduced in the Transwell system, where direct contact is prevented and cancer cells are exposed only to factors released by the immune cells, including soluble mediators such as Fas and TRAIL^25^ (Extended Data Fig. 2b). The PROGENY analysis of TCGA-BLCA data suggested that the tumor cell death machinery may involve JAK/STAT signaling, which is activated downstream of IFNɣ. A previous report and our data support the dependency of anti-tumor immunity on IFNɣ receptor expression on BLCA cells, but not MHC Class II^92^. We demonstrated that STAT1 and STAT3, which are activated and phosphorylated downstream of IFNɣ receptor engagement and JAK signaling, were upregulated in HT1376 cells following activated Jurkat-PD-1 co-culture. The ratio of phosphorylated STAT1 to phosphorylated STAT3 has been proposed to predict immunotherapy response in other cancers^93^ and this ratio is largely unchanged in our co-culture system. This suggests that the IFNɣ pathway is upregulated in our co-cultures but does not likely influence immunosensitivity. Altogether, BLCA cells integrate a myriad of cell fate decision signals that point toward cell death.

The route to cell death can be visualized as a 3D topological landscape, where a variety of cell death signals can push a cell toward different programmed cell death pathways such as apoptosis, ferroptosis, pyroptosis, or necroptosis^39,94,95^. Beyond apoptosis, ferroptosis, necroptosis, and cuproptosis are also involved in immunotherapy sensitivity in *in vitro* models of other cancers as well as BLCA^96–102^. We observe decreases in ferroptosis inhibitors but not activated ferroptosis. Instead, our protein expression data show inhibition of ferritinophagy and autophagy and activation of necroptosis are common in both J82 and HT1376 cells.

Autophagy has been shown to induce PD-L1 expression in BLCA cells^103^ and promote cancer progression in other cancers^104^. We propose that the main route by which SIS3 immunosensitizes BLCA cells is by downregulation of ferritinophagy and autophagy, which removes the brakes for the BLCA cells to descend the cell death landscape following multiple cell death paths. Our study endorses the exploration of the utility of SMAD3 inhibition for simultaneous prevention of TGFβ-induced tumor progression and autophagy-induced tumor protection.

### CONCLUSION

cSTAR-based core molecular signaling network reconstruction identified that SMAD3 is a key driver of oncogenic transformation and immunoresistance in immunoresistant BLCA cells. SMAD3 inhibition sensitizes BLCA cells to CD4^+^ T cell-mediated cytotoxicity by suppressing autophagy and driving cells toward alternative cell death pathways. Crucially, cSTAR captures the transition from immunoresistance to immunosensitivity and offers avenues for pharmacological exploitation of BLCA cell vulnerabilities along this transition. Failure of immunotherapy may therefore reflect not only impaired T cell function, but also the adoption of cancer cell states that are intrinsically resistant to cytotoxic lymphocytes. Understanding the molecular mechanisms that sustain these states may reveal therapeutic strategies to overcome tumor-intrinsic immunoresistance and enhance immunotherapy efficacy.

## MATERIALS AND METHODS

### BLCA AND JURKAT CELL CULTURE

HCV-29, RT4, T24, J82, and HT1376 cells were a generous gift from Lars Dyrskjøt Andersen, and BC-3C cells were purchased from DSMZ. Cell culture media for HCV-29 cells was Dulbecco’s modified eagle’s medium (DMEM, Gibco) + 10% fetal bovine serum (FBS, Gibco). Minimum Essential Medium (MEM, Gibco) + 10% FBS + 2% L-glutamine (Gibco) + 1% Non-essential amino acids (Gibco) was used for J82 and HT1376 cells. McCoy’s 5A (Modified) Medium (Gibco) + 10% FBS + 1% L-glutamine was used for RT4, T24, and BC-3C cells. Jurkat-PD-1 and all T cell stimulator (TCS) cells were a generous gift from Peter Steinberger^32^. RPMI 1640 Medium + 10% heat-inactivated FBS + 1% L-glutamine was used for all Jurkat and TCS cells (hereafter referred to as “Jurkat media”). For co-culture of BLCA and Jurkat cells, all BLCA cell lines were gradually weaned onto Jurkat media, passaging twice at increasing concentrations of Jurkat media relative to the original BLCA cell media. The identity of the HCV-29, RT4, T24, J82, and HT1376 cell lines were verified using STR profiling by the Eurofins Cell Line Authentication (CLA) service. All cells were grown without antibiotics and tested negative in monthly mycoplasma checks.

### ACTIVATION AND FLOW CYTOMETRY OF JURKAT CELLS

Jurkat-PD-1 activation methods included 1) 24-hour treatment with anti-CD3/CD28 Dynabeads (Thermo Fisher) 2) 3-hour treatment with 100 nM each Phorbol-12-myristate-13-acetate (PMA, Sigma-Aldrich) and Ionomycin calcium salt from Streptomyces conglobatus (Sigma-Aldrich) followed by 3 washes with Jurkat media to remove the PMA/Ionomycin, or 3) 24-hour co-incubation with T cell stimulator (TCS) at a 1:4 ratio (TCS: Jurkat-PD-1) cells. PD-1 expression in Jurkat-PD-1 cells was validated using TCS and TCS expressing PD-L1 (TCS-PD-L1). Jurkat-PD-1 cells co-cultured with TCS-PD-L1 were also treated with 25 ng/ml of the anti-PD-1 immune checkpoint inhibitor pembrolizumab.

TCS cells of mouse origin were separated from Jurkat-PD-1 cells of human origin by flow cytometry using an anti-human CD45 antibody conjugated to APC-H7 (Biolegend). Activation and apoptosis of Jurkat-PD-1 cells were measured by flow cytometry using an APC Annexin V Apoptosis Detection Kit with 7-AAD (BioLegend) according to the manufacturer’s instructions. Surface CD4 and PD-1 expression was tested on Jurkat-PD-1 cells across three passages post-thaw by staining Jurkat-PD-1 cells with an anti-CD4 antibody conjugated to PE (BioLegend) and an anti-PD-1 antibody conjugated to Brilliant Violet 421 (BioLegend) for 20 min in Annexin V staining buffer (BioLegend), followed by washing with Annexin V staining buffer and addition of 7-AAD viability dye (BioLegend) to stain dead cells. All flow experiments were conducted using a Cytoflex LX cytometer. Fluorescence Minus One (FMO) controls were used to determine gates for each fluorophore. UltraComp eBeads^TM^ were used for all antibody single-color compensation controls, and plain Jurkat and Jurkat-PD-1 cells were used for 7-AAD, GFP, and Annexin V single-color compensation controls. All flow data was analyzed using CytExpert software (Beckman Coulter).

### VIABILITY DOSE-RESPONSE ASSAYS

BLCA cells were plated at a density of 7500 cells per well for HCV-29, 10,000 cells per well for RT4, 2500 cells per well for J82 and T24, and 5000 cells per well in 96 well plates for HT1376 and BC-3C and allowed to attach for 24 hours. Initial viability was measured by MTS Following PMA/Ionomycin activation (activated Jurkat-PD-1) or equivalent concentration of DMSO (0.01%) (non-activated Jurkat-PD-1) and subsequent washing of Jurkat-PD-1 cells, BLCA cells were co-cultured with Jurkat-PD-1 cells at a ratio of 10:1 (Jurkat-PD-1: BLCA cell) or drugs (Extended Data Table S1) for 72 hours in duplicate. To determine the effects of residual PMA/Ionomycin following Jurkat-PD-1 activation, the final wash was added directly to the BLCA and control HCV-29 cells. Jurkat-PD-1 cells were then detached from BLCA cells by carefully pipetting to detach, washed once with PBS, and visually inspected to confirm removal of Jurkat-PD-1 but minimal detachment of BLCA cells. Viability was measured by MTS assay (CellTiter 96 ® Aqueous MTS Reagent Powder, Promega) according to the manufacturer’s instructions and absorbance read on a plate reader at 490 nm using a SpectraMax M3 microplate reader (Molecular Devices). Viability was then calculated relative to the respective DMSO control for each cell line. For each drug and co-culture condition, at least 3 independent experiments were conducted. Area Under the Curve (AUC) was calculated using GraphPad Prism, then drug normalized by dividing by the drug dose.

### WHOLE-CELL PROTEOMICS

#### Sample preparation (initial proteomics experiment)

All cell lines were seeded in triplicate in 10 cm dishes at 2.2 million cells per plate (HCV-29), 4 million cells per plate (RT4), 2.5 million cells per plate (T24) 3 million cells per plate (J82), or 4 million cells per plate (HT1376 and BC-3C). Cells were cultured for 48 hours, harvested using Trypsin-EDTA 0.05% (Gibco), washed with ice-cold PBS, liquid aspirated, and pellets flash-frozen at −80°C. Pellets were then lysed by sonication in a lysis buffer comprised of 8M urea in 50 mM Tris-HCl with phosphatase and proteinase inhibitors (Roche). The protein concentration of cell lysate was measured using the Pierce^TM^ BCA protein assay (Thermo Fisher Scientific) according to the manufacturer’s instructions. 400 µg total protein was adjusted to a total volume of 50 µL. The proteins were then reduced with 8 mM of dithiothreitol and mixed for 30 min at room temperature, then carboxylated using 20 mM iodoacetamide and mixed for 30 min in the dark at room temperature. The proteins were then diluted in 500 µL of 50 mM Tris-HCl and digested with 20 µg Trypsin/Lys-C (Promega, V5073) diluted in Tris-HCl and mixed overnight at 37°C. Digestion was terminated by adding trifluoroacetic acid to 1% and cleaned using Sep-Pak (50mg) c18 cartridges (WAT054955; Waters) then dried and stored at −80°C for proteomic analysis.

#### Sample preparation (drug perturbation experiment)

For the drug perturbation experiment, all cells were seeded in 10 cm dishes at 3 million cells per dish (HCV-29), 3.5 million cells per dish (RT4), 2.5 million cells per dish (J82), and 4 million cells per dish (HT1376). The cells were allowed to adhere for 24 hours. Each cell line was then individually treated in triplicate with the drugs and concentrations listed in Extended Data Table S1, or with 0.1% DMSO as a control. Each experiment included HCV-29 + 0.1% DMSO samples in triplicate to act as a batch control. After 24 hours of drug treatment, cells were rinsed in ice-cold PBS, lysed directly on the plate with 1x RIPA buffer (Brennan & Company) supplemented with protease and phosphatase inhibitors, and collected using a cell scraper. Cells were then incubated on ice for 30 min to facilitate lysis, centrifuged at 15,000g for 10 min at 4°C, and the supernatant was collected. Protein was measured using the Pierce^TM^ BCA protein assay (Thermo Fisher Scientific) according to the manufacturer’s instructions. 50 µg of total protein per sample were digested using the Protein Aggregation capture (PAC) method according to^105^ using the KingFisher Flex (Themo Scientific). Briefly, proteins were reduced with 8 mM of dithiothreitol and mixed for 30 min at room temperature, followed by carboxylation using 20 mM iodoacetamide and shaking for 30 min in the dark at room temperature. Protein aggregates were then bound to MagReSyn Hydroxyl beads (ReSyn Biosciences, MR-HYX010) in 70% acetonitrile. Samples were then washed in 3 changes of 500 µL 95% acetonitrile, followed by two washes in 500 µL 70% EtOH, and finally digested using sequencing-grade trypsin (Promega, V5111).

#### Mass spectrometry

1 µg of each proteome sample from the initial experiment, and a pooled sample from both experiments, was analyzed using a Bruker timsTOF Pro mass spectrometer connected to an Evosep One liquid chromatography system. Tryptic peptides were resuspended in 0.1% formic acid and loaded onto an Evosep tip. The Evosep was configured to pick up each tip, elute and separate the peptides using a set chromatography method^106^. The mass spectrometer was operated in positive ion mode with a capillary voltage of 1200–1700 V, dry gas flow of 3 l/min and a dry temperature of 180°C. All data was initially acquired with the instrument operating in a data-dependent analysis parallel accumulation serial fragmentation mode (dda-PASEF). Trapped ions were selected for ms/ms using parallel accumulation serial fragmentation (PASEF). A scan range of (100-1700 m/z) was performed at a rate of 4 PASEF MS/MS frames to 1 MS scan with a cycle time of 0.53s^107^. The resultant files from the pooled samples were used to create the dia-PASEF method within Bruker timsControl software. Dia-PASEF settings used were: window mass width 26.0 Da, mass overlap 1 Da, mass steps per cycle 33 or 36, mass range 374.3 – 1200.3 or 321.6 – 1222.6 m/z. All samples were subsequently acquired using data-independent analysis parallel accumulation serial fragmentation (dia-PASEF)^108^.

#### Data Analysis

Data acquired using dia-PASEF was analyzed using either DIA-NN 2.0.2 Academia (Data-Independent Acquisition by Neural Networks)^109^ or FragPipe version 22.0^110^. For DIA-NN analysis, the Homo sapiens subset from the UniProt Swiss-Prot database (reviewed, downloaded February 2025) was used to generate a spectral library in library-free mode. Specific search settings included cysteine carbamidomethylation as a fixed modification, methionine oxidation and N-terminal acetylation as variable modifications, N-terminal methionine excision enabled, maximum missed cleavages 1, min precursor +2, max precursor +4. Proteoform scoring was used, cross-run normalisation was set to retention time dependent, and library generation was set to RT profiling. Precursor FDR was set to 1%. For FragPipe analysis, the DIA_SpecLib_Quant_diaPASEF workflow was employed using FragPipe 22.0 with MSFragger 4.1, diaTracer 1.1.5, Philosopher 5.1.1 and IonQuant 1.10.27. A spectral library was generated from the dda-PASEF runs using MSFragger 4.1, with Philosopher 5.1.1 used for target-decoy based FDR filtering. The dia-PASEF data were then quantified against this library using DIA-NN 1.8.2 with smart profiling and peak-centred quantification. All output was filtered at 1% FDR at both the precursor and protein group level.

#### Proteomic data pre-processing

Raw data were log2-transformed and filtered to remove proteins that didn’t achieve at least 67% valid values in at least one group. For the initial proteomics experiment, imputation was performed using Perseus for missing values using normal distribution with a width of 0.3 and a down-shift of 1.8 separately for each column. For the drug perturbation experiment, minProb imputation was performed using custom R code. The dataset was then annotated using GOBP names and KEGG names.

#### Differential protein expression analysis

For proteomics experiments before drug treatments, differential protein expression was assessed in Perseus using multiple ANOVA with false discovery rate (FDR) 5%. The expression levels were then standardized using a Z-score and clustered using hierarchical clustering with Euclidian distances.

For the drug perturbation experiment, differential protein expression relative to HCV-29 + DMSO control was assessed using limma. Protein fold changes were then filtered for adjusted p-value <0.05.

### PERTURBATION REVERSE-PHASE PROTEIN ARRAY (RPPA)

The same protein samples used for proteomics in the drug perturbation experiment were also used for RPPA. 200 µg total protein at a concentration of 2 µg/µL in RIPA buffer was used for RPPA. Samples were prepared for printing by addition of 4X Sample buffer and heat denaturation at 95°C for 5 minutes. All samples were diluted to 1.5 mg/ml (D1), 0.75 mg/ml (D2), 0.375 mg/ml (D3) and 0.1875 mg/ml (D4) using PBS containing 10% Glycerol. Array spotting was carried out with the Quanterix 2470 Arrayer platform using 185μM pins. Each sample was spotted at four dilutions (D1-D4) onto single pad ONCYTE® SuperNOVA nitrocellulose slides at a 500μM spot-to-spot distance. 120 signaling pathway markers (and appropriate controls) listed in Extended Data Table S2 were profiled in a standard RPPA assay as described briefly below. A more detailed procedure is described previously^111^. After spotting, the slides were incubated with antigen retrieval solution (1x Reblot strong) for 10 mins before being placed in a microfluidic structure to individually address the arrays with primary or secondary antibody solutions. Following blocking buffer (Superblock T20) for 10 mins, the detection of marker antibodies was performed in a two-step sequential assay: 1) Incubation of an array with primary analyte-specific antibody in blocking buffer for 60 mins at room temperature, then 2) Removal of excess antibody by washing arrays with PBS-T, followed by a further incubation with blocking buffer and PBS-T washes, and incubation with secondary antibody (Dylight-800-labeled anti-species antibodies diluted 1:2500 in Superblock T20) for 30 mins. After further washing and slide drying, the arrays were imaged in the Innopsys Innoscan 710 scanner. Blank signals were determined by omitting the primary antibody from step 1 and instead incubating the array with Superblock T20 alone, followed by step 2.

Sample loading on arrays was determined by staining one slide with fast green protein dye and scanning at 800nm. Microarray images are analyzed using Mapix software (Innopsys) with feature (spot) diameter of the grid set as 270μm. The average signal intensity was determined for each individual feature and the median background from the adjacent area was subtracted from each feature signal leading to a net signal per feature. Fluorescence intensity for each feature on the array was measured. Quality control was performed for linear fit through the 4-point dilution series for all samples on all arrays using a flag system. The median values from the 4-point dilution series are calculated and used as a measure of fluorescence intensity. Relative Fluorescence Intensity (RFI) values were calculated using relative abundance of total and phosphorylated proteins across the sample set. For each sample, the ratio of median RFI of the dilution series for both the target and fast green stain was calculated. Finally, the data from the 3 technical and biological replicates is averaged and standard deviation values calculated. Each protein measurement was first normalized by the GAPDH level, and then on the value of the same protein in the DMSO control of the same cell line to obtain fold changes.

### BULK MRNA SEQUENCING SAMPLE COLLECTION

BC-3C cells were seeded in 6 well plates at 400,000 cells per well in triplicate for each treatment. After adhering for 24 hours, cells were treated with either 15 µM SB525334 (TGFβR inhibitor) or 10 µM SIS3 (SMAD3 inhibitor) or 0.1% DMSO control. After 24 hours drug treatment, RNA was collected using RNeasy Mini Kit (Qiagen) with genomic DNA removal according to the manufacturer’s instructions. Library preparation, quality control, and differential gene expression analysis steps were conducted by Novogene Europe (Cambridge, UK).

#### Library preparation

Messenger RNA was purified from 1 µg total RNA using poly-T oligo-attached magnetic beads. After fragmentation, first strand cDNA was synthesized using random hexamer primers, followed by second strand cDNA synthesis using dTTP. The library then underwent end repair, A-tailing, adapter ligation, size selection, amplification, and purification. The library was checked with Qubit and real-time PCR for quantification and Bioanalyzer for size distribution detection. After library quality control, different libraries were pooled based on the effective concentration and targeted data amount, then subjected to Illumina sequencing.

#### Quality control

Raw data (raw reads) of fastq format were processed through fastp software. In this step, clean data (clean reads) were obtained by removing reads containing adapter, reads containing ploy-N and low quality reads from raw data. At the same time, Q20, Q30 and GC content the clean data were calculated. All downstream analyses were based on high quality, clean data. The steps of data processing were as follows: (1) Discard a paired read if either one read contains adapter contamination; (2) Discard a paired read if more than 10% of bases are uncertain in either one read; (3) Discard a paired read if the proportion of low quality (Phred quality<=5) bases is over 50% in either one read.

#### RNA-seq Differential Gene Expression analysis

Reference genome and gene model annotation files were downloaded from the genome website. HISAT2 (2.2.1) was used to build the index of the reference genome and align paired-end clean reads to the reference genome. featureCounts (2.0.6) was used to count the read numbers mapped to each gene. FPKM of each gene was calculated based on gene length and reads count mapped to this gene. Differential expression analysis for two conditions/groups was performed using the DESeq2 R package (1.42.0). The resulting p-value is adjusted using the Benjamini and Hochberg’s methods. The threshold of significant differential expression is adjusted p value <= 0.05 and |log2(foldchange)| >= 1.

### STV CONSTRUCTION

#### Building STVs from total proteomics data

For proteomics data, the State Transition Vectors (STVs) were built using classification machine learning methods, as described previously^30^. In brief, we first selected highly variable proteins in perturbation data. Statistically significant (adjusted p-value < 0.05) mass spec proteomics log fold changes from all comparisons were collated into a single table. Proteins with a high standard deviation across treatments (standard deviation > 0.8) in at least one cell line were collected, which totaled 1199 proteins. We then applied the Support Vector Machine (SVM) algorithm with a linear kernel to construct a hyperplane that maximizes the separation between opposing basal cell states (i.e. DMSO control conditions) in the proteomics dataspace. These pairs of cell states include 1) cancerous (RT4, J82, and HT1376 cell lines) versus non-cancerous (HCV-29 cell line), and 2) immunoresistant (J82 and HT1376 cell lines) versus immunosensitive (HCV-29 and RT4 cell lines).

The confusion matrix was diagonal in all cases, indicating error-free separation of the cell states. The normal vectors, which determine the SVM-built hyperplane, were normalized to unit length and were defined as the State Transition Vectors (STVs). We named these vectors STV_onc_ for the cancerous vs. non-cancerous separation and STV_imres_ for the separation between immunoresistant and immunosensitive cell lines.

### CLINICAL DATA ANALYSIS: BUILDING STV_RISK_ FROM BULK RNA-SEQ DATA

For clinical data, the classification methods previously used to construct STVs^30^ are not directly applicable, because clinical survival data do not naturally comprise two distinct patient states. Therefore, to build STV_risk,_ we applied regression methods to find molecular patterns associated with favorable and unfavorable treatment outcomes. Human bulk RNA-seq and clinical data were obtained from The Cancer Genome Atlas Bladder Urothelial Carcinoma (TCGA-BLCA)^112^, Atezolizumab phase II clinical trial IMvigor210^55^, and a cohort with different immune checkpoint treatments^56^ (GEO accession number GSE176307). Perturbation data for the BC-3C cell line, apart from SMAD3 inhibition, were obtained from the LINCS L1000 database^113^. Only the genes present in both bulk RNA-seq data and landmark genes from the LINCS L1000 database were included.

The STV_risk_ was constructed using a Cox Proportional Hazards regression model to evaluate the relationship between patient survival and gene expression levels in the TCGA dataset. Elastic net Cox regression (*Glmnet* R package^42,43^) was applied to transcriptomics data. Briefly, elastic net Cox regression estimates the model coefficients by minimizing the following cost function *K*(***α***)

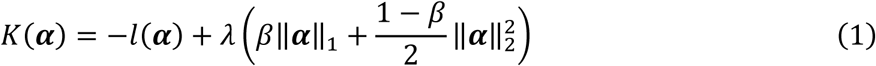

Here, ***α*** is a vector of coefficients, which are being determined in this procedure, *l*(***α***) is the log partial likelihood function for the Cox model, ‖***α***‖_1_is an L1 norm of vector ***α***, and ‖***α***‖_2_is an L2 norm of vector ***α***. For the Cox model, the log partial likelihood function reads

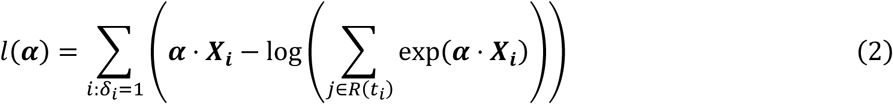

Here ***X_i_*** is a vector of omics data for patient (*i*), *δ_i_* is an indicator function that is 1 if the event of interest has occurred and 0 otherwise, *t_i_* are the event times, and *R*(*t_i_*) denotes the set of all individuals at risk at time *t_i_*. Overall survival is used for Cox regression, thus, event in this case meant death of patient from bladder cancer.

Fitting a Cox model with an elastic net penalty requires selecting optimal values for the regularization parameter *λ* and the mixing parameter *β*. To find optimal *λ* and *β* values, 5-fold cross validation was used, and the concordance index (C-index) was used to evaluate the predictive performance of the survival model. The resulting vector of coefficients was normalized to unit length and defined as an STV_risk_.

A prognostic score was calculated as the dot product of STV_risk_ and the log2-transformed gene expression levels. TCGA transcriptomics data were subsequently used to estimate pathway activities using PROGENY signature^41^. These activities were used as input to the same regularized Cox regression pipeline (*Glmnet* R package^42,43^) to find pathways predictive of patient survival (Fig. 2b).

#### Constructing 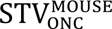 from single-cell RNA-seq data

To construct 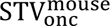 in transcriptomic dataspace, we used classification methods to identify molecular patterns associated with oncogenic transformation in animal models. Mouse bladder urothelial carcinoma scRNA-seq data were obtained from a semi-spontaneous, carcinogen-induced (OH-BBN) murine model of urothelial cancer (Hgf-Cdk4^R24C^ mice)^57^ (the Gene Expression Omnibus (GEO) accession number GSE174182). Data integration and clustering were performed using Seurat (v4.4.1). Murine cells with >25% mitochondrial reads were excluded, and only cells expressing 200–8,000 genes were retained. Datasets were normalized and integrated using SCTransform. Dimensionality reduction was performed via principal component analysis (PCA), followed by UMAP using the first 30 principal components (PCs). Clustering was conducted with the Louvain algorithm using 3,000 variable genes, 30 PCs, a resolution of 0.05, and a random seed of 1.

Cluster selection was guided by urothelial marker genes (Cdh1, Upk1a, Upk1b, Upk2, Upk3a, Ivl), yielding 15,954 bladder urothelial cells. The UMI-corrected matrix was total-count normalized (scaling factor = 10,000) and pseudo-log2 transformed. Cells were labeled according to their origin (healthy control, and bladder cancer) and used as input for the SVM. The resulting normal vector was normalized to unit length and used as the STV of oncogenic transformation 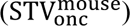.

### DPDs AND GLOBAL RESPONSES OF CORE NETWORK MODULES CALCULATING DYNAMIC PHENOTYPIC DESCRIPTORS (DPDS)

DPD calculations are outlined in detail in our previous publication^30^. Briefly, when an STV is constructed using an SVM with a linear kernel, the DPD of a given data point is the distance from that point to the corresponding separating hyperplane, with the sign indicating the position of the point relative to the STV. The equation defining the DPD is the following,

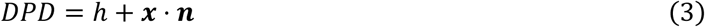

Here ***x*** is a data point, for which a DPD score is calculated, ***n*** is the STV, and ℎ is an intercept for the corresponding separating hyperplane. Eq. (3) was used to calculate the following DPDs: *DPD_onc_* and *^DPD^_imres_*.

For DPDs calculated in the transcriptomic data space, the STVs were built based on different datasets than in-house perturbation data. *STV_risk_* was constructed using patient data, 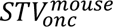 was built using mouse data for the perturbation experiments in the BC-3C cell line^113^. In these cases, the intercept (ℎ) cannot be reliably determined, because of batch effects, although the STV (***n***) can still be reliably determined. Accordingly, the *DPD_risk_* and 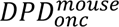 scores for the perturbation data points were calculated as,

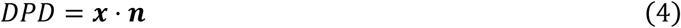

As explained in the main text, for cell lines HCV-29, RT4, J82 and HT1376, we introduced a new DPD for apoptosis suppression 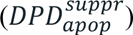 based on the equal contribution of two RPPA antibodies, cleaved Caspase-3 and cleaved PARP. 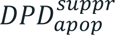 scores were calculated using Eq. 4. All STVs used to calculate the DPD scores are listed in Table 1 and shown in Fig. 3.

### SURVIVAL ANALYSIS OF DPD_IMRES_ AND DPD_RISK_ USING PATIENT PROTEOMIC AND TRANSCRIPTOMIC DATA

To validate the utility of DPD_imres_ for classifying clinical patient proteomic data, we searched PRIDE/ProteomeXchange, iProX, MassIVE, jPOST, the Proteomic Data Commons and Europe PMC for mass spectrometry (MS) proteomic datasets of urothelial carcinoma that included both NMIBC and MIBC patients. Datasets were included if they provided patient-level protein abundance measurements that could be linked to clinical outcomes. Three cohorts met these criteria (Extended Data Table S3).

Each STV entry consists of one or more UniProt accessions and a gene symbol. Entries were mapped to cohort proteins by gene symbol (case-insensitive). If the gene symbol was unavailable, matching was performed using UniProt accession, allowing matches to any accession in the protein group after removal of isoform suffixes. When several cohort rows matched, we used the row with the most non-missing values.

Protein abundances reported on a linear scale were log2-transformed. Each protein was then median-centered across the samples of its cohort, without variance scaling. Missing values were set to zero after centering, which is equivalent to the cohort median, so they did not contribute to the score. Coverage of STV_imres_ was quantified as both the fraction of STV entries matched and the fraction of the squared STV weight carried by the matched entries (Extended Data Table S4).

Because only approximately 17% of the proteins in the STV_imres_ were detected in the Qu et al. (2023) study, its survival analysis (Fig. 3e) is considered supportive evidence only. The limited overlap may partly reflect the use of plasma proteomics in the Qu et al. (2023) study. Nevertheless, the concordance of its survival analysis pattern with those obtained in the Dressler et al. (2024) and Xu et al. (2022) studies supports the conclusions drawn from the latter studies.

For both DPD_imres_ and DPD_risk_, clinical variables were taken from the supplementary tables and source-data files of each publication. Patient overall survival (OS) was harmonized to a time (months) and an event indicator (1 = event, 0 = censored). DPD values were standardized within each cohort using a Z-score, and associations were tested using Cox regression. Univariable models were used to calculate a Hazard Ratio (HR) per 1 SD of DPD with a Wald 95% confidence interval and p-value. The proportional hazards assumption was tested with scaled Schoenfeld residuals. Covariates (Extended Data Tables S3 and S4) with >20% missing values were not used, and complete cases were analyzed. The sign of every nominally significant univariable association was cross-checked against a median-split log-rank comparison. We then compared the C-index of the clinical-only model with that of the clinical + DPD model, and tested the gain with a likelihood-ratio test. Statistical significance was set at a two-sided p < 0.05. As a result, the STV_risk_ yielded a C-index equal to 0.994 for the training TCGA dataset, 0.533 for the IMvigor210 data, and 0.562 for the GSE176307 dataset. STV_imres_ yielded C-indexes of 0.688 for the Dressler et al. 2024 cohort, 0.705 for the Qu et al. 2023 cohort, and 0.742 for the Xu et al. 2022 cohort.

To identify DPD_imres_ and DPD_risk_ thresholds that significantly separate patients into high- and low-risk groups, each cohort and endpoint was analyzed separately. Every observed DPD value was considered as a candidate threshold, provided that at least 20% of patients fell on each side. Each threshold was tested by survival of the two groups on either side of the threshold was compared with the log-rank test. The threshold that provides the most statistically significant separation between high- and low-DPD groups in all cohorts was used to make the Kaplan-Meier curves in Fig. 3. As a reference for the inflation introduced by the search, the maximally selected log-rank statistic^114^ was computed over the same range of thresholds, with p-values from the improved Bonferroni approximation implemented in the maxstat package^115^. Kaplan-Meier curves were then generated with custom R code.

Data harmonization was conducted using Python 3.13 with pandas 2.3.3, NumPy 2.4.1 and SciPy 1.17.0. Survival analysis and statistical analyses were performed in R 4.6.1 with survival 3.8.12 (Cox models, concordance, Kaplan–Meier, log-rank tests), maxstat 0.7.26, survminer 0.5.2 and ggplot2 4.0.3.

#### cSTAR Network Inference

The cSTAR reconstruction of cell-specific signaling networks was based on the acquired RPPA data for the HCV-29, RT4, J82, and HT1376 cell lines. To determine the global (systems-level) responses of signaling modules, we first calculated global response coefficients for the specific RPPA antibodies representing the module outputs, using a second-order differencing scheme as described previously^30^. The global response of each module was then calculated as the weighted sum of the corresponding antibody-level global response coefficients. The antibodies used as module outputs and their weight coefficients are presented in Extended Data Table S5. Responses of the phenotypic DPD modules were calculated using a first-order differencing scheme^30^ applied to mass spectrometry data, except for the apoptosis suppression DPD (DPD^suppr^), which was determined from RPPA data as described above.

For cSTAR reconstruction of the core network in BC-3C cells from transcriptomic data, we used the LINCS L1000 perturbation dataset^113^ together with our in-house SMAD3 inhibition data. Global response coefficients for both core network and DPD modules were calculated as described previously^50^.

To infer cSTAR networks, we used Modular Response Analysis (MRA)^47^. MRA takes as inputs (i) the global response matrix (*R_ij_*), (ii) a prior specifying known and forbidden connections between nodes, and (iii) a perturbation matrix describing the changes in the nodes directly perturbed in each treatment. To improve robustness of network inference, we modified previously developed MRA methods, as follows. We first used Bayesian MRA (BMRA)^47,116^ to identify non-zero connections, and then used Ordinary Least Squares MRA^47,117^, allowing only the connections identified by BMRA. As a result, we obtained local response matrix (***r****=r_ij_*), which described both core network wiring and cell state transitions^30^. According to the MRA theory^59^, the matrix *-**r**^-^*^1^ quantifies the response of each module to the individual 1% activation of every other module.

### SYNERGY BETWEEN DRUG TREATMENT AND ACTIVATED JURKAT-PD-1 CELLS

For synergy experiments, BLCA cells were plated in 96 well plates as in viability experiments above. BLCA cells were immediately treated with four to five concentrations between 2.5 and 10 µM SIS3, 75 nM and 250 nM JNK IX, 75 and 250 nM MBQ-167, 75 nM and 250 nM Pictilisib,2.5 µM and 15 µM SB525334, 0.25 and 2 µM Infigratinib, or 0.94 and 5.6 µM Lapatinib. All treatments were standardized to ensure 0.1% DMSO. After 24 hours, media containing drugs was refreshed, and non-activated or activated Jurkat-PD-1 cells were added at ratios of 5:1, 10:1, or 15:1 (Jurkat-PD-1: BLCA) for synergy quantification. After 3 days, Jurkat-PD-1 cells were carefully removed by pipetting, BLCA cells washed once with PBS, and visually inspected for complete removal of Jurkat-PD-1 cells prior to viability measurement by MTS assay. For PMA/Ionomycin experiments, five concentrations of PMA/Ionomycin between 0.1 and 1 nM based on an estimated residual PMA/Ionomycin concentration following Jurkat-PD-1 washing at 0.5 nM were added instead of Jurkat-PD-1 cells. All combinations were measured in duplicate for each experiment, and 3 independent experiments were conducted. Viability was quantified relative to DMSO control. The SynergyFinder tool was used to generate 2D synergy contour plots, quantify the mean Loewe synergy score, and generate p-values for significant synergy or antagonism as described previously^118^. Loewe Isoboles were generated with custom Python code.

### PRIMARY HUMAN CD4^+^ T CELL CULTURE

PBMCs from four healthy human donors were purchased from 2BScientific and VWR. CD4^+^ T cells were isolated using a CD4^+^ T cell isolation kit (Miltenyi) according to the manufacturer’s instructions. T cells were validated for CD4 expression by flow cytometry with an anti-CD4 antibody conjugated to FITC (Biolegend). T cell culture media was comprised of RPMI-1640 base media (Gibco), 1% L-glutamine (Gibco), 10% human serum (Merck), and 1% penicillin/streptomycin (Gibco). T cells were activated to proliferate by culturing together with Dynabeads^TM^ Human T-Activator CD3/CD28 beads (Thermo Fisher Scientific) in accordance with the manufacturer’s instructions. Ethical approval for use of PBMCs from anonymized healthy donors was received from UCD Research Ethics Committtee.

### PRIMARY CD4^+^ T CELL CYTOTOXICITY ASSAY

J82 cells were seeded at 15,000 cells per well and HT1376 cells at 30,000 cells per well of a 24-well plate. The cells were then treated with 10 µM SIS3, 100 nM JNK IX for J82 or 250 nM JNK IX for HT1376, or 0.1% DMSO for 24 hours. The drug concentrations used were the respective highest synergy drug concentration in the activated Jurkat-PD-1 synergy experiments. T cells were grown until they reached log growth phase, when the beads were removed and the cells further activated with 5 ng/ml PMA and 500 ng/ml ionomycin for 5 hours. They were then washed thrice in 1 ml culture media, centrifuging at 600g for 4 min each wash. Viable cells were counted and added to J82 and HT1376 cells at a ratio of 2:1 (CD4^+^ T cell: BLCA cell) and SIS3, JNK IX, or DMSO was refreshed. T cells and BLCA cells were co-cultured with drugs for 72 hours. T cells were harvested by pipetting, and BLCA cells harvested by trypsinization. The T cells and BLCA cells were stained together with 3,3’-Dioctadecyloxacarbocyanine Perchlorate (DiO) to exclude non-cellular debris, propidium iodide (PI) as a viability dye (Biolegend), and anti-CD45 conjugated to APC-H7 diluted 1:50 in 0.22 µM-filtered 3% FBS in PBS. The samples were run on a Cytoflex LX cytometer with the B525 filter for DiO, Y585 filter for PI, and R763 filter for APC-H7. Fluorescence Minus One (FMO) controls were used to determine gates for each fluorophore, and single-color controls were used for compensation. All flow data was analyzed using CytExpert software (Beckman Coulter). BLCA cell death as measured by DiO^+^CD45^low^PI^+^ percentage were quantified relative to DMSO controls. T cell death as measured by DiO^+^CD45^high^PI^+^ percentage were quantified relative to CD4^+^ T cell/BLCA cell co-culture condition.

Percentage of cancer cells was measured by DiO^+^CD45^-^PI^-^ events divided by the sum of DiO^+^CD45^-^PI^-^ and DiO^+^CD45^+^PI^-^ events. Between 2 and 4 replicates of each donor were averaged, and all are quantified as n=4 independent donors.

### CELL DEATH PATHWAY WESTERN BLOTS

J82 cells were seeded at 250,000 cells per well and HT1376 cells at 400,000 cells per well of a 6-well plate in triplicate for all conditions. J82 cells and HT1376 cells were treated for 24 hours with 10 µM SIS3, 100 nM JNK IX for J82 or 250 nM JNK IX for HT1376, 250 nM MBQ-167, or 0.1% DMSO. Drugs were then refreshed and activated Jurkat-PD-1 cells were added at a ratio of 7.5:1 (Jurkat-PD-1: J82) for J82 cells and 10:1 (Jurkat-PD-1: HT1376) for HT1376 cells. After 48 hours co-culture, Jurkat-PD-1 cells were removed by brief trypsinization and washing with ice-cold PBS. Jurkat-PD-1 cell removal was confirmed visually and using CD45 antibody staining by Western blot (data not shown). 125 µL ice-cold NP40 (20 mM Tris HCl pH 7.5, 150 mM NaCl, 0.5% NP40 with protease and phosphatase inhibitors (Roche)) was added to each well and cells collected with a cell scraper. Lysates were clarified by centrifugation at 16,000g for 10 minutes at 4°C. Protein concentration of lysates was determined by BCA assay according to the manufacturer’s instructions (Pierce^TM^ BCA Protein Assay Kit, Thermo Fisher Scientific), 10 µg of protein were resolved by SDS PAGE using 10% acrylamide gels in a Mini-PROTEAN Tetra cell system (Biorad). Proteins were then transferred on a PVDF membrane using the Xcell II^TM^ wet transfer system (Invitrogen). Membranes were blocked for 1 hour at room temperature using 4% BSA in TBS-Tween 20 0.1% (TBST). Membranes were then incubated with primary antibodies (Extended Data Table S6) diluted 1:1000 in blocking buffer from 2 hours at room temperature to overnight at 4°C. Membranes were then washed thrice in TBST and incubated with HRP-conjugated secondary antibodies diluted at 1:3000 in blocking buffer at room temperature for 1 hour. Blots were developed with homemade ECL reagent or SuperSignal^TM^ West Femto Reagent (Thermo Fisher Scientific) using an iBright^TM^ CL750 Imaging System (Thermo Fisher Scientific). Membranes were stripped using mild stripping buffer (15g glycine, 1g SDS, 10 mL Tween 20 in 1 mL water, pH 2.2). GAPDH was used as a loading control and was conducted for all membranes. Bands were quantified using the Gel tools in Fiji (ImageJ), and all values were quantified relative to GAPDH control bands on the same membrane followed by DMSO-only treatment for each cell line.

### LIPID PEROXIDATION ASSAY

J82 and HT1376 cells were seeded in a 24 well plate at 12.5 x 10^3^ and 2.5 x 10^3^ cells per well, respectively. The cells were immediately treated with 10 µM SIS3, 100 nM JNK IX for J82 or 250 nM JNK IX for HT1376, 250 nM MBQ-167, or 0.1% DMSO for 24 hours. Activated Jurkat-PD-1 cells were then added at an effector:target ratio of 10:1 and co-cultured for 48 hours. The Jurkat-PD-1 cells were then mechanically removed and BODIPY 581/591 C11 sensor dye (Thermo Fisher) added for 30 min in Jurkat media at 37°C. In its reduced form, the dye fluoresces red, whereas in its oxidized form it fluoresces green. Cells were washed thrice in PBS, trypsinized, and resuspended in 22 µM-filtered 3% FBS in PBS. The cells were then protected from light and immediately run on a Cytoflex LX flow cytometer. The mean fluorescence intensity from red (Texas Red filter) and green (FITC filter) fluorescence were measured by flow cytometry. Lipid peroxidation was quantified using the ratio of oxidized (green) to reduced (red) signal: a high ratio indicates high lipid peroxidation, while a low ratio indicates low lipid peroxidation.

### STATISTICS

Brown-Forsythe test and matched one-way ANOVA followed by Tukey’s correction were used for primary CD4^+^ T cell cytotoxicity assays and Western blot experiments. One-way ANOVA F statistics and degrees of freedom (DFs) are provided in Extended Data Table S7. Drug-Jurkat-PD-1 synergy experiments were analyzed for significant synergy and antagonism as described previously^118^. Viability experiments, HLA-DR expression, and JNK activation analyzed by two-tailed, two-sample t-test. Jurkat-PD-1 cell apoptosis and activation following drug treatment and BLCA cell co-culture was analyzed using two-way ANOVA. Normality was tested using a Shapiro-Wilk test and visualization on a QQ plot. All statistics were conducted in GraphPad Prism except for Loewe synergy scores, which were analyzed using SynergyFinder.

No statistical methods were used to predetermine sample size. The sample sizes were chosen based on established standards in the literature for in vitro viability (n = 3 independent biological replicates) and Western blot (n = 6 independent biological replicates) assays. This sample size was deemed sufficient to detect the observed large biological effect sizes with high reproducibility across independent runs. Individual data points are plotted to ensure full data transparency.

## DATA AVAILABILITY STATEMENT

All raw data generated in the study will become publicly available upon publication.

## CODE AVAILABILITY STATEMENT

The code used for the analysis will become publicly available upon publication.

## Supplementary Text

The cSTAR predictions of immunosensitization in the immunoresistant cell lines J82 and HT1376 (Fig. 4) were derived from cell-specific network responses inferred using STV_imres_, which was constructed by separating the basal-state proteomic patterns of the immunosensitive cell lines HCV-29 and RT4 from those of the immunoresistant cell lines J82 and HT1376 (Fig. 3). Although main predictions that used the STV_imres_ were corroborated experimentally, some co-culture experiments in the immunosensitive non-cancerous cell line HCV-29 yielded results that differed from the model’s predictions. Specifically, whereas the model predicted that the SMAD3 pathway did not contribute to immunoresistance in HCV-29 and ERBB1/2 signaling negatively contributes to immunoresistance, inhibition of SMAD3 with SIS3 and ERBB1/2 with Lapatinib sensitizes HCV-29 cells to activated Jurkat-PD-1 cytotoxicity, although the overall synergy is not significant (Extended Data Fig. 10b). Interestingly, we found that HCV-29 cells de-activate and induce apoptosis in Jurkat-PD-1 cells following their activation by PMA/ionomycin (Extended Data Fig. 10c–e). This suggests that SIS3 and Lapatinib cause cell death in HCV-29 cells, and therefore decrease Jurkat-PD-1 cell death due to HCV-29 cell co-culture, which potentiates activated Jurkat-PD-1 cytotoxicity in HCV-29.

To account for residual PMA/Ionomycin after Jurkat-PD-1 activation as a potential confounding factor in the experimental validations of cSTAR predictions, we first added the final wash following the Jurkat-PD-1 activation procedure to BLCA cells and assessed its effect on cell viability. While all cells except HT1376 experienced slight decreases in viability, only BC-3C was statistically significant (Extended Data Fig. 11a). We then used a PMA/ionomycin dose-response curve (Extended Data Fig. 11b) to obtain a conservative estimate of the residual concentration of PMA/ionomycin in the final wash at approximately 0.5 nM, although the actual concentration used in activated Jurkat-PD-1 cytotoxicity experiments is likely between 2- and 5-fold lower than 0.5 nM due to sample dilution. Loewe synergy between PMA/Ionomycin at concentrations above and below 0.5 nM and SIS3 at the doses used in drug and activated Jurkat-PD-1 synergy experiments was quantified using SynergyFinder. Although the non-cancerous HCV-29 cell line showed significant synergy (Extended Data Fig 11c), albeit at concentrations of PMA/Ionomycin that are higher than what we estimate is present in culture after Jurkat-PD-1 activation and washing (between 0.5 and 1 nM, Extended Data Fig 11d), none of the cancer cell lines (J82, HT1376, BC-3C) showed measurable synergy (Extended Data Fig. 11c,d). This suggests that the synergy observed between SIS3 and activated Jurkat-PD-1 cells in the immunoresistant cell lines is primarily due to sensitization of BLCA cells to the cytotoxic activity of the activated Jurkat-PD-1 cells, and not residual PMA/Ionomycin effect on the BLCA cells.

In summary, discrepancies between cSTAR predictions and experimental validation can largely be explained by detrimental effects of FGFR inhibition and HCV-29 cells on Jurkat-PD-1 cell activity and viability. Additionally, the combination of SIS3 and residual PMA/Ionomycin is synergistic only in HCV-29, suggesting that observed synergy between drug treatments and activated Jurkat-PD-1 cells may be at least partially due to residual PMA/Ionomycin in HCV-29. However, we confirmed that residual PMA/Ionomycin is not a contributing factor in the synergy between SIS3 and activated Jurkat-PD-1 cells in the immunoresistant cell lines J82, HT1376, and BC-3C.

## ACKNOWLEDGEMENTS

The authors wish to acknowledge Günther Eissner and Myriam Nabhan for their discussions on cancer immunology. We also thank Lars Dyrskjøt and Peter Steinberger for providing many of the cell lines used in this study. The results published here are in part based upon data generated by The Cancer Genome Atlas (TCGA) Research Network: https://www.cancer.gov/tcga.

## AUTHOR CONTRIBUTIONS

O.S.R., W.K., S.Y.T.R, and B.N.K. conceived the study. S.Y.T.R, N.R., and T.S. conducted wet lab experiments. S.Y.T.R. analyzed wet lab experiments. J.Y. and O.S.R. performed bioinformatics and cSTAR analysis. K.W. performed mass spectrometry proteomics. A.M. and N.O.C. conducted RPPA experiments and data processing. S.Y.T.R., O.S.R., W.K. and B.N.K. wrote the manuscript.

## SUPPLEMENTARY INFORMATION

Supplementary information is available for this paper.

## MATERIALS AND CORRESPONDENCE

Correspondence and requests for materials should be addressed to

Reprints and permissions information is available at www.nature.com/reprints.

## EXTENDED DATA FIGURE LEGENDS

**Extended Data Figure 1.**
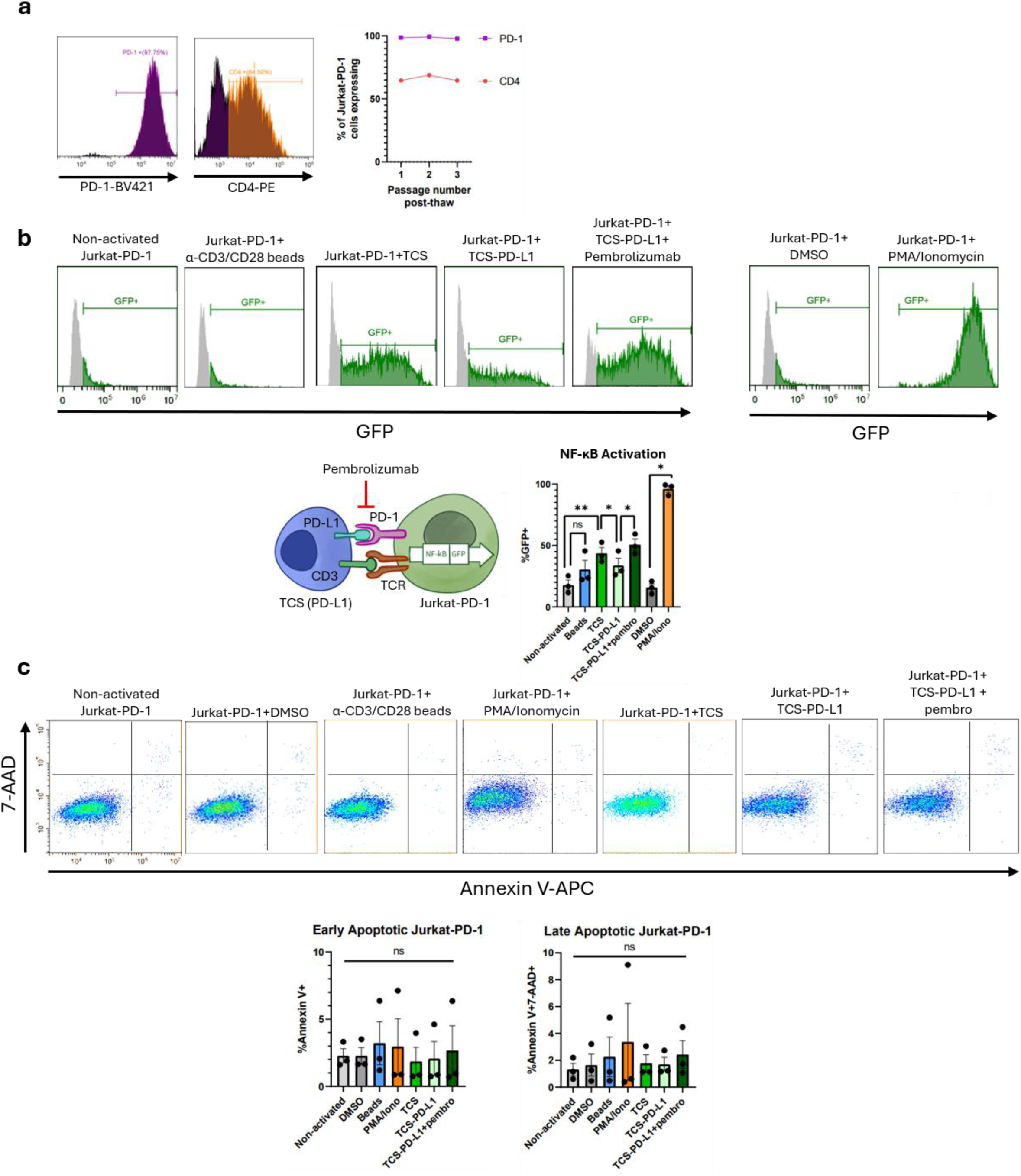
CD4 and PD-1 expression, NF-κB activation and apoptosis in Jurkat reporter cells for different activation methods. **a**. Percentages of Jurkat-PD-1 cells expressing PD-1 (purple) and CD4 (orange) measured by flow cytometry over three passages post-thaw. **b.** GFP reporter expression indicating NF-κB activation levels in Jurkat-PD-1 cells following activation by α-CD3/CD28 beads, 100 nM each PMA/Ionomycin, TCS cells cultured at 1:4 ratio TCS:Jurkat-PD-1, or TCS-PD-L1 cells cultured at a 1:4 ratio TCS-PD-L1:Jurkat-PD-1 with or without the immune checkpoint inhibitor pembrolizumab. **c.** Apoptosis of Jurkat-PD-1 cells following various activation methods quantified by Annexin V and 7-AAD staining in flow cytometry assays. Bottom right quadrant indicates early apoptotic (Annexin V+ alone) cells, and top right quadrant indicates late apoptotic (Annexin V+ 7AAD+) cells. n=3-4 independent experiments. *p<0.05, **p<0.01 by matched one-way ANOVA followed by Tukey’s correction.

**Extended Data Figure 2.**
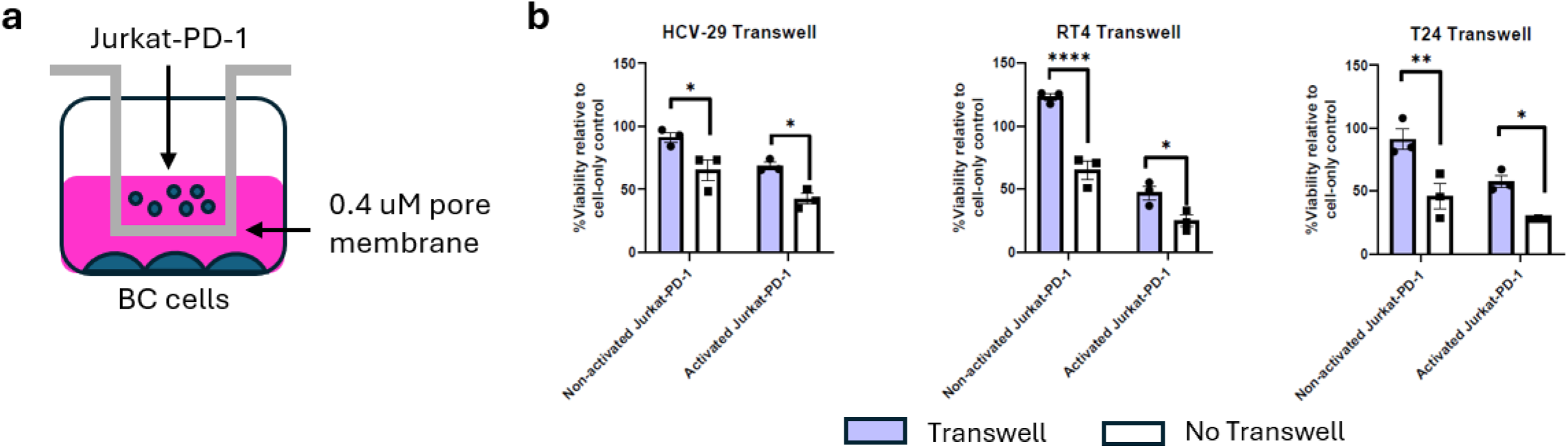
Jurkat-PD-1 cytotoxicity against immunosensitive BLCA cells is enhanced by direct cell–cell contact. **a**. Schematic of the Transwell experimental setup. Jurkat-PD-1 cells were placed in the upper chamber, allowing diffusion of soluble factors through the membrane pores while preventing direct cell–cell contact. **b**. Viability of BLCA cells following 3-day co-culture with Jurkat-PD-1 cells in the Transwell system, measured by MTS assay. Viability in the Transwell system was compared with that measured in co-culture experiments without the Transwell system (Figure 1). n=3 independent experiments. *p<0.05, **p<0.01, ***p<0.001, ****p<0.0001 by two-way ANOVA.

**Extended Data Figure 3.**
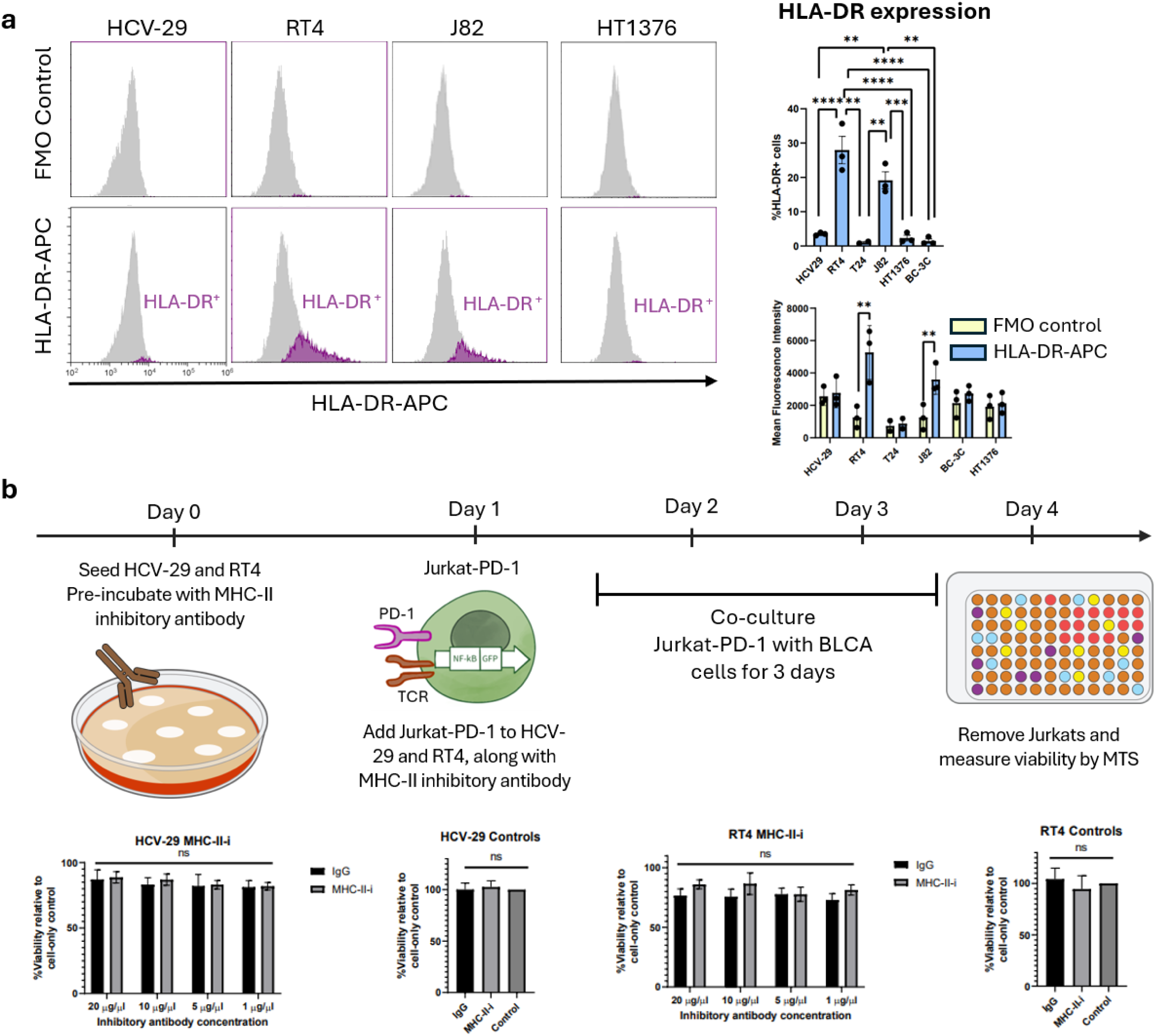
MHC class II (HLA-DR) is expressed on RT4 and J82 cells but does not influence Jurkat-PD-1 cytotoxicity. **a.** Surface expression of HLA-DR (MHC class II) was quantified by flow cytometry in live (7-AAD–negative) BLCA cells using an antibody specific for HLA-DR conjugated to the APC fluorophore. Fluorescence minus one (FMO) control comprised BLCA cells stained with 7-AAD alone but not with the HLA-DR antibody. Percentage of HLA-DR^+^ cells and mean fluorescence intensity (MFI) are quantified in the right panel. **b.** MHC class II was inhibited using a blocking antibody (MHC-II-i) in immunosensitive BLCA cells prior to and simultaneously with non-activated Jurkat-PD-1 cells. Jurkat-PD-1 cells were removed mechanically, and viability was measured by MTS. n=3 independent experiments. *p<0.05, **p<0.01, ***p<0.001, ****p<0.0001 by one-way ANOVA followed by Tukey’s correction.

**Extended Data Figure 4.**
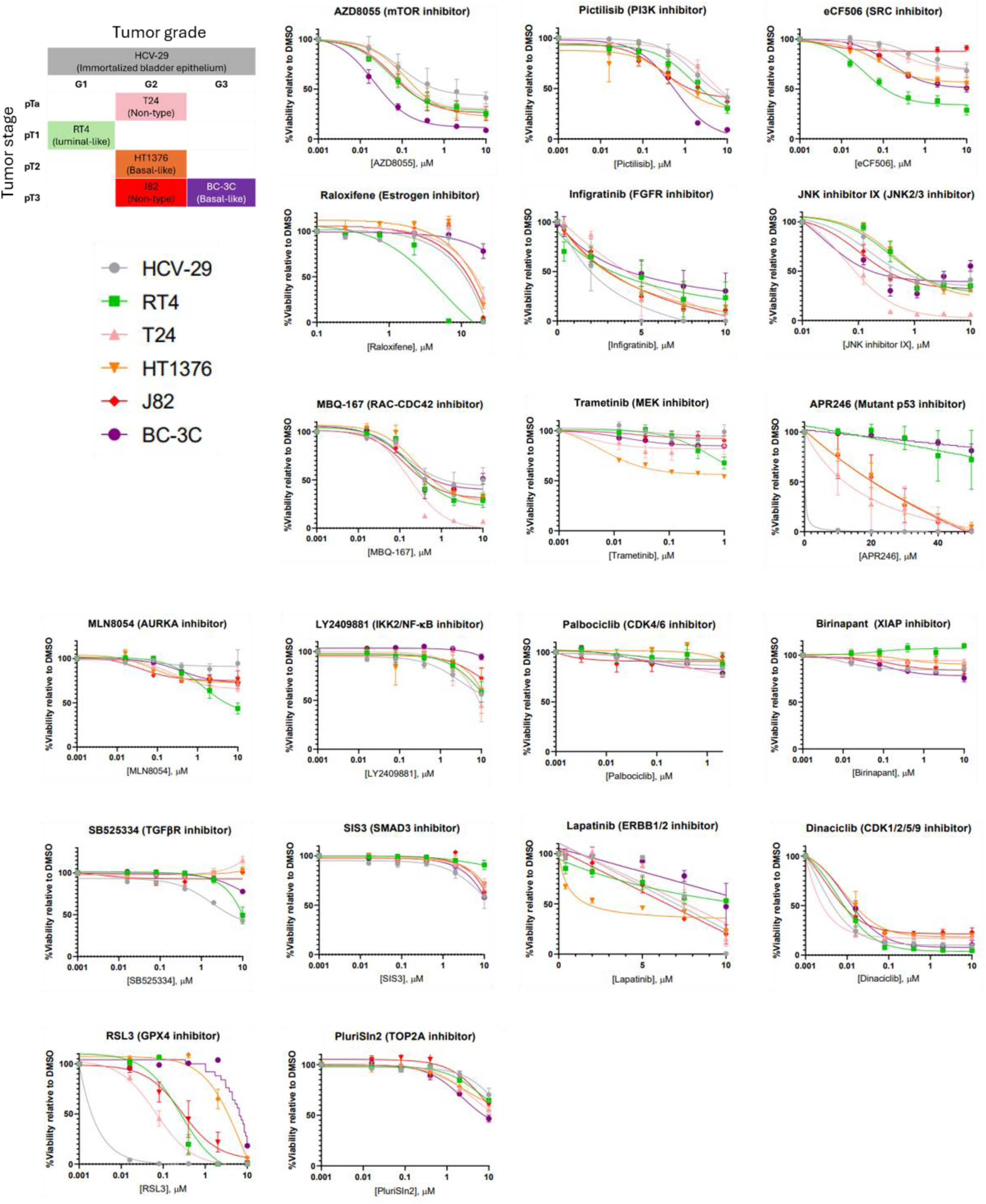
Full MTS drug screen data. The tumor grade, tumor stage, and subtype of each cell line is listed in the upper left. All cell lines were treated for 72 h using the drugs and concentrations listed in Extended Data Table S1. Viability was measured using MTS assay pre- and 72 h post-drug treatment. Graph curves are non-linear regression conducted using GraphPad Prism. n=3-4 independent experiments.

**Extended Data Figure 5.**
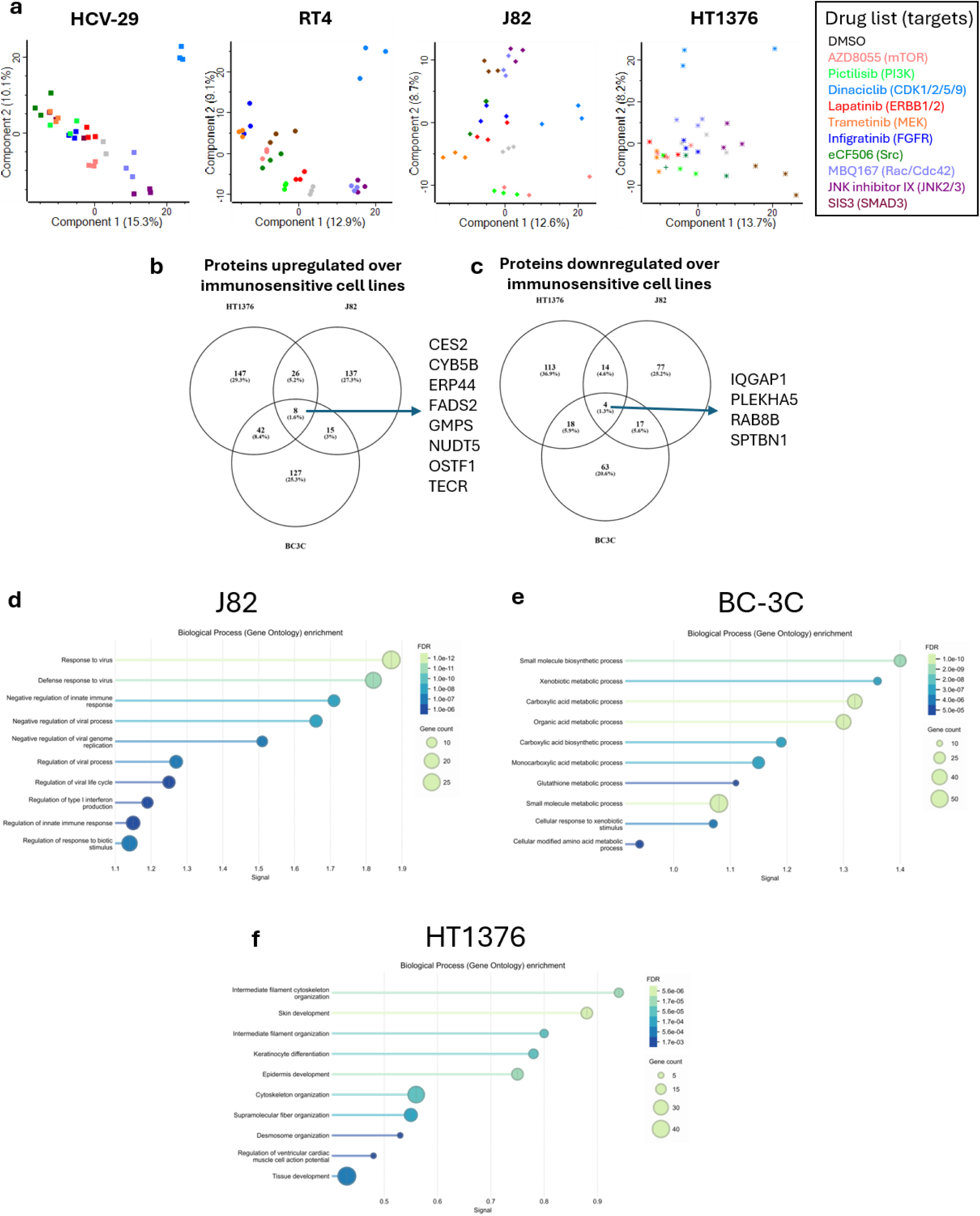
Proteomic signatures of immunoresistant (J82, BC-3C, and HT1376) cell lines versus immunosensitive (HCV-29, RT4, T24) cell lines. **a.** PCA plots of HCV-29, RT4, J82, and HT1376 cells treated with 10 different drugs and DMSO control for 24 h. Drug concentrations are listed in Extended Data Table S1. Protein expression is quantified using proteomics. Proteomic quantification of proteins upregulated (**b**) and downregulated (**c**) in unperturbed immunoresistant cell lines (J82, HT1376, BC-3C) relative to unperturbed immunosensitive cell lines (HCV-29, RT4, T24). **d–f.** Gene ontology enrichment analysis of pathways in J82 (**d**), BC-3C (**e**), and HT1376 (**f**) relative to immunosensitive cells (HCV-29, RT4, and T24).

**Extended Data Figure 6.**
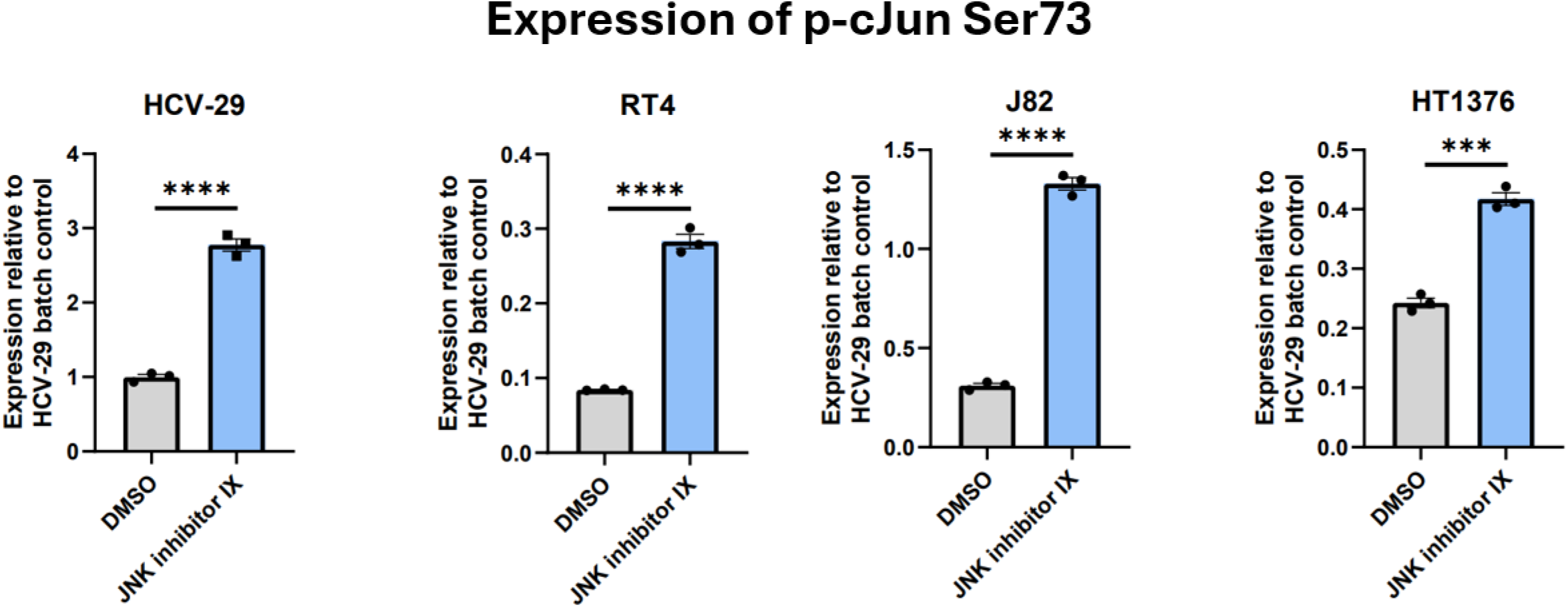
JNK inhibitor IX paradoxically activates the JNK pathway. Phosphorylated cJun Ser73, a marker of JNK activation, was quantified by RPPA following JNK inhibitor IX treatment or DMSO control. In all cell lines measured, phosphorylated cJun Ser73 increased following 24 h treatment with JNK inhibitor IX, indicating paradoxical activation of the JNK pathway. n=3 replicates, ****p<0.0001 by two-sample t-test.

**Extended Data Figure 7:**
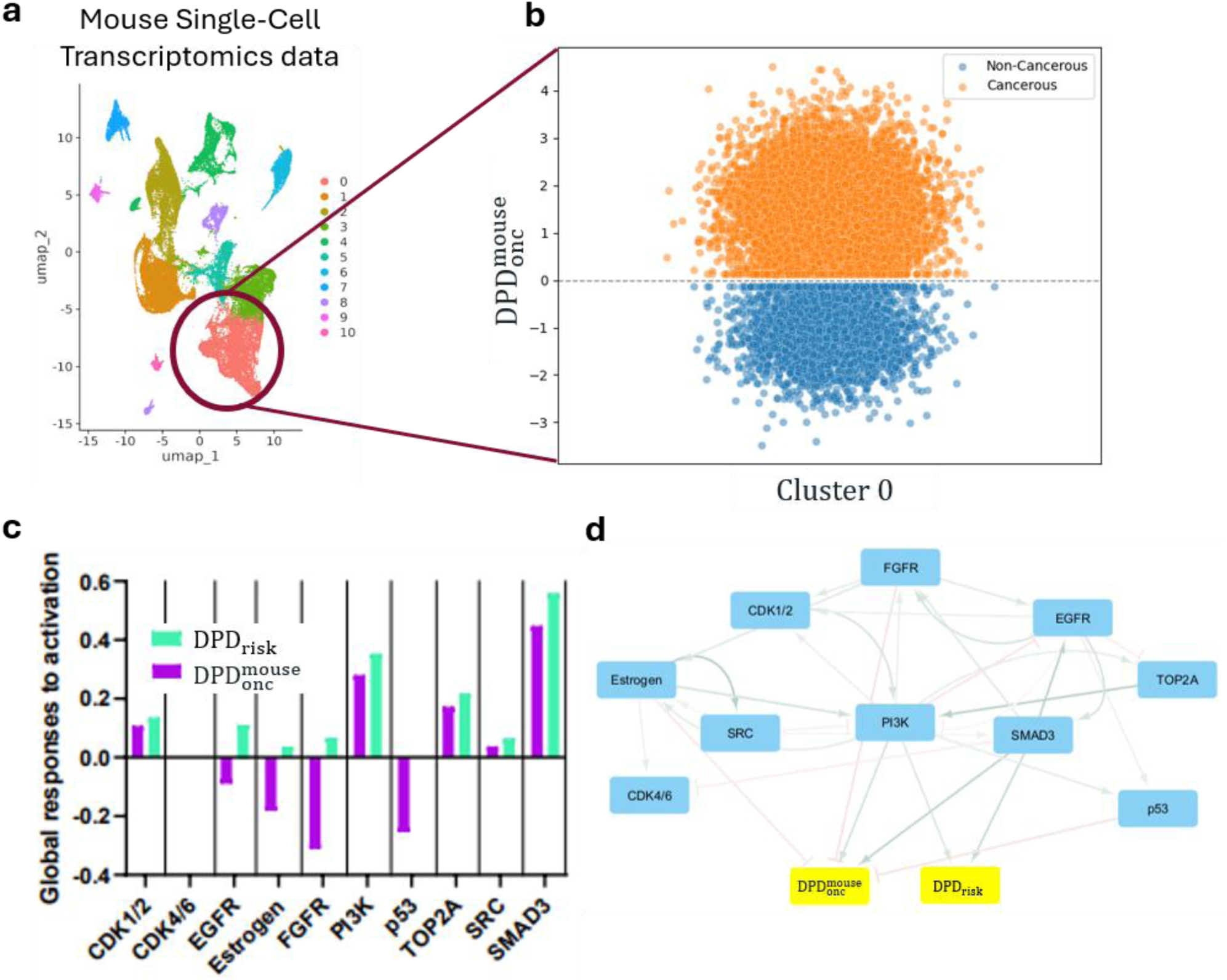
A DPD derived from mouse single-cell RNA-seq data recapitulates the major findings obtained using human-derived DPDs. **a.** Single-cell RNA-seq data of whole bladders from a semi-spontaneous, carcinogen-induced (OH-BBN) mouse model of urothelial cancer (Hgf-Cdk4^R24C^ mice) at the non-muscle-invasive and muscle-invasive stages, and from healthy control bladders, were obtained from GEO (accession number GSE174182). The epithelial-like cell cluster (cluster 0, circled in red) was selected for downstream analysis. **b.** 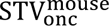 was generated by separation of non-cancerous and cancerous cells in the single-cell RNA-seq space. The scatterplot shows the separation of cancerous (orange) and non-cancerous (blue) cells by their corresponding 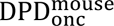 scores. **c.** Global systems-level effects of pathways perturbed in BC-3C cells on 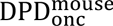. RNA-seq data was acquired from the LINCS database (transcriptomic responses of the CDK1/2, CDK4/6, ERBB1/2, Estrogen, FGFR, PI3K, p53, TOP2A, and SRC pathways to pharmacological perturbations) and integrated with in-house generated RNA-seq data for SMAD3 inhibition. **d.** 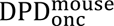 was merged into the BC-3C network from Fig. 4 and added as an additional inferred DPD node (yellow boxes). Local pathway effects are represented by green arrows for positive direct connections and red blocked arrows for negative direct connections.

**Extended Data Figure 8.**
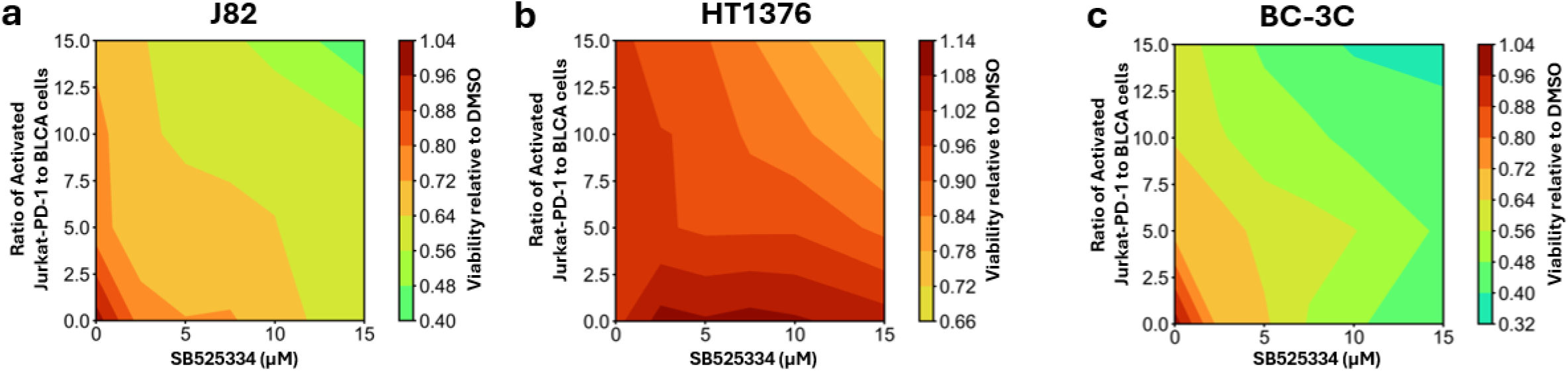
TGFβR inhibition does not substantially increase immunosensitivity in immunoresistant BLCA cell lines. J82 cells (**a**), HT1376 cells (**b**), and BC-3C cells (**c**) were pre-treated with TGFβR inhibitor SB525334 for 24 h, followed by co-culture with PMA/Ionomycin-activated Jurkat-PD-1 for 72 h. Viability of BLCA cells was measured by MTS after 72 h, and synergy between SB525334 and activated Jurkat-PD-1 was observed using Loewe isoboles where convex curves indicating synergy and concave curves indicating antagonism. Loewe isoboles represent an averaged summary of n=3 independent experiments.

**Extended Data Figure 9.**
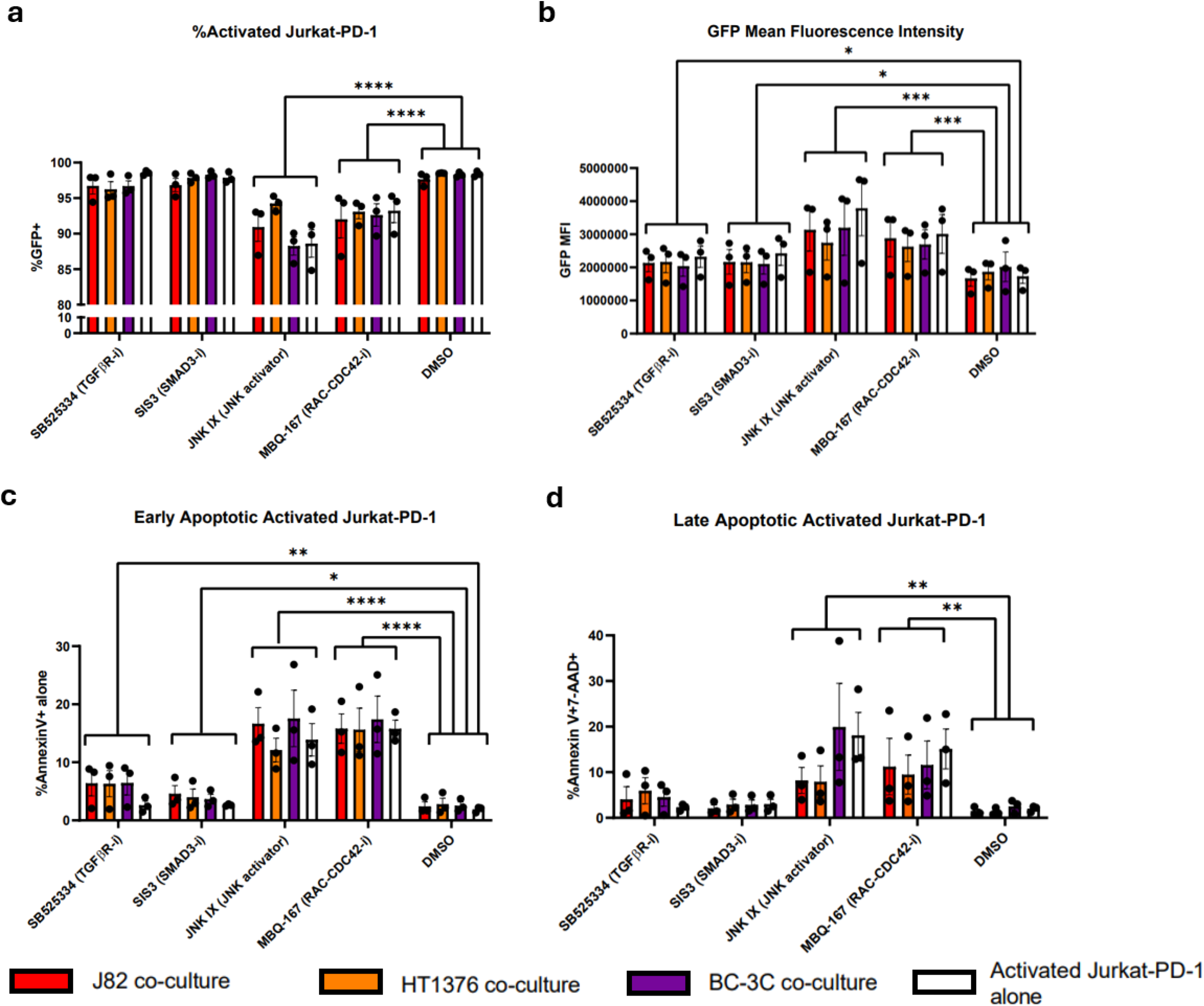
All drugs tested modulate Jurkat-PD-1 activation, while JNK IX and MBQ-167 induce apoptosis in activated Jurkat-PD-1 cells. Immunoresistant J82, HT1376, or BC-3C cells were pre-treated for 24 hours with 10 µM SIS3, 15 µM SB525334, 100 nM JNK IX (J82) or 250 nM JNK IX (HT1376), 250 nM MBQ-167, or DMSO control. Following pre-treatment, drugs were refreshed and activated Jurkat-PD-1 cells were added at an effector:target ratio of 10:1. The effects of each drug and BLCA cell line on activated Jurkat-PD-1 cell activation and apoptosis was quantified using flow cytometry. **a.** The percentage of activated Jurkat-PD-1 cells was quantified using the percentage of GFP⁺ cells in flow cytometry. **b.** The overall activation status of the Jurkat-PD-1 cells was measured using GFP mean fluorescence intensity in flow cytometry. **c-d.** Quantification of early apoptosis (**c,** Annexin V⁺ only) and late apoptosis (**d,** Annexin V⁺/7-AAD⁺) in activated Jurkat-PD-1 cells by flow cytometry. n=3 independent experiments, *p<0.05, **p<0.01, ***p<0.001 using two-way ANOVA.

**Extended Data Figure 10.**
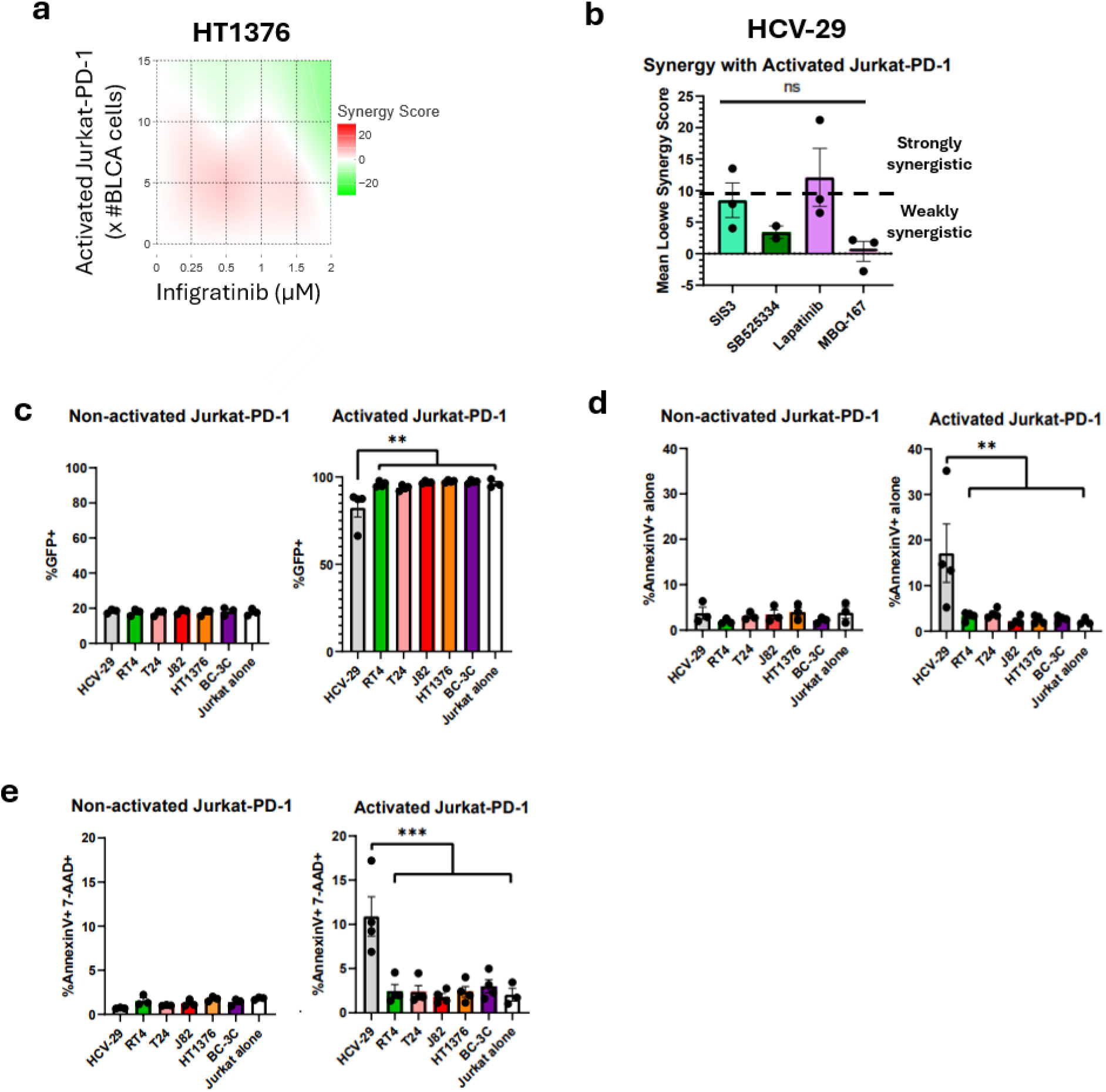
Experimental results indicating discrepancies between cSTAR predictions and observations. **a**. 2D contour plot of Loewe synergy scores for combinations of the FGFR inhibitor Infigratinib (0–2 µM) and activated Jurkat-PD-1 cells at effector-to-target (E) ratios of 15, 10 and 5 in HT1376 cells. High synergy score (red) indicates synergy, while low synergy score (green) indicates antagonism. Loewe synergy scores and 2D contour plot generated using the SynergyFinder tool based on cell viability relative to DMSO. **b**. Mean Loewe synergy scores calculated using SynergyFinder for combinations of activated Jurkat-PD-1 cells and inhibitors of SMAD3 (SIS3), TGFβR (SB525334), ERBB1/2 (Lapatinib) or RAC–CDC42 (MBQ-167), at E ratios of 15, 10 and 5. Synergy scores were calculated based on the viability of HCV-29 cells following treatment with the respective drug combinations. Loewe synergy scores greater than 10 indicate strong synergy. **c**. Flow cytometry quantification of GFP as a readout for NF-κB activation of Jurkat-PD-1 cells following 72-hour co-culture with BLCA cells and control HCV-29 cells. **d-e**. Apoptosis in Jurkat-PD-1 cells following co-culture with BLCA cells and non-cancerous HCV-29 control cells was measured using AnnexinV and 7-AAD staining in flow cytometry. Early apoptotic cells are measured by AnnexinV staining alone (**d**), while late apoptotic cells are measured by AnnexinV and 7-AAD double positivity (**e**). n=3 independent experiments, *p<0.05, **p<0.01, ***p<0.001 using one-way ANOVA.

**Extended Data Figure 11.**
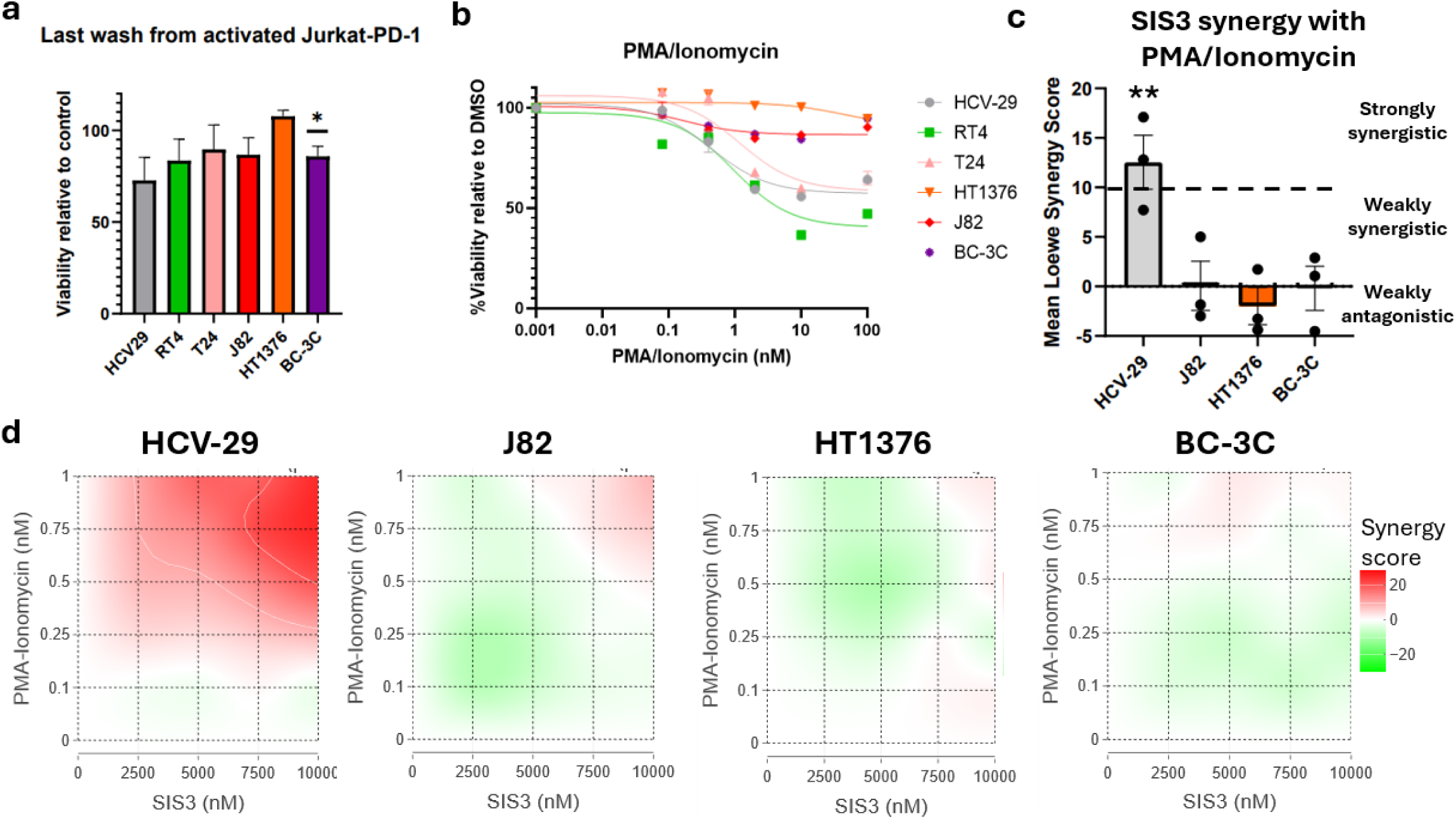
SMAD3 inhibition does not sensitize BLCA cells but increases the sensitivity of control HCV-29 cells to cytotoxicity induced by residual PMA/Ionomycin. **a.** To evaluate the effects of residual PMA/Ionomycin on BLCA cells and control HCV-29 cells, the Jurkat-PD-1 cells were activated with 100 nM each of PMA and ionomycin, washed thrice in FBS-containing media, and the final wash was directly added to the target cells. Cell viability was measured by MTS after 72 h. *p<0.05 compared to DMSO control by paired two-sample t-test. **b.** Dose-response curves of PMA/Ionomycin effect on the viability of all cell lines were used to estimate the quantity of residual PMA/Ionomycin in the final wash from panel **a**, which is estimated around 0.5 nM. **c.** The mean Loewe synergy scores of pre-incubated SIS3 (between 0 and 10 µM) and PMA/Ionomycin (between 0 and 1 nM) was calculated using SynergyFinder. ***p<0.001 using SynergyFinder. **d**. 2D contour maps of Loewe synergy at each drug combination for all four cell lines used for measuring synergy between SIS3 and Jurkat-PD-1-induced cytotoxicity. Red areas represent regions of synergy, while green areas represent regions of antagonism. n=3 independent experiments.

**Extended Data Figure 12.**
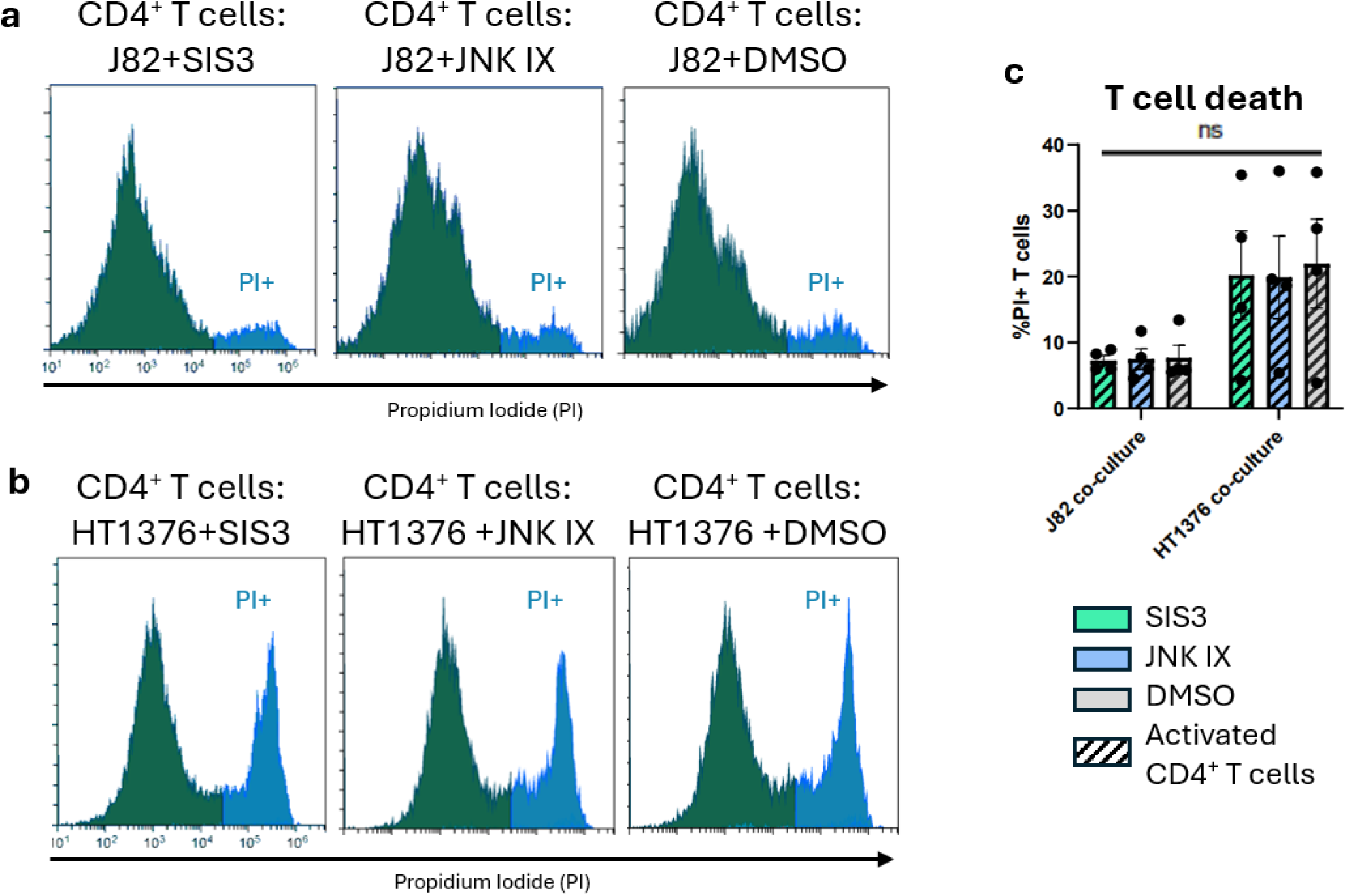
Neither SMAD3 inhibition by SIS3 nor JNK activation by JNK IX affects CD4⁺ T cell death. Propidium iodide (PI)–based flow cytometry of CD4⁺ T cells following co-culture with J82 cells (**a**) and HT1376 cells (**b**). Flow traces are 1 representative experiment of n=4 independent donors. BLCA cells were separated from CD4^+^ T cells using CD45 staining in flow cytometry. c. Quantification of dead CD4^+^ T cells following co-culture using the percentage of PI^+^ cells. Each datapoint in the bar graph represents a single donor, which is an average of 2-4 technical replicates using each donor.

**Extended Data Figure 13.**
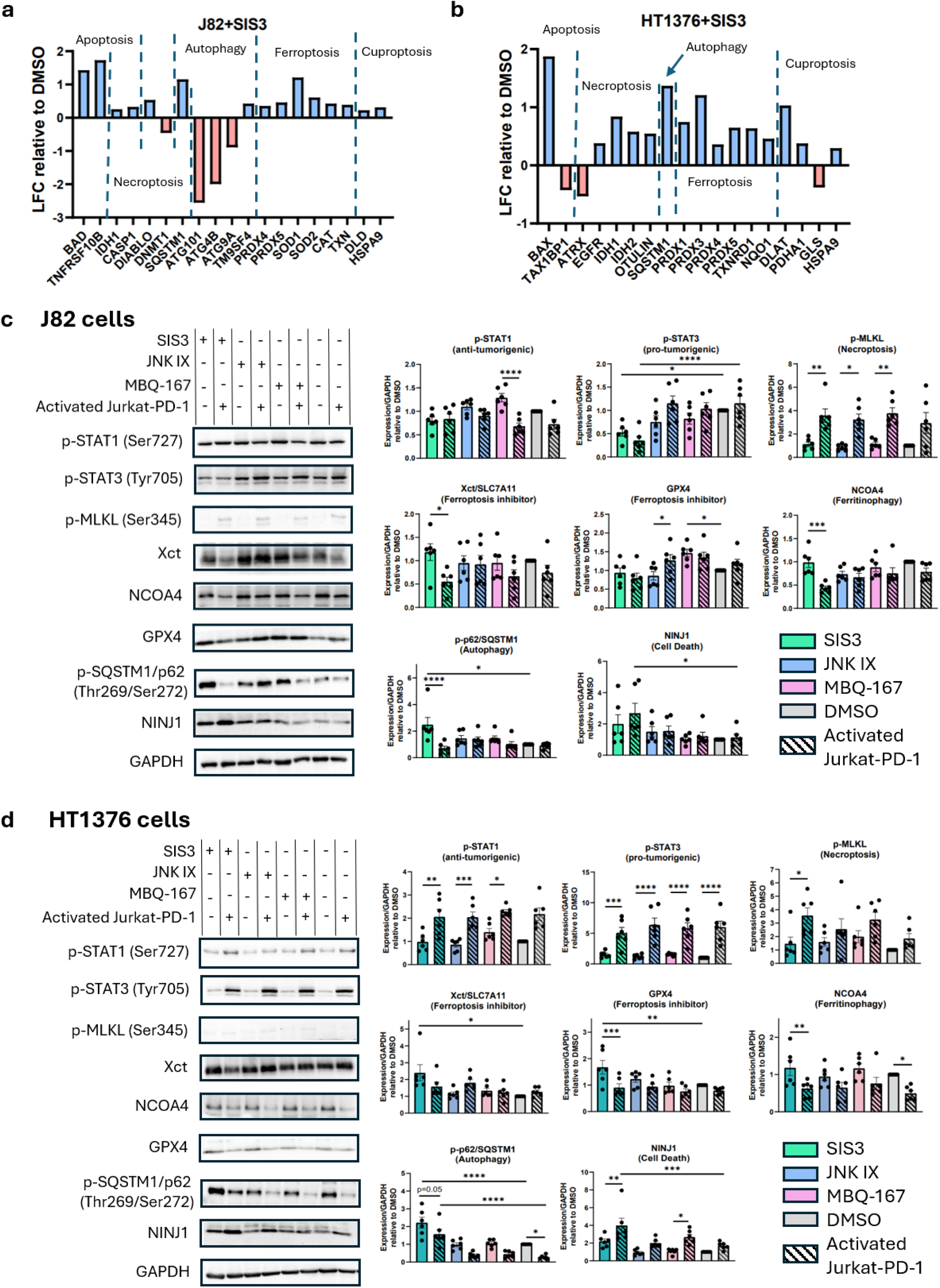
SIS3 in combination with activated Jurkat-PD-1 cells modulates cell death mechanisms in BLCA cells. **a-b.** Fold changes in proteins associated with apoptosis, necroptosis, autophagy, ferroptosis and cuproptosis in J82 (**a**) and HT1376 (**b**) cells following 24 h of SIS3 treatment, quantified by proteomics relative to the respective DMSO controls. Only proteins showing statistically significant changes are presented.c**–d.** Further characterization of cytotoxicity mechanisms by immunoblotting of selected markers (Extended Data Table S6) of the STAT, ferroptosis, ferritinophagy, and other cell death-related pathways following 24 h of pre-incubation with drugs and 48 h of co-culture with activated Jurkat-PD-1 cells and drugs together. n=6 biological replicates from 2 independent experiments, *p<0.05, **p<0.01, ***p<0.001, ****p<0.0001 by one-way ANOVA followed by Tukey’s correction, where bars show statistically significant pairs. Western blot images are from 1 representative replicate.

**Extended Data Figure 14.**
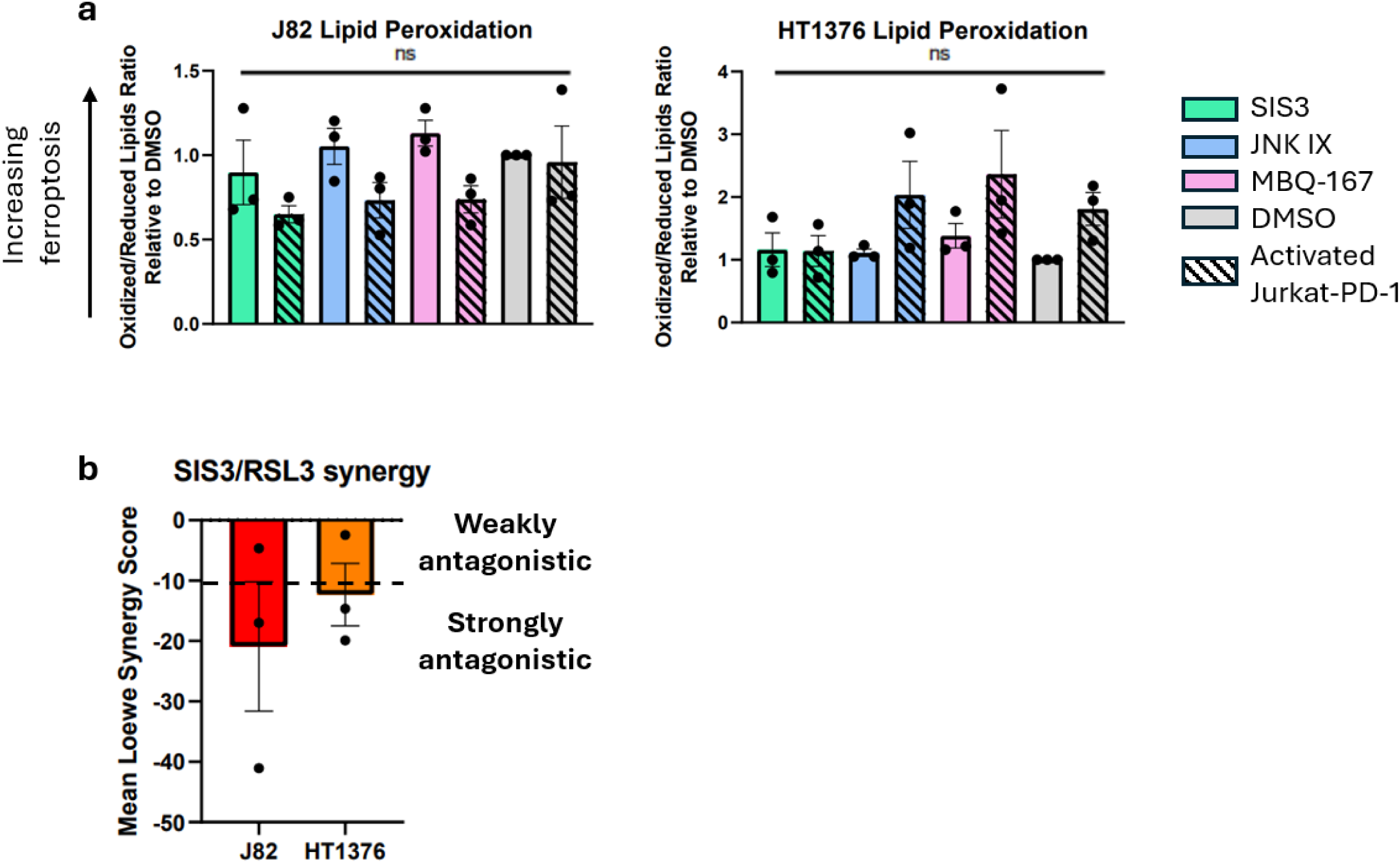
Effects of SMAD3 inhibition on lipid peroxidation and ferroptosis sensitivity in J82 and HT1376 cells. **a**. Lipid peroxidation in J82 and HT1376 cells was quantified by flow cytometry using the BODIPY sensor dye following drug treatment and co-culture with activated Jurkat-PD-1 cells. The reduced form of the dye fluoresces red, whereas the oxidized form fluoresces green. Mean fluorescence intensities were measured using the Texas Red and green FITC filters, respectively. The ratio of oxidized (green) to reduced (red) signal serves as a readout of lipid peroxidation: a high ratio indicates high lipid peroxidation, while a low ratio indicates low lipid peroxidation. **b**. Drug synergy between SIS3 (SMAD3 inhibitor) and RSL3 (GPX4 inhibitor) was evaluated by MTS assay and Loewe synergy was calculated by SynergyFinder. A negative Loewe synergy score indicates antagonism, and a Loewe synergy score <-10 indicates strong antagonism. n=3 independent experiments. One-way ANOVA followed by Tukey’s correction was conducted with no significant differences identified.

**Extended Data Table S1:**
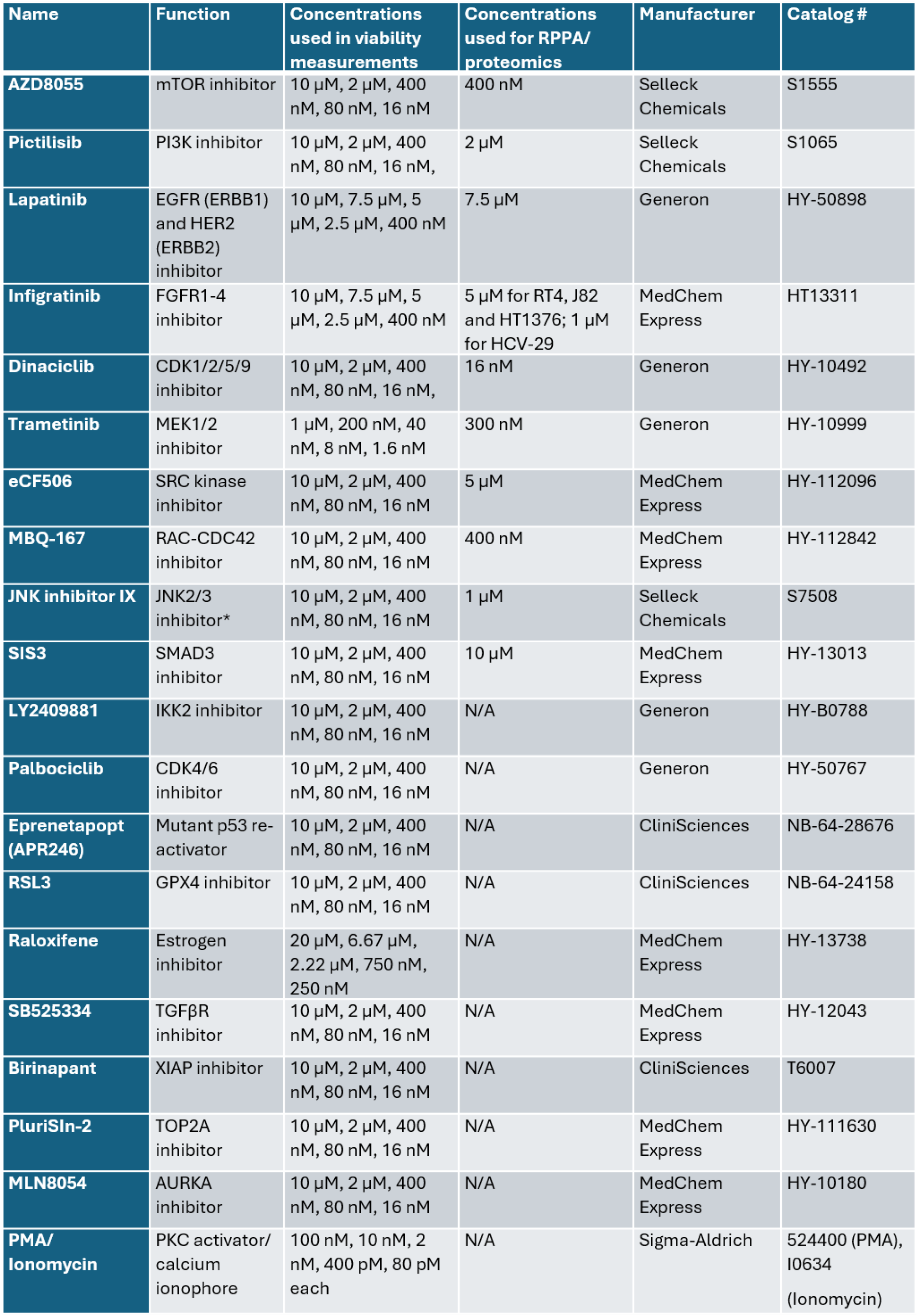
Chemicals used in this study.

**Extended Data Table S2:**
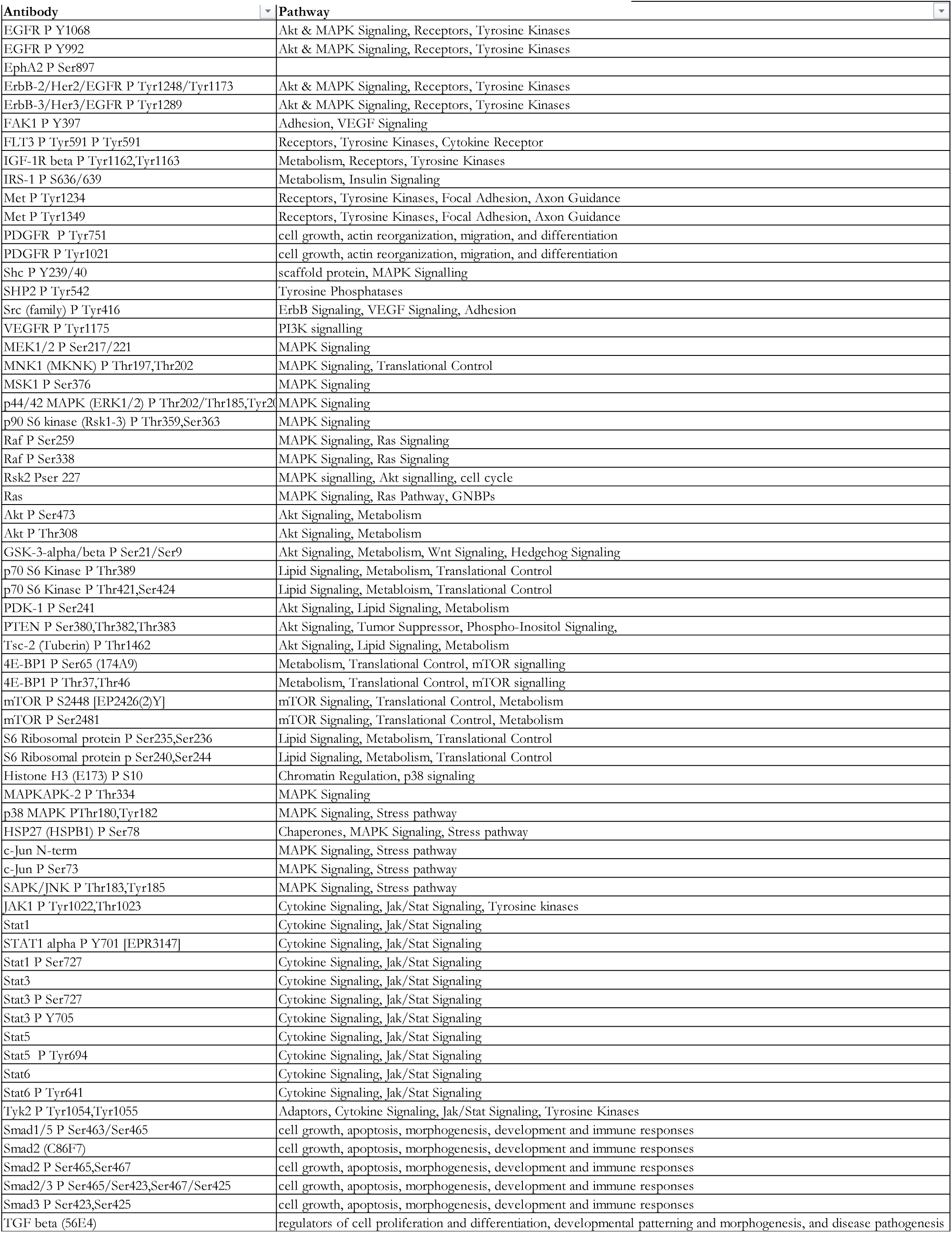
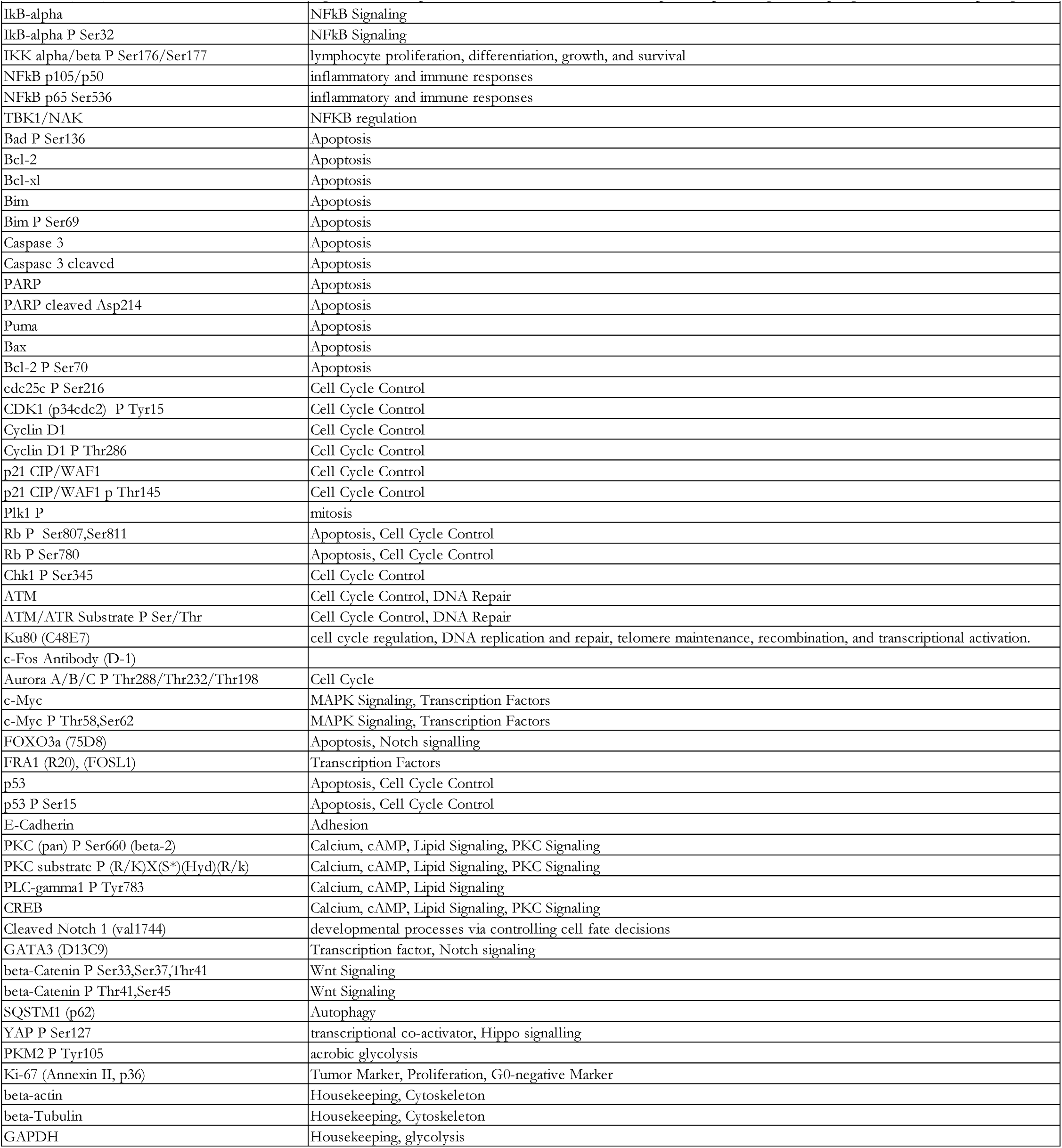
Analytes targeted in RPPA.

Analytes targeted in RPPA
| Antibody | Pathway |
| --- | --- |
| EGFR P Y1068 | Akt & MAPK Signaling, Receptors, Tyrosine Kinases |
| EGFR P Y992 | Akt & MAPK Signaling, Receptors, Tyrosine Kinases |
| EphA2 P Ser897 |  |
| ErbB-2/Her2/EGFR P Tyr1248/Tyr1173 | Akt & MAPK Signaling, Receptors, Tyrosine Kinases |
| ErbB-3/Her3/EGFR P Tyr1289 | Akt & MAPK Signaling, Receptors, Tyrosine Kinases |
| FAK1 P Y397 | Adhesion, VEGF Signaling |
| FLT3 P Tyr591 P Tyr591 | Receptors, Tyrosine Kinases, Cytokine Receptor |
| IGF-1R beta P Tyr1162,Tyr1163 | Metabolism, Receptors, Tyrosine Kinases |
| IRS-1 P S636/639 | Metabolism, Insulin Signaling |
| Met P Tyr1234 | Receptors, Tyrosine Kinases, Focal Adhesion, Axon Guidance |
| Met P Tyr1349 | Receptors, Tyrosine Kinases, Focal Adhesion, Axon Guidance |
| PDGFR P Tyr751 | cell growth, actin reorganization, migration, and differentiation |
| PDGFR P Tyr1021 | cell growth, actin reorganization, migration, and differentiation |
| Shc P Y239/40 | scaffold protein, MAPK Signalling |
| SHP2 P Tyr542 | Tyrosine Phosphatases |
| Src (family) P Tyr416 | ErbB Signaling, VEGF Signaling, Adhesion |
| VEGFR P Tyr1175 | PI3K signalling |
| MEK1/2 P Ser217/221 | MAPK Signaling |
| MNK1 (MKNK) P Thr197,Thr202 | MAPK Signaling, Translational Control |
| MSK1 P Ser376 | MAPK Signaling |
| p44/42 MAPK (ERK1/2) P Thr202/Thr185,Tyr204 | MAPK Signaling |
| p90 S6 kinase (Rsk1-3) P Thr359,Ser363 | MAPK Signaling |
| Raf P Ser259 | MAPK Signaling, Ras Signaling |
| Raf P Ser338 | MAPK Signaling, Ras Signaling |
| Rsk2 Pser 227 | MAPK signalling, Akt signalling, cell cycle |
| Ras | MAPK Signaling, Ras Pathway, GNBPs |
| Akt P Ser473 | Akt Signaling, Metabolism |
| Akt P Thr308 | Akt Signaling, Metabolism |
| GSK-3-alpha/beta P Ser21/Ser9 | Akt Signaling, Metabolism, Wnt Signaling, Hedgehog Signaling |
| p70 S6 Kinase P Thr389 | Lipid Signaling, Metabolism, Translational Control |
| p70 S6 Kinase P Thr421,Ser424 | Lipid Signaling, Metabolism, Translational Control |
| PDK-1 P Ser241 | Akt Signaling, Lipid Signaling, Metabolism |
| PTEN P Ser380,Thr382,Thr383 | Akt Signaling, Tumor Suppressor, Phospho-Inositol Signaling, |
| Tsc-2 (Tuberin) P Thr1462 | Akt Signaling, Lipid Signaling, Metabolism |
| 4E-BP1 P Ser65 (174A9) | Metabolism, Translational Control, mTOR signalling |
| 4E-BP1 P Thr37,Thr46 | Metabolism, Translational Control, mTOR signalling |
| mTOR P S2448 [EP2426(2)Y] | mTOR Signaling, Translational Control, Metabolism |
| mTOR P Ser2481 | mTOR Signaling, Translational Control, Metabolism |
| S6 Ribosomal protein P Ser235,Ser236 | Lipid Signaling, Metabolism, Translational Control |
| S6 Ribosomal protein p Ser240,Ser244 | Lipid Signaling, Metabolism, Translational Control |
| Histone H3 (E173) P S10 | Chromatin Regulation, p38 signaling |
| MAPKAPK-2 P Thr334 | MAPK Signaling |
| p38 MAPK PThr180,Tyr182 | MAPK Signaling, Stress pathway |
| HSP27 (HSPB1) P Ser78 | Chaperones, MAPK Signaling, Stress pathway |
| c-Jun N-term | MAPK Signaling, Stress pathway |
| c-Jun P Ser73 | MAPK Signaling, Stress pathway |
| SAPK/JNK P Thr183,Tyr185 | MAPK Signaling, Stress pathway |
| JAK1 P Tyr1022,Thr1023 | Cytokine Signaling, Jak/Stat Signaling, Tyrosine kinases |
| Stat1 | Cytokine Signaling, Jak/Stat Signaling |
| STAT1 alpha P Y701 [EPR3147] | Cytokine Signaling, Jak/Stat Signaling |
| Stat1 P Ser727 | Cytokine Signaling, Jak/Stat Signaling |
| Stat3 | Cytokine Signaling, Jak/Stat Signaling |
| Stat3 P Ser727 | Cytokine Signaling, Jak/Stat Signaling |
| Stat3 P Y705 | Cytokine Signaling, Jak/Stat Signaling |
| Stat5 | Cytokine Signaling, Jak/Stat Signaling |
| Stat5 P Tyr694 | Cytokine Signaling, Jak/Stat Signaling |
| Stat6 | Cytokine Signaling, Jak/Stat Signaling |
| Stat6 P Tyr641 | Cytokine Signaling, Jak/Stat Signaling |
| Tyk2 P Tyr1054,Tyr1055 | Adaptors, Cytokine Signaling, Jak/Stat Signaling, Tyrosine Kinases |
| Smad1/5 P Ser463/Ser465 | cell growth, apoptosis, morphogenesis, development and immune responses |
| Smad2 (C86F7) | cell growth, apoptosis, morphogenesis, development and immune responses |
| Smad2 P Ser465,Ser467 | cell growth, apoptosis, morphogenesis, development and immune responses |
| Smad2/3 P Ser465/Ser423,Ser467/Ser425 | cell growth, apoptosis, morphogenesis, development and immune responses |
| Smad3 P Ser423,Ser425 | cell growth, apoptosis, morphogenesis, development and immune responses |
| TGF beta (56E4) | regulators of cell proliferation and differentiation, developmental patterning and morphogenesis, and disease pathogenesis |

**Extended Data Table S2 (continued from previous page):****Analytes targeted in RPPA**
|  |  |
| --- | --- |
| IkB-alpha | NFkB Signaling |
| IkB-alpha P Ser32 | NFkB Signaling |
| IKK alpha/beta P Ser176/Ser177 | lymphocyte proliferation, differentiation, growth, and survival |
| NFkB p105/p50 | inflammatory and immune responses |
| NFkB p65 Ser536 | inflammatory and immune responses |
| TBK1/NAK | NFkB regulation |
| Bad P Ser136 | Apoptosis |
| Bcl-2 | Apoptosis |
| Bcl-xl | Apoptosis |
| Bim | Apoptosis |
| Bim P Ser69 | Apoptosis |
| Caspase 3 | Apoptosis |
| Caspase 3 cleaved | Apoptosis |
| PARP | Apoptosis |
| PARP cleaved Asp214 | Apoptosis |
| Puma | Apoptosis |
| Bax | Apoptosis |
| Bcl-2 P Ser70 | Apoptosis |
| cdc25c P Ser216 | Cell Cycle Control |
| CDK1 (p34cdc2) P Tyr15 | Cell Cycle Control |
| Cyclin D1 | Cell Cycle Control |
| Cyclin D1 P Thr286 | Cell Cycle Control |
| p21 CIP/WAF1 | Cell Cycle Control |
| p21 CIP/WAF1 p Thr145 | Cell Cycle Control |
| Plk1 P | mitosis |
| Rb P Ser807,Ser811 | Apoptosis, Cell Cycle Control |
| Rb P Ser780 | Apoptosis, Cell Cycle Control |
| Chk1 P Ser345 | Cell Cycle Control |
| ATM | Cell Cycle Control, DNA Repair |
| ATM/ATR Substrate P Ser/Thr | Cell Cycle Control, DNA Repair |
| Ku80 (C48E7) | cell cycle regulation, DNA replication and repair, telomere maintenance, recombination, and transcriptional activation. |
| c-Fos Antibody (D-1) |  |
| Aurora A/B/C P Thr288/Thr232/Thr198 | Cell Cycle |
| c-Myc | MAPK Signaling, Transcription Factors |
| c-Myc P Thr58,Ser62 | MAPK Signaling, Transcription Factors |
| FOXO3a (75D8) | Apoptosis, Notch signalling |
| FRA1 (R20), (FOSL1) | Transcription Factors |
| p53 | Apoptosis, Cell Cycle Control |
| p53 P Ser15 | Apoptosis, Cell Cycle Control |
| E-Cadherin | Adhesion |
| PKC (pan) P Ser660 (beta-2) | Calcium, cAMP, Lipid Signaling, PKC Signaling |
| PKC substrate P (R/K)X(S*)(Hyd)(R/k) | Calcium, cAMP, Lipid Signaling, PKC Signaling |
| PLC-gamma1 P Tyr783 | Calcium, cAMP, Lipid Signaling |
| CREB | Calcium, cAMP, Lipid Signaling, PKC Signaling |
| Cleaved Notch 1 (val1744) | developmental processes via controlling cell fate decisions |
| GATA3 (D13C9) | Transcription factor, Notch signaling |
| beta-Catenin P Ser33,Ser37,Thr41 | Wnt Signaling |
| beta-Catenin P Thr41,Ser45 | Wnt Signaling |
| SQSTM1 (p62) | Autophagy |
| YAP P Ser127 | transcriptional co-activator, Hippo signalling |
| PKM2 P Tyr105 | aerobic glycolysis |
| Ki-67 (Annexin II, p36) | Tumor Marker, Proliferation, G0-negative Marker |
| beta-actin | Housekeeping, Cytoskeleton |
| beta-Tubulin | Housekeeping, Cytoskeleton |
| GAPDH | Housekeeping, glycolysis |

**Extended Data Table S3:** Published studies with protein abundances linked to patient outcomes.

| Cohort | Disease and treatments | MS data format | Covariates used |
| --- | --- | --- | --- |
| Dressler et al., 2024<sup>50</sup> | 242 tumour samples from 196 patients; NMIBC and MIBC; FFPE | TMT ratios relative to the pooled reference channel (linear scale) | Pathological stage: pTa low grade; pTa high grade or pTis; pT1a/b; pT1c; pT2; >pT2 or pN1. |
| Xu et al., 2022<sup>51</sup> | 116 treatment-naive patients; 45 NMIBC, 71 MIBC; FFPE | Label-free DDA, iBAQ-based fraction-of-total-abundances (linear scale) | Muscle invasion (MIBC vs NMIBC), age and sex. Grade was not used because none of the seven low-grade tumours had an event. |
| Qu et al., 2023<sup>52</sup> | 362 patients with upper-tract urothelial carcinoma; pre-operative plasma (discovery cohort) | DIA, iBAQ-based fraction-of-total-abundances (linear scale) | T stage (≥T2), nodal status (N+), grade (high vs low), age and sex. |

**Extended Data Table S4:** Proteins in each study that matched the entries of STV_imres_.

| Cohort | Entries matched (of 1,199) | Fraction of squared weight |
| --- | --- | --- |
| Dressler et.al. 2024 | 983 | 0.79 |
| Xu et.al. 2022 | 1,015 | 0.87 |
| Qu et.al. 2023 | 158 | 0.17 |

**Extended Data Table S5:**
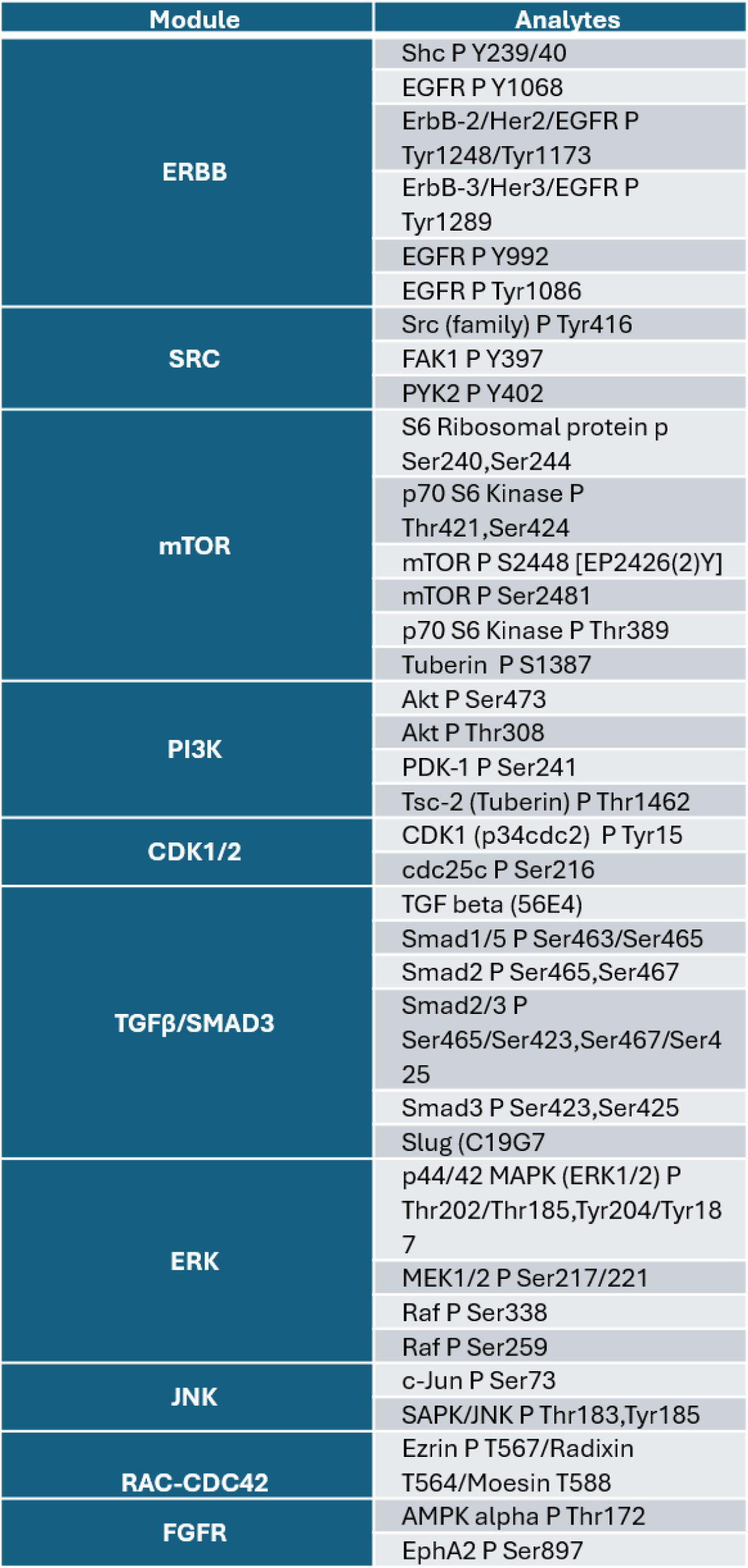
RPPA analyte outputs of each module.

**Extended Data Table S6:** Primary antibodies for Western blot.

| Antibody | Source | Catalog # |
| --- | --- | --- |
| Phosphorylated STAT1 (Ser727) | Cell Signaling Technologies | 8826 |
| Phosphorylated STAT3 (Tyr705) | Cell Signaling Technologies | 9145 |
| Xct/Slc7a11 | Cell Signaling Technologies | 12691 |
| NCOA4 | Cell Signaling Technologies | 66849 |
| GPX4 | Cell Signaling Technologies | 59735 |
| NINJ1 | Cell Signaling Technologies | 49761 |
| Phosphorylated MLKL (Ser345) | Cell Signaling Technologies | 37333 |
| Phosphorylated SQSTM1/p62 (Thr269/Ser272) | Cell Signaling Technologies | 13121 |
| CD45 | Invitrogen | MA537809 |
| GAPDH | Cell Signaling Technologies | 2118 |

**Extended Data Table S7:**
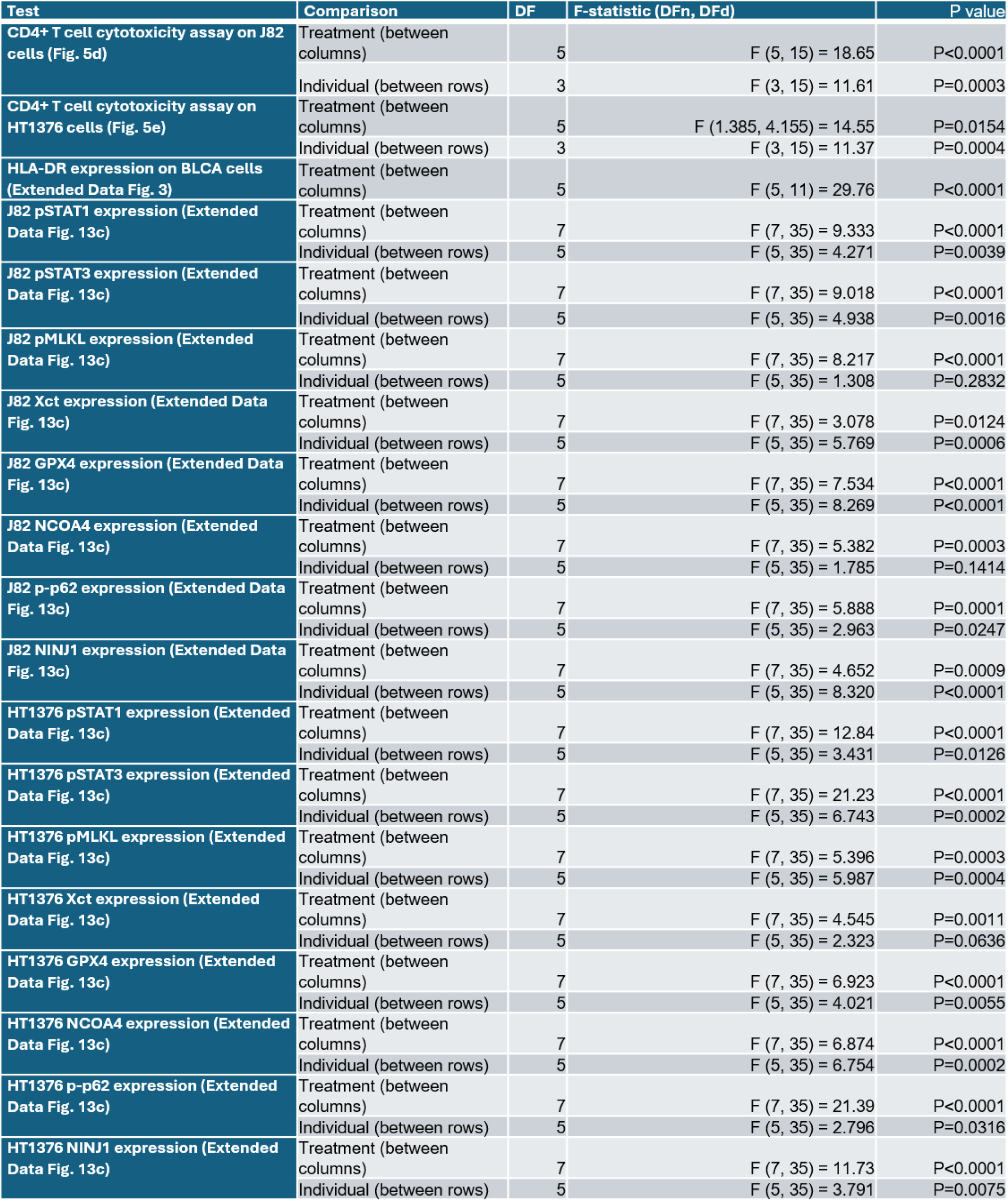
Statistics for ANOVA.

**Supplementary Figure 1.**
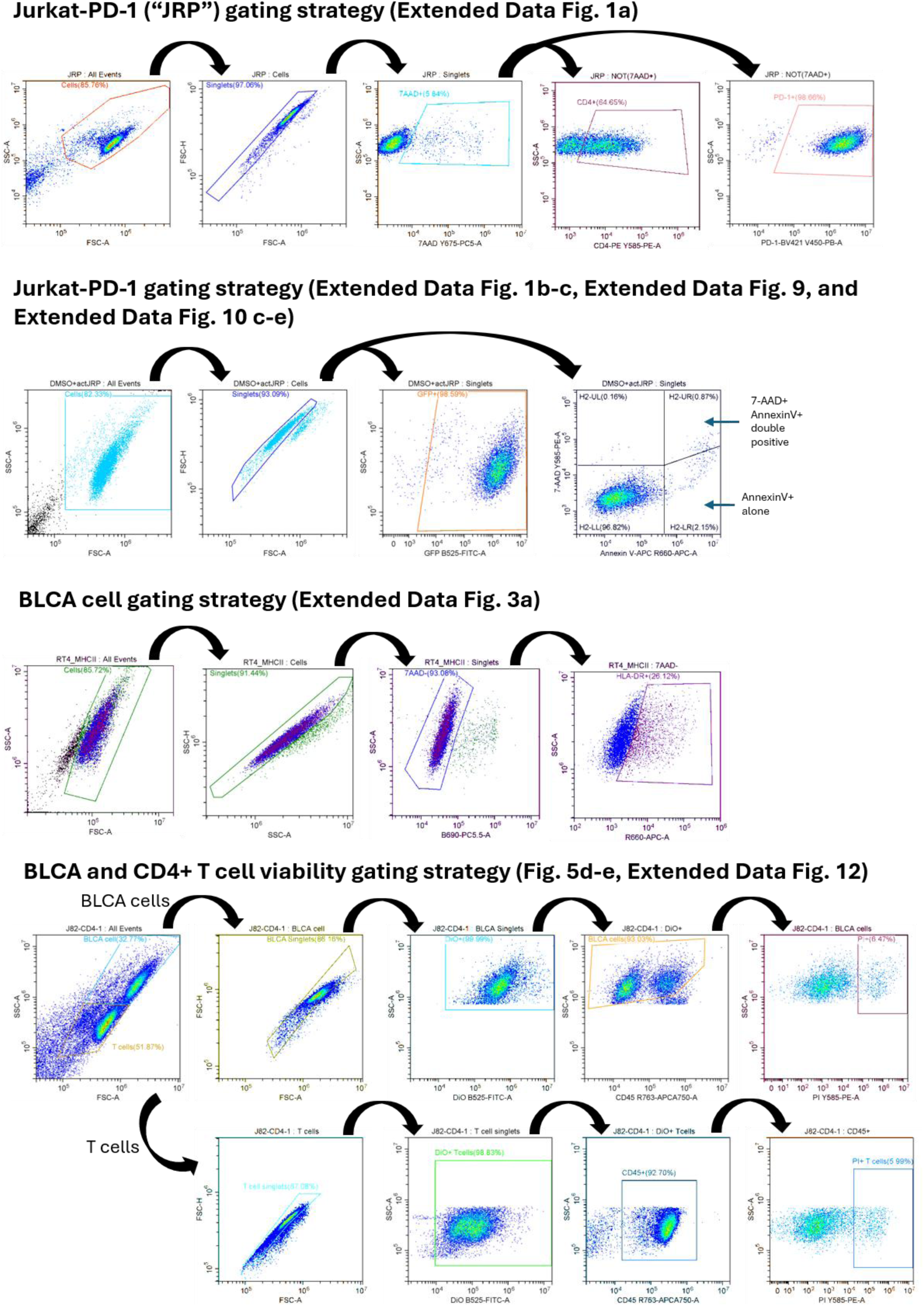
Flow cytometry analysis gating strategy for all experiments and related figures.

